# Phylogenomic Analyses Reveal Widespread Gene Flow During the Early Radiation of Oaks and Relatives (Fagaceae: Quercoideae)

**DOI:** 10.1101/2023.04.25.538215

**Authors:** Shuiyin Liu, Yingying Yang, Qin Tian, Zhiyun Yang, Shufeng Li, Paul J. Valdes, Alex Farnsworth, Heather R. Kates, Carolina M. Siniscalchi, Robert P. Guralnick, Douglas E. Soltis, Pamela S. Soltis, Gregory W. Stull, Ryan A. Folk, Tingshuang Yi

## Abstract

Oaks (*Quercus*), one of the most species-rich and ecologically dominant woody plant clades in the Northern Hemisphere, are well known for their propensity to hybridize and form syngameons, complexes where alleles are readily exchanged among closely related species. While hybridization has been extensively studied towards the tips of the oak phylogeny, the extent, timeline, and evolutionary scenarios of hybridization during the early radiation of oaks and related genera (Quercoideae) remain poorly known. Using an expansive new dataset of nuclear and chloroplast sequences (including up to 431 spp.), we conducted a multifaceted phylogenomic investigation of *Quercus* aimed at characterizing gene-tree and cytonuclear (chloroplast-nuclear) discordance and identifying ancient reticulation in the early evolution of the group. We document extensive nuclear gene-tree and cytonuclear discordance at deep nodes in *Quercus* and Quercoideae, with *Quercus* recovered as non-monophyletic in the chloroplast phylogeny. Analyses recovered clear signatures of gene flow against a backdrop of incomplete lineage sorting, with gene flow most prevalent among major lineages of *Quercus* and Quercoideae during their initial radiation, dated to the early-middle Eocene. Ancestral reconstructions including fossil data suggest that the ancestors of *Castanea+Castanopsis*, *Lithocarpus*, and the Old World oak clade co-occurred in North America and Eurasia, while the ancestors of *Chrysolepis, Notholithocarpus,* and the New World oak clade co-occurred in North America, offering ample opportunity for hybridization in each region. Following this initial phase of radiation and reticulation, we detected multiple niche shifts in *Quercus* and other Quercoideae genera that likely facilitated their expansion into new habitats arising from post-Eocene climatic changes. Our study shows that hybridization—perhaps in the form of ancient syngameons similar to those seen today—has been a common and important process throughout the evolutionary history of oaks and their close relatives.

---

Oaks (*Quercus*) are a diverse clade (ca. 435 spp.) of woody plants with vast ecological and economic importance and are well known for their propensity to hybridize across various phylogenetic and spatial scales (Hardin 1975; Whittemore and Schaal 1991; Kim et al. 2018; Crowl et al. 2020). Gene flow among oaks, although often taxonomically confounding, has also been highlighted as an evolutionary asset, allowing for the exchange of adaptive alleles across species boundaries (Petit et al. 2003; Dodd and Afzal-Rafii 2007; Leroy et al. 2020a; Nagamitsu et al. 2020). In some cases, sympatric or partially sympatric oak species form syngameons, where they are able to maintain relatively clear species boundaries, in a morphological sense, despite active gene flow (Burger 1975; Cannon and Petit 2020; Leroy et al. 2020b). While numerous studies have investigated gene flow toward the tips of oak phylogeny (e.g., Eaton et al. 2015; Hauser et al. 2017), less attention has been paid to the extent and evolutionary consequences of deep/ancient hybridization in *Quercus*.

Hybridization among closely related oak species (within groups of species recognized taxonomically as sections) is well documented (e.g., Curtu et al. 2007; Moran et al. 2012; Eaton et al. 2015; Feng et al. 2016; An et al. 2017); introgression between species from different sections has also been reported (Simeone et al. 2016; McVay et al. 2017; Crowl et al. 2020; Zhou et al. 2022). Less well understood is the extent of reticulation during the early evolution of *Quercus* and relatives within the broader context of the Fagaceae phylogeny. While the broad-scale nuclear phylogeny of *Quercus* appears well-resolved and congruent with morphology and the fossil record, high levels of gene-tree discordance have been documented along the backbone, possibly stemming from a combination of ancient gene flow (Crowl et al. 2020; Hipp et al. 2020) and incomplete lineage sorting (ILS; Zhou et al. 2022). Ancient gene flow is further supported by extensive levels of deep discordance between chloroplast and nuclear phylogenies (i.e., cytonuclear discordance): in contrast to nuclear phylogenies which show *Quercus* as monophyletic, chloroplast phylogenies typically show *Quercus* as non-monophyletic, with oak sections showing various relationships with other genera of Fagaceae subfamily Quercoideae (*Notholithocarpus*, *Chrysolepis*, *Lithocarpus*, *Castanopsis, Castanea,* and *Trigonobalanus*) (e.g., Manos et al. 2008; Xiang et al. 2014; Simeone et al. 2016; Yang et al. 2021; Zhou et al. 2022).

Despite accumulating evidence for ancient gene flow and reticulation in *Quercus*, limited or incompatible sampling of extant species for both nuclear and chloroplast data has made it difficult to pinpoint the phylogenetic locations of ancient reticulation. Furthermore, no synthetic examinations of the fossil record have been conducted to clarify the timing and geographic contexts of putative ancient reticulation events. Consequently, an integrative phylogenomic and paleobotanical study is needed to reconstruct detailed scenarios of ancient hybridization during the early radiation of this important clade, which has dominated temperate and subtropical forests in the Northern Hemisphere since the later Cenozoic (Barrón et al. 2017). Concomitantly, a “total evidence” approach, as applied here, establishes a framework useful not only for understanding the timing and location of deep reticulation in oaks, but also for addressing questions about the conditions in Earth history that may have promoted reticulation across the tree of life more generally.

Here we examine a newly constructed phylogenomic dataset, generated using both transcriptome and target-enrichment sequencing and including extensive nuclear and chloroplast loci across over 400 species, to infer phylogenetic relationships and instances of ancient gene flow in the early evolution of *Quercus*. Distinguishing gene flow from ILS is a challenging task, in part because no available methods can infer species trees while accommodating or parsing multiple sources of conflict (Morales-Briones et al. 2021b). Some of the most powerful discriminatory approaches, e.g. network methods, are currently unable to adequately scale given their heavy computational cost (e.g., Solís-Lemus et al. 2017; Wen et al. 2018). We therefore leverage a variety of currently available methods to dissect conflict and detect gene flow in early oak evolution—in concert with dating analyses and ancestral area and niche reconstructions. In particular, we integrate paleogeographic and paleoclimatic information from the extensive fossil record of Quercoideae to better place reticulation events in a geographic, geologic, and ecological context. In sum, we aim to (1) disentangle deep discordance among nuclear and chloroplast phylogenies of *Quercus* and relatives (Quercoideae); (2) identify the temporal window and phylogenetic locations of ancient reticulation events among sections of *Quercus*, and between major *Quercus* lineages and other genera of Quercoideae; and (3) assess whether the ancestors of *Quercus* lineages and other genera of Quercoideae overlapped in geographic and environmental space, thus providing opportunities for gene flow among these lineages consistent with the molecular evidence.

## Materials and Methods

### Study System and Taxon Sampling

We sampled 423 species of Fagaceae representing 314 species of *Quercus* (covering ∼72% of oak diversity and ≥ 58% of the species from each of the eight currently recognized sections: *Lobatae*, *Protobalanus*, *Ponticae*, *Virentes*, and *Quercus* of subgenus *Quercus*, and *Cyclobalanopsis*, *Ilex*, and *Cerris* of subgenus *Cerris* [Denk et al. 2017 and Hipp et al. 2020]) and 109 species of oak relatives (≥ 16% of the species from each of the remaining genera of Fagaceae; see Supplementary Table S1 for details). This includes *Fagus* (Fagoideae) as well as the six other genera of Quercoideae (beyond *Quercus*): *Notholithocarpus*, *Chrysolepis*, *Lithocarpus*, *Castanopsis*, *Castanea*, *Trigonobalanus*. Of the Fagaceae samples, 391 were newly sequenced using hybrid enrichment (hereafter, Hyb-Seq); the remaining 81 samples were obtained from the transcriptome dataset of Yang et al. (2021) and GenBank (Supplementary Table S2; accessed 1 May 2021). For outgroups, we included 15 species (seven were newly sequenced Hyb-Seq samples and eight were available transcriptomes) representing the major lineages of Fagales (i.e., Betulaceae, Casuarinaceae, Juglandaceae, Myricaceae, Nothofagaceae, and Ticodendraceae).

### Library Preparation and Sequencing

For hybrid enrichment, we used a set of exonic baits for 100 housekeeping genes designed for use across the rosid clade of angiosperms, but with a focus on the nitrogen-fixing clade, which includes Fagaceae. This locus panel and bait set, the “NitFix loci” reported in Folk et al. (2021), Fu et al. (2022) and Kates et al. (2022), were developed using MarkerMiner (Chamala et al. 2015) as universal across the rosids based on 78 phylogenetically representative transcriptomes from the 1KP project (Matasci et al. 2014; Leebens-Mack et al. 2019). Beyond the default filtering criteria implemented in MarkerMiner, which account for length, copy number, and other criteria (see Chamala et al. 2015), loci in the panel had at least 50% species coverage of the included rosid species. Loci were further manually curated to remove those with introns that were long (i.e., > 500 bp) or numerous, and exemplar alignments were trimmed to exclude positions present for a minority of taxa, suspected microsatellite regions, and loci from the organellar genomes. Isolated DNAs were submitted to Rapid Genomics (Gainesville, FL, USA) for quantification, library preparation, hybrid enrichment with the NitFix loci, and multiplex Illumina sequencing. Additional details of the sampling-to-sequencing workflow are given in the Supplementary Methods.

### Read Processing, Assembly, and Orthology Inference

For the hybrid enrichment dataset, raw reads were cleaned using Trimmomatic v.0.36 (Bolger et al. 2014) and assembled using HybPiper (Johnson et al. 2016). Orthology inference was carried out using several phylogenetic methods (i.e., the “monophyletic outgroup” and “rooted ingroup” approaches) from the pipelines of Yang and Smith (2014) and Morales-Briones et al. (2021a). For the transcriptome dataset, we used the pipeline of Yang and Smith (2014) for read processing, assembly, transcript processing, and homology and orthology inference. Further details on assembly, gene cluster processing, homolog tree inference, tip trimming, orthology inference, and ortholog cleaning are provided in the Supplementary Methods; an overview of how these Hyb-Seq and transcriptome datasets were used in downstream analyses is presented in Supplementary Figures S1 and S2 and Table S3. As described further in Supplementary methods, a single ortholog-filtered dataset was chosen for Hyb-Seq (“HYB-98RT” in Supplementary methods) and one for transcriptomics (“RNA-2821RT”) on the basis of relatively more loci and higher taxon coverage in each for examination of gene-tree discordance and gene flow.

### Plastome Assembly

We assembled plastomes from 391 Hyb-Seq and 60 transcriptome samples using a modified bash script (https://github.com/ryanafolk/Assembly-tools/). Forty-seven complete plastomes of Fagaceae and one of Betulaceae (used as an outgroup) were also downloaded from GenBank (accessed 5 April 2021). After filtering, the final dataset included 175 species of *Quercus* and 47 species from other genera of Fagaceae (222 plastomes of Fagaceae in total) as well as one plastome from outside Fagaceae (*Carpinus monbeigiana*, Betulaceae) used to root the tree (Supplementary Table S4). More details on plastome assembly and filtering are provided in the Supplementary Methods. An overview of the plastome dataset used in downstream analyses is provided in Supplementary Figure S3 and Table S3.

### Phylogenetic Analyses

We generated phylogenetic trees using maximum likelihood (ML) and coalescent-based methods. Concatenated ML analyses of the nuclear data were conducted in RAxML v.8.2.11 (Stamatakis 2014) under an unpartitioned GTR-GAMMA model with bootstrap support (BS) estimated by 1000 fast bootstrap replicates; individual nuclear gene trees were also generated using this same approach. Preliminary ML analyses of the concatenated dataset using 100 fast bootstrap replicates showed similar topologies under an unpartitioned GTR-GAMMA model and a gene-partitioned GTR-GAMMA model (results of which are provided in Supplementary Materials). Coalescent-based species tree inference was conducted using the nuclear gene trees in ASTRAL-III v.5.6.3 (Zhang et al. 2018) and support values were estimated with local posterior probabilities (LPP; Sayyari and Mirarab 2016). The chloroplast phylogeny was reconstructed through concatenated ML analyses (including 1000 bootstrap replicates) in two ways: with an unpartitioned GTR-GAMMA model and a GTR-GAMMA model partitioned by the large single-copy region, small single-copy region, and inverted repeat.

### Dissecting Discordance and Detecting Gene Flow

As a first step, we identified five deep nodes—concerning the phylogenetic positions of *Castanea* + *Castanopsis*, *Lithocarpus*, *Chrysolepis*, *Notholithocarpus*, and *Quercus* sect. *Cyclobalanopsis* (Supplementary Table S5)—showing elevated levels of conflict among nuclear genes and between nuclear and chloroplast datasets based on our results as well as those of previous studies (e.g., Manos et al. 1999; Hubert et al. 2014; Yang et al. 2021). Before investigating hybridization, we ruled out whether the data lack sufficient resolution by performing a polytomy test (Sayyari and Mirarab 2018), asking whether hard polytomies could be rejected at these uncertain nodes. We also used approximately unbiased (AU) tests (Shimodaira 2002) to examine whether particular tree topologies could be significantly rejected by the data. The phylogenetic signal of each topology was also quantified using the pipeline of Shen et al. (2017). Further details on our analyses of phylogenetic relationships, genomic conflict, and phylogenetic signal are provided in the Supplementary Methods. Lastly, we performed phyparts (Smith et al. 2015) and Quartet Sampling (QS; Pease et al. 2018) analyses to explore the overall phylogenetic patterns of gene tree discordance and to distinguish weak from strong support for conflict on a per-branch basis.

To investigate hybridization more directly, we first applied coalescent simulations to generate gene tree predictions under the inferred species tree and investigate whether the instances of deep cytonuclear discordance that we observed were due to ancient gene flow or ILS, following several studies (e.g., Folk et al. 2017; Morales-Briones et al. 2018; Stull et al. 2020). We further used the newly developed pipeline of Cai et al. (2021) to estimate the relative contributions of ILS, gene flow, and gene tree estimation error to the observed gene tree discordance. Additional details of coalescent simulations and implementation of Cai’s pipeline are provided in Supplementary Methods.

To further investigate lineages potentially involved in deeper-time instances of introgression (as well as the timing and directionality of such introgression), we first estimated a dated phylogeny using BEAST v.2.6 (Bouckaert et al. 2014) based on 20 nuclear genes selected using SortaDate (Smith et al. 2018) and ten well-vetted and previously used fossils (see Supplementary Table S6 for detailed prior settings). We also performed dating analyses of the chloroplast and nuclear trees to facilitate a comparison of the ages of deep divergences in these trees using treePL (Smith and O’Meara 2012). We used two calibration strategies (i.e., (1) two calibrations: crown group [CG] of Fagaceae, and CG of *Fagus*; and (2) four calibrations: CG of Fagaceae, CG of *Fagus*, stem group [SG] of the East Asian clade in sect. *Cerris*, and SG of sect. *Lobatae*) for these analyses, given that alternative relationships in the chloroplast tree precluded the use of several fossils used in the main nuclear dating analysis (see Supplementary Methods for details). We then applied the five-taxon *D*-statistic (*D*_FOIL_; Pease and Hahn 2015) to oaks and their relatives (*Notholithocarpus*, *Chrysolepis*, *Lithocarpus*, *Castanopsis*, and *Castanea*) using Ex*D*_FOIL_ (Lambert et al. 2019). On the basis of site-count patterns, this method can detect (with extremely low false-positive rates) the taxa involved in instances of introgression, as well as, in some cases, the directionality of gene flow (Pease and Hahn 2015). More details of Ex*D*_FOIL_ analysis are provided in Supplementary Methods.

We also used phylogenetic network analyses conducted using PhyloNet v.3.8.2 (Than et al. 2008; Wen et al. 2018) to explore whether reticulate evolution has occurred at deep locations in the Quercoideae phylogeny. Because the PhyloNet approach is computationally intensive, it was not possible to conduct network analyses with our complete species sampling. Three strategies were therefore used to generate three separate reduced-representation datasets (hereafter planA, planB, and planC; see Supplementary Methods for a full description of each). For each dataset, we performed model selection for the optimal network using AICc (Sugiura 1978) and BIC (Schwarz 1978). Further details on selecting representatives, network analyses, and model selection are provided in the Supplementary Methods.

### Ancestral Range and Niche Estimations

Ancestral range and niche estimations were conducted to assess the potential geographic and ecological overlap in deep time of lineages potentially involved in ancient hybridization events. We first delimited six biogeographic regions based on extant distributions of Fagaceae species (Manos and Stanford 2001): East Asia, South and Southeast Asia, Central and Western Asia, Europe (including a portion of North Africa), temperate and subtropical North America, and tropical America. The estimates of ancestral biogeographic range were conducted in the R package BioGeoBEARS (Matzke 2013) under the DEC, DIVALIKE, and BAYAREALIKE models, using the maximum clade credibility tree from the BEAST analysis of Hyb-Seq dataset as the input tree. The maximum ancestral range size was constrained to three. The three biogeographic models were compared using AIC to select the optimal one.

For ancestral niche estimations, we first collected occurrence records for each sampled species from GBIF (https://www.gbif.org/), iDigBio (https://www.idigbio.org/), NSII (http://nsii.org.cn/2017/home-en.php), and the literature (last accessed 16 November 2021), and 187,117 occurrence records were retained after filtering. We assembled 35 environmental layers at 30-seconds resolution. After the removal of highly correlated variables (Pearson’s *r* ≥ 0.7), we were left with 12 representative variables covering aspects of climate, topography, soil, and land cover (see the Supplementary Methods for details). To integrate the statistical distribution of niche space occupancy of all species, predicted niche occupancy profiles (PNOs, Evans et al. 2009) were applied to ancestral niche estimation using BayesTraits v.2.0 (http://www.evolution.rdg.ac.uk/BayesTraitsV2.html), implemented using the python wrapper ‘ambitus’ (Folk et al. 2018b). Briefly, this approach proportionally samples 100 samples for environmental value from a PNO profile for each present-day species and variable, and subsequently each sample is used as observed value of each species in a single ancestral niche estimation analysis (leading to 1200 estimation analyses); ancestral niche predictions are constructed from pooled posterior samples. Each analysis was run on the best ML tree and 1000 RAxML fast bootstrap trees from Hyb-Seq dataset as a means of accounting for phylogenetic uncertainty. Further details on occurrence cleaning, variable reduction, and the parameters used in ancestral niche estimation are provided in the Supplementary Methods.

We also included paleogeographic information from the fossil record of oaks and their relatives, as a complement to the geographic and niche reconstructions using extant species, with the general goal reconstructing geographic and ecological scenarios of ancient hybridization during the early evolution of Quercoideae. We used fossil-based ecological niche modeling (PaleoENM; Meseguer et al. 2015; Myers et al. 2015) to project the potential distribution of oaks and relatives in the past—instead of biogeography and niche reconstructions with an extant-extinct tree (e.g., Meseguer et al. 2018; Landis et al. 2021; Zhang et al. 2022). This was done instead of a total-evidence phylogenetic approach (involving the placement of fossils in the tree through combined molecular-morphological analyses), given the challenge accurately placing fossils with severely limited numbers of morphological characters (Grímsson et al. 2015, 2016; Barrón et al. 2017; Liu et al. 2020). The PaleoENM approach was performed as follows: we compiled comprehensive Paleogene fossil distribution data for oaks and relatives from the literature and online resources (CAD, Xing et al. 2016; PBDB, https://paleobiodb.org) (last accessed 21 November 2022); 468 fossil occurrences were retained after filtering (Supplementary Materials). We then extracted paleo-climate values using the paleo-coordinates of each fossil from seven paleo-climate models for seven periods (Early Paleocene, Late Paleocene, Early Eocene, Middle Eocene, Late Eocene, Early Oligocene, and Late Oligocene; Valdes et al. 2017; 2021). The paleo-coordinates were converted from modern coordinates using the “reconstruct” function from the R package “chronosphere” (Kocsis and Raja 2020). Each paleo-climate model includes eight temperature and precipitation layers at 5-minutes resolution. These climate values were used for multiple comparisons of niche among oaks and relatives during the Paleogene, and estimations of past climate tolerances with two strategies (i.e., using fossils from the Paleocene and Eocene, and using fossils from the Paleocene, Eocene and Oligocene) that subsequently were projected into a suite of paleo-climate models (i.e., environmental suitability rules are projected geographically through time). The mahalanobis distance was calculated to represent the environmental suitability in each grid cell. More details on the processing of fossil data, the paleo-climatic variables, and the PaleoENM analysis are provided in the Supplementary Methods.

## Results and Discussion

### New Insights into the Phylogeny of Oaks and Relatives

We generally recovered consistent backbone relationships across analyses (ASTRAL and RAxML) of the nuclear datasets (two Hyb-Seq and two transcriptome datasets), supporting the monophyly of *Quercus* (BS = 48–100; LPP = 0.49–1) and phylogenetic structure consistent with the taxonomic division of *Quercus* into two subgenera and eight sections (Fig. 1; Supplementary Fig. S4–S11; see Supplementary Results for more details). These findings are consistent with other recent studies (e.g., Denk et al. 2017; Hipp et al. 2020; Zhou et al. 2022). The placement of sect. *Cyclobalanopsis*, which is characterized by concentric lamellae on the cupule, has long been controversial (Manos et al. 1999; Hubert et al. 2014; Deng et al. 2018). Our concatenated ML and ASTRAL trees (except for the ML tree of HYB-98RT) both placed sect. *Cyclobalanopsis* with sect. *Ilex* and sect. *Cerris* to form an Old World oak clade (i.e., subg. *Cerris*; BS = 99–100; LPP = 0.44–1). The quantification of phylogenetic signal demonstrated that this topology was supported by the highest proportion of genes and sites, yet two alternative topologies (i.e., sect. *Cyclobalanopsis* sister to the remaining oaks [T2] or sister to *Notholithocarpus* [T3]) also had strong gene-wise and site-wise phylogenetic signal (Supplementary Fig. S12e; S13e; see Supplementary Results). Topology T3 of sect. *Cyclobalanopsis*, with 7%–10% clade probabilities across simulated gene trees (Supplementary Fig. S14–S15), was partially within ILS predictions, while gene flow between the clade of sect. *Ilex* + sect. *Cerris* and the American oak clade (i.e., subg. *Quercus*), as inferred by species networks (Supplementary Fig. S16e; 16f) and reported previously (Schnitzler et al. 2004; Mir et al. 2006; Zhou et al. 2022), might explain T2 for sect. *Cyclobalanopsis*. Thus, we argue this conflict may represent not only ILS but also past introgression in the early history of oak diversification.

**FIGURE 1.**
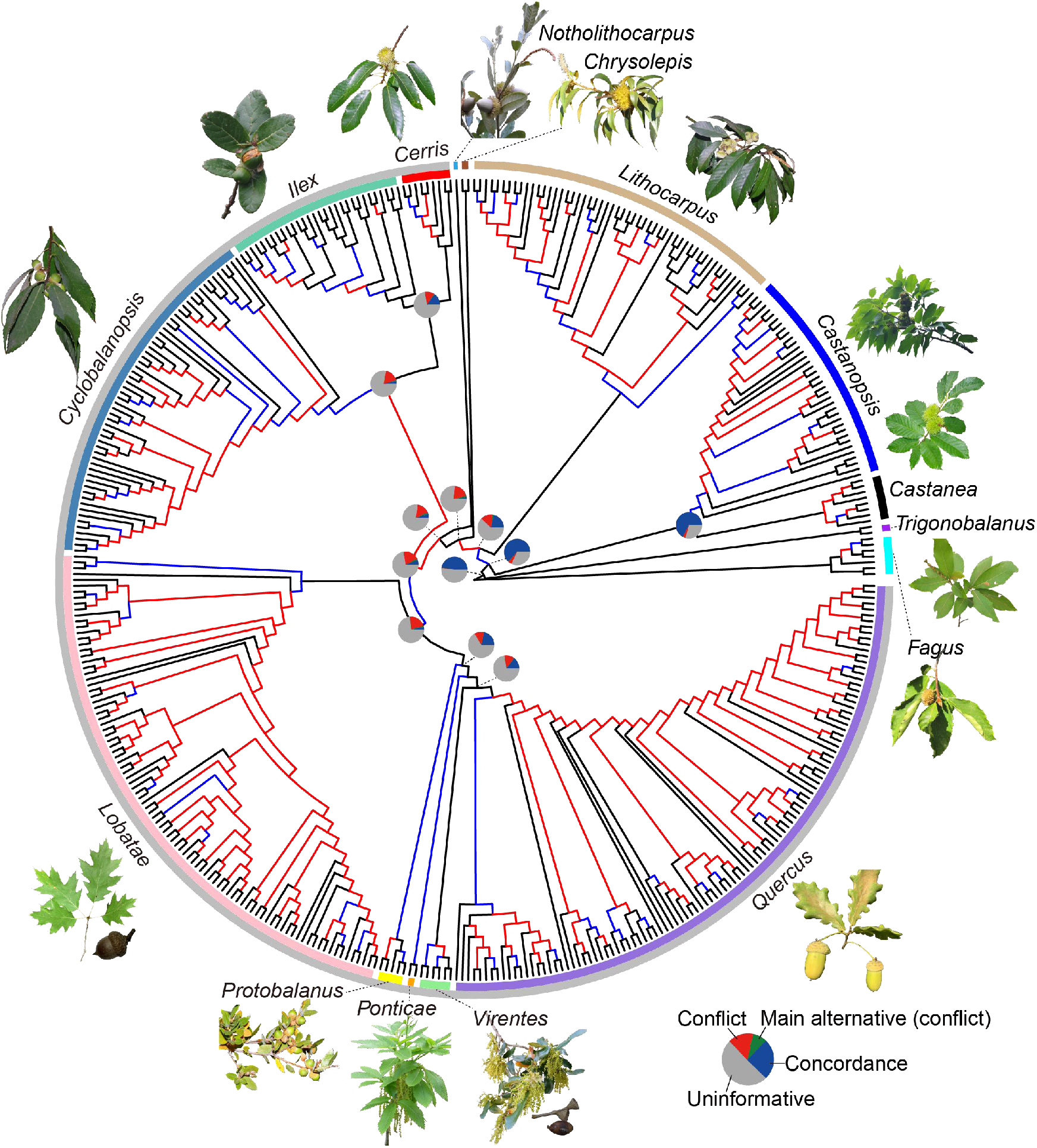
Cladogram of the species tree of oaks and relatives inferred by ASTRAL-III based on the HYB-98RT dataset including 431 species and 98 loci. Tip labels are shown in Supplementary Figure S5. Branches showing consistent relationships between ASTRAL-III, RAxML, and the four nuclear datasets (HYB-98RT, HYB-89MO, RNA-2821RT, and RNA-977MO) are colored with black (LPP ≥ 0.95 or BS ≥ 95% in all eight species trees) and with blue (LPP < 0.95 or BS < 95% in any one of the eight trees). Branches showing conflicting relationships among the eight trees are colored with red. Pie charts for major clades show the percentage of 2821 nuclear gene trees (RNA-2821RT) that support that clade (blue), a main alternative topology (green), all of the remaining alternatives (red), or are uninformative (i.e., < 70% BS or inadequate taxon sampling; gray) (see Supplementary Fig. S18-S19 for phyparts results of full-taxa sets). Fagaceae genera (external circle) and oak sections (internal circle) are indicated by colored bars, and their pictured representatives are: *Fagus grandifolia* by Bruce Kirchoff, *Chrysolepis chrysophylla* by J. Maughn, *Notholithocarpus densiflorus* by theforestprimeval, *Q. chrysolepis* by copepodo, and *Q. pontica* by peganum from https://search.creativecommons.org/; *Trigonobalanus doichangensis* by Li Chen (with permission); *Q. rubra* and *Q. robur* by Yingying Yang; others by Shuiyin Liu.

Regarding relationships among subclades within oak sections, we recovered some well-supported relationships that differed from previous studies. For instance, in sect. *Ilex*, a well-resolved grade (BS = 74–100; LPP = 0.53–0.99) of six subclades was documented comprising two East Asian groups (BS = 100; LPP = 0.96–1), a Himalaya-Mediterranean group (sclerophyllous; BS = 100; LPP = 0.92–0.96), two southwest-China groups (BS = 100; LPP = 0.98–0.99), and a Himalayan subalpine group (sclerophyllous; BS = 100; LPP = 0.99–1) (Supplementary Fig. S4–S7); a sister relationship between the two sclerophyllous groups (as found by Hipp et al. 2020) was not recovered, which was consistent with Jiang et al. (2019). In sect. *Cyclobalanopsis*, we recovered *Q. gaharuensis*, a species inhabiting Borneo, Sumatra, and Malaysia that had not been previously included in any phylogenetic studies, as sister to the rest of the section with full or high support (BS = 100; LPP = 0.92; Supplementary Fig. S4–S7). The CTB group, characterized by having compound trichome bases (Deng et al. 2014; Deng et al. 2018), was previously believed to be sister to the rest of the section (Deng et al. 2018; Hipp et al. 2020). This section was previously inferred to have originated in the Paleotropics (Deng et al. 2018), and the placement of *Q. gaharuensis* sister to the rest of the section, as found here, corroborates this previous finding. In other sections (e.g., sect. *Lobatae* and sect. *Quercus*), there is considerable conflict concerning phylogenetic relationships (Supplementary Figs. S4–S11) likely due to ILS (resulting from rapid radiations) as well as active and/or ancient gene flow among the constituent species (Hipp 2015; Cannon and Petit 2020; Kremer and Hipp 2020).

More broadly, our nuclear analyses support the monophyly of the eight genera in Fagaceae (i.e., *Quercus*, *Notholithocarpus*, *Chrysolepis*, *Lithocarpus*, *Castanopsis*, *Castanea*, *Trigonobalanus*, and *Fagus*) and provide new insights into their inter-relationships as a result of our more extensive sampling of both taxa and loci (Supplementary Fig. S4–S11) compared with previous studies (e.g., Manos and Stanford 2001; Manos et al. 2001, 2008; Lang et al. 2006; Xiang et al. 2014; Yang et al. 2018; Jiang et al. 2021; Zhou et al. 2022). In the Quercoid clade of Quercoideae (i.e., all genera excluding *Trigonobalanus*; referred to as the hypogeous seed clade in Zhou et al. 2022), we recovered a nuclear topology of *Castanea* + *Castanopsis* sister to the remaining Quercoids; this relationship showed high proportions of gene-wise and site-wise phylogenetic signal (Supplementary Fig. S12a; S13a) and relatively high numbers of concordant gene trees and quartets in the phyparts and QS results (Fig. 1; Supplementary Fig. S17–S19). Concerning the relationship between *Lithocarpus* and *Chrysolepis*, collapsing these branches (i.e., *Lithocarpus*, *Chrysolepis*, and *Notholithocarpus* + *Quercus*) into a polytomy was significantly rejected in the transcriptome dataset (Supplementary Table S5). Although most inferred species trees recovered *Chrysolepis* sister to the clade of *Notholithocarpus* + *Quercus* [T1], an alternative topology (i.e., *Chrysolepis* sister to *Lithocarpus* [T2], which was also recovered by Zhou et al. [2022] using 2124 nuclear loci) had similar proportions of supporting genes and sites (potentially indicating the presence of ILS; Supplementary Fig. S13c) or even higher proportions (Supplementary Fig. S12c). Topology T2 for *Chrysolepis* showed 13%–17% clade probabilities in the simulated gene trees (Supplementary Fig. S14–S15), while T1 showed 23%–46% (Fig. 2a; Supplementary Fig. S20–S23), indicating that to some extent they both were within ILS predictions. However, ancient introgression between *Chrysolepis* and *Lithocarpus*, as inferred by *D*_FOIL_ tests, might be responsible for T2 of *Chrysolepis* (Supplementary Table S7). Similarly, ancient introgression between *Chrysolepis* and *Quercus* might explain T1 of *Chrysolepis* (Supplementary Fig. S24–S25 and Table S7). The likely combination of both ILS and ancient hybridization in this case makes it difficult, if not impossible, to confidently reconstruct a bifurcating species tree for *Chrysolepis* and its immediate relatives (*Lithocarpus*, and *Notholithocarpus* + *Quercus*). Future studies employing multifaceted analyses of the complete nuclear genome sequence data might provide clearer resolution of the placement of this genus as well as further insight on the nature of the observed phylogenomic conflict.

**FIGURE 2.**
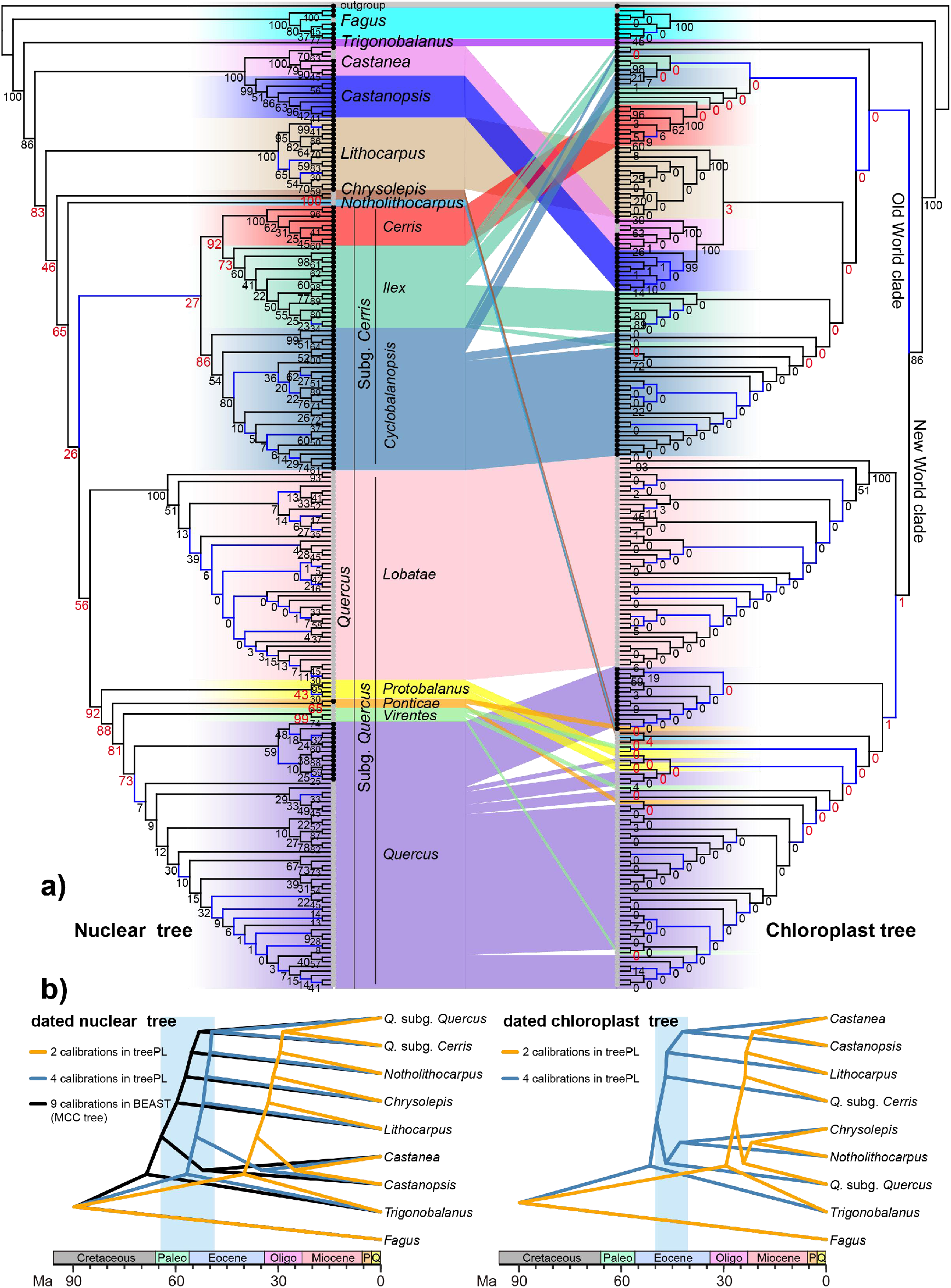
Discordance between dated nuclear and chloroplast phylogenies for oaks and relatives. **a)** Cophylogeny showing incongruence between the nuclear ASTRAL (left; HYB-98RT) and unpartitioned chloroplast ML (right) trees. Tip labels are shown in Supplementary Figure S20. Branches with low support values (LPP < 0.5; BS < 50%) are colored with blue. Clade frequencies of gene trees from the coalescent simulations are shown near the nodes; clade frequencies associated with deep cytonuclear discordances are red and enlarged. Black dots at the tips denote an Old World distribution, while gray dots denote a New World distribution. **b)** Time-calibrated nuclear and chloroplast trees showing a lag of the divergences of *Quercus* and the other main clades of Quercoideae in chloroplast tree compared to the nuclear tree. Only the divergence times of major lineages are shown here (see Supplementary Materials for the full dated trees). The blue window (ranging from the early Paleocene to middle Eocene) encompasses the time-frame of divergences of the main lineages in Quercoideae, based on analyses with four and nine calibrations. Abbreviations: Paleo, Paleocene; Oligo, Oligocene; P, Pliocene; Q, Quaternary.

### Dissecting Causes of Gene Tree Heterogeneity and Deep Cytonuclear Discordance

We found decisive support for a single topology in several important areas of the phylogeny (i.e., stem groups of *Fagus* and *Trigonobalanus*; and crown groups of *Castanea* + *Castanopsis* and the Quercoid clade of Quercoideae; Fig. 1; Supplementary Fig. S17). However, overall, we observed high levels of gene-tree discordance across Fagaceae phylogeny, particularly within the clade formed by *Lithocarpus*, *Chysolepis*, *Notholithocarpus*, and *Quercus*. Within this clade of four genera, conflict increases toward the terminals (Supplementary Fig. S18–S19). Results from co-quantification of ILS, gene flow, and estimation error showed that ILS (3.25%–36.31%) and gene flow (2.19%–13.02%) explained most of the observed gene tree discordance; gene tree estimation error accounted for a lower proportion (0.62%–3.57%) (Supplementary Fig. S26; see Supplementary Results for details). It is noteworthy that gene flow estimates differed between datasets (Supplementary Fig. S26), suggesting that taxon sampling might determine how much gene flow is observable in any particular dataset. Similarly, we found that the gene tree variation explained by the model of these three factors decreased as taxon sampling increased (see *R*^2^ in Supplementary Fig. S26), suggesting that observed gene tree discordance is influenced by taxon sampling.

High levels of cytonuclear discordance were observed at both deep and shallow levels within *Quercus* and across Quercoideae (Fig. 2a; Supplementary Fig. S20–S23), as has been shown by previous studies (e.g., Manos et al. 1999; Simeone et al. 2016; Pham et al. 2017; Yang et al. 2021; Zhou et al. 2022). However, our increased sampling compared to earlier efforts, with close taxonomic complementarity between the nuclear and chloroplast datasets, provided greater resolution of the patterns of cytonuclear incongruence across Quercoideae. In our chloroplast phylogeny, *Quercus* was recovered as non-monophyletic, with members of the genus forming two geographically distinct clades with the other genera of Quercoideae: a clade of (*Lithocarpus*, (*Castanea*, *Castanopsis*)) nested in the Old World oak clade (corresponding to the traditionally recognized subg. *Cerris*); and a clade with *Chrysolepis* + *Notholithocarpus* nested in the New World oak clade (corresponding to the traditionally recognized subg. *Quercus*) (Fig. 2a; Supplementary Fig. S27–S28; see Supplementary Results for details). Given that both gene flow and ILS can result in discordance between nuclear and chloroplast phylogenies (e.g., Rieseberg and Soltis 1991; Folk et al. 2017; Rose et al. 2021), we conducted various analyses to identify the causes of the observed instances of cytonuclear discordance. Our coalescent simulations demonstrated that the observed cytonuclear discordance was partially within ILS predictions at shallow levels, but at deeper levels we generally observed ∼0% frequencies of observed clades in the gene trees simulated under ILS (Fig. 2a; Supplementary Fig. S20–S23), which suggests that the chloroplast backbone topology is not within ILS expectations and supports gene flow as the source of the observed deep conflicts, in agreement with the findings of Zhou et al. (2022). We found that chloroplast topologies of *Castanea* + *Castanopsis*, *Lithocarpus*, *Chrysolepis*, and *Notholithocarpus* were supported by appreciable proportions of nuclear genes (15%–52%) and sites (9%–39%) (Supplementary Fig. S12a–d; S13a–d), suggesting that both chloroplast haplotypes and nuclear alleles were introgressed during historic instances of gene flow between *Quercus* and relatives. This scenario is further supported by the *D*_FOIL_ tests, which detected evidence of ancient introgression between some Eurasian genera—i.e., *Lithocarpus*, *Castanea,* and *Castanopsis*—and the Old World oak clade as well as between North American *Chrysolepis* and the New World oak clade (Supplementary Table S8); overall, suspected ancient introgression mostly involved distantly related lineages that geographically overlap. While introgression among these genera was not detected in a previous effort using ABBA-BABA tests (Zhou et al. 2022), this discrepancy is most likely due to the relatively low number of four-taxon tests conducted for low taxon sampling (5,316,699 tests in Hyb-Seq dataset and 224,998 tests in transcriptome dataset VS. 25, 882 tests in Zhou et al. 2022). Overall, our analyses consistently show that ancient gene flow, rather than ILS, is the primary source of deep cytonuclear discordance in Quercoideae.

Similar geographic patterns of ancient gene flow (including cytonuclear discordance) were also found between species of oaks from clades recognized as sections—for example, between *Q. pontica* (sect. *Ponticae*) and the Roburoid lineage (sect. *Quercus*), between species of sect. *Cyclobalanopsis* and some species of sect. *Ilex*, between sect. *Cerris* and some species of sect. *Ilex*, and between sect. *Protobalanus* and the Dumosae lineage (sect. *Quercus*) (all lineage names following [Hipp et al. 2020]). In these cases, the chloroplast topologies were not within ILS predictions (Fig. 2a; Supplementary Fig. S20–S23), suggesting instead the occurrence of historic asymmetrical gene flow between these lineages, leading to a local geographic structure in the chloroplast phylogenies at odds with the nuclear species-tree reconstruction. Furthermore, an abundance of nodes with moderate or weak BS support in chloroplast trees, particularly toward the tips (Supplementary Fig. S27–S28), suggests that plastomes have generally tracked geographic structure and have limited information on recent oak divergences, in agreement with previous studies (Yang et al. 2016; Pham et al. 2017). Collectively, these findings indicate that the plastome is not a reliable source of data for resolving phylogenetic relationships at any evolutionary scale in Quercoideae, although paired with the nuclear genome, the plastome is useful for shedding light on historic gene flow in the evolutionary history of oaks.

### Genomic, Geographic, and Climatic Evidence for Widespread Gene Flow During the Eocene in Two Geographic Centers

Widespread signals of ancient gene flow are inferred between oaks and all other genera in Quercoideae, as well as within oaks, as indicated by *D*_FOIL_ tests (Supplementary Fig. S24–S25), PhyloNet results (Supplementary Fig. S16), and patterns of cytonuclear discordance (Fig. 2a; Supplementary S20–S23; see Supplementary Results for details). This finding is consistent with other studies of ancient introgression between oaks and their relatives (e.g., Simeone et al. 2016; Yang et al. 2021; Zhou et al. 2022), among sections of oaks (e.g., Manos et al. 1999; Mir et al. 2006, 2009; McVay et al. 2017), and within oak sections (e.g., Sullivan et al. 2016; An et al. 2017; Ortego et al. 2018; Castillo-Mendoza et al. 2019).

Given the complex modern and historical distribution of oaks and relatives (Zhou et al. 1999; Grímsson et al. 2015, 2016; Barrón et al. 2017; Cannon et al. 2018; Liu et al. 2020), the question is raised as to whether the lineages suggested by molecular data to have undergone hybridization had ancestral distributions and ecologies consistent with that scenario. The comparison of divergence times from chloroplast and nuclear estimates showed that the divergences of *Quercus* and the other main clades of Quercoideae (*Notholithocarpus, Chrysolepis*, *Lithocarpus*, and *Castanopsis + Castanea*) in the chloroplast tree occurred in the middle Eocene, following the inferred divergence times from the nuclear tree (Paleocene to early Eocene, depending on the dating approach; Fig. 2b; see Supplementary Results for details), and younger than a recently inferred chloroplast dating result (early to middle Eocene; Zhou et al. 2022) due to the different strategy of calibrations. Ancestral range estimations using both extant species (under the DEC model; Fig. 3a; Supplementary Fig. S29) and the fossil record (PaleoENM analysis; Fig. 3b; Supplementary Fig. S30-S31) generally supported the possibility of historical overlap facilitating gene flow, but a different picture of the geographic dynamic was revealed by these two approaches. Biogeographic results based on extant species supported an East Asian origin for Quercoideae, with subsequent dispersals to North America, Southeast Asia, and Europe during the Paleocene-Eocene (Fig. 3a; Supplementary Fig. S29), a globally warm period with available land bridges and limited barriers in the Northern Hemisphere (Tiffney and Manchester 2001). The estimated ancestral distributions of the stem and crown groups of three Old World genera—*Castanea, Castanopsis*, and *Lithocarpus*—and the Old World oak clade all contained the East Asia region; the estimated ancestral distributions of the stem and crown groups of two New World genera—*Chrysolepis* and *Notholithocarpus—*and the New World oak clade all contained the North America region. However, the PaleoENM analysis, which reconstruct suitable habitat rather than the dispersal process, suggested that major lineages of Quercoideae were quite widespread across the Northern Hemisphere during the early Paleogene; four Old World lineages (*Castanea, Castanopsis*, *Lithocarpus*, and subg. *Cerris*) potentially co-occurred in North America and Eurasia during the Paleocene-Eocene; and three New World lineages (*Chrysolepis*, *Notholithocarpus*, and subg. *Quercus*) potentially co-occurred in North America (Fig. 3b; Supplementary Fig. S30-S31). The distribution of known fossils is similarly broad (Fig. 3b; Kvaček and Walther 1989; Zhou et al. 1999; Grímsson et al. 2015, 2016; Barrón et al. 2017). And both reconstructed potential distributions and fossil records of these lineages were found to have expanded and increased toward the Oligocene (particularly for temperate lineages like subg. *Quercus*; Fig. 3b; Supplementary Fig. S30), suggesting the rapid diversification of crown lineages in Quercoideae after the Eocene, consistent with previous findings (Hipp et al. 2020; Kremer and Hipp 2020; Zhou et al. 2022). The disagreement between the relatively narrow range reconstruction of biogeographic reconstruction and the fossil record might be explained by limitations of relying only on data on extant species. High extinction rates in the North Hemisphere, particularly North America, Europe, and Central Asia, since the Oligocene, reflect cooling and drying trends which would have removed many lineages in these regions, obscuring historical distributions.

**FIGURE 3.**
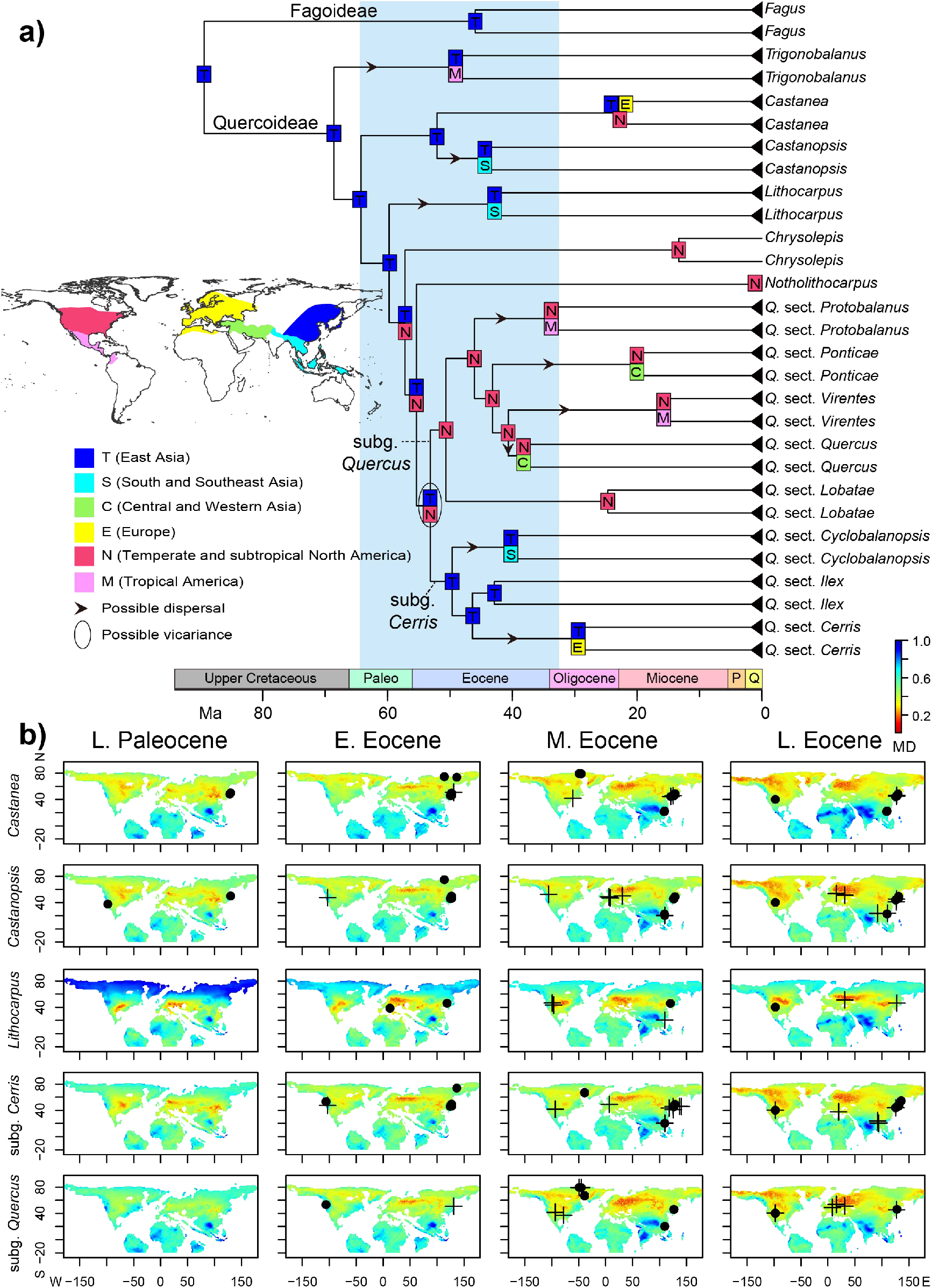
Overlapped historical geographic space between ancestors of *Quercus* lineages and other genera of Quercoideae involved in deep reticulation. **a)** Ancestral range estimation of Fagaceae under the DEC model with extant species. Only the reconstructed distributions for the stem and crown groups of eight genera in Fagaceae and eight sections in *Quercus* are shown here (see Supplementary Figure S29 for the full reconstruction). The blue rectangle encompasses the temporal window of divergences of the major lineages of *Quercus* and Quercoideae. Abbreviations: Paleo, Paleocene; P, Pliocene; Q, Quaternary. **b)** Potential distribution of the major lineages of *Quercus* and Quercoideae from Late Paleocene to Late Eocene as inferred by fossil-based ecological niche modeling. These maps were generated by projecting the climatic tolerances of Paleocene and Eocene fossil taxa onto four paleo-climate scenarios. Similar pattern of past distribution was obtained by projecting the climatic tolerances of Paleocene, Eocene, and Oligocene fossils onto six paleo-climate scenarios (Supplementary Fig. S30). The fossil distribution of *Chrysolepis* and *Notholithocarpus* during the Paleogene is provided in Supplementary Figure S31. A MD (mahalanobis distance) score of < 0.3 (red) corresponds with highly suitable region for each Quercoideae lineages; climatically unsuitable regions are colored green to blue. Dots and crosses denote the location of pollen and macrofossil taxa, respectively. Abbreviations: E., Early; M., Middle; L., Late.

Our ancestral reconstructions suggest that the lineages involved in ancient gene flow not only showed broad geographic overlap but also had similar climatic and ecological preferences during the Eocene; this finding was robust to the inclusion of fossil data (Fig. 4; Supplementary Fig. S32-33). In particular, the most recent common ancestor (MRCA) of the Old World oak clade (i.e., subg. *Cerris*) and the MRCAs of *Castanea* + *Castanopsis* and *Lithocarpus* were estimated to have co-occurred in warm and moist habitats characteristic of evergreen broadleaf forests (Fig. 4a). The MRCA of the New World oak clade (i.e., subg. *Quercus*) and the MRCA of *Chrysolepis* (or *Notholithocarpus*) were inferred to have co-occurred in habitats similar to those above yet with relatively less water and heat and higher hydrothermal variation (Fig. 4a; Supplementary Table S9). We note, however, that these reconstructed ancestral overlaps could be an artifact of each lineage encompassing considerable niche breadths today, with intermediate conditions generally inferred for the ancestors. Another challenge is that gene flow could serve to homogenize ecological niches among reticulating lineages, obscuring any differences present prior to hybridization. Nevertheless, these reconstructions are broadly consistent with the hybridization scenarios supported by other data types and analyses.

**FIGURE 4.**
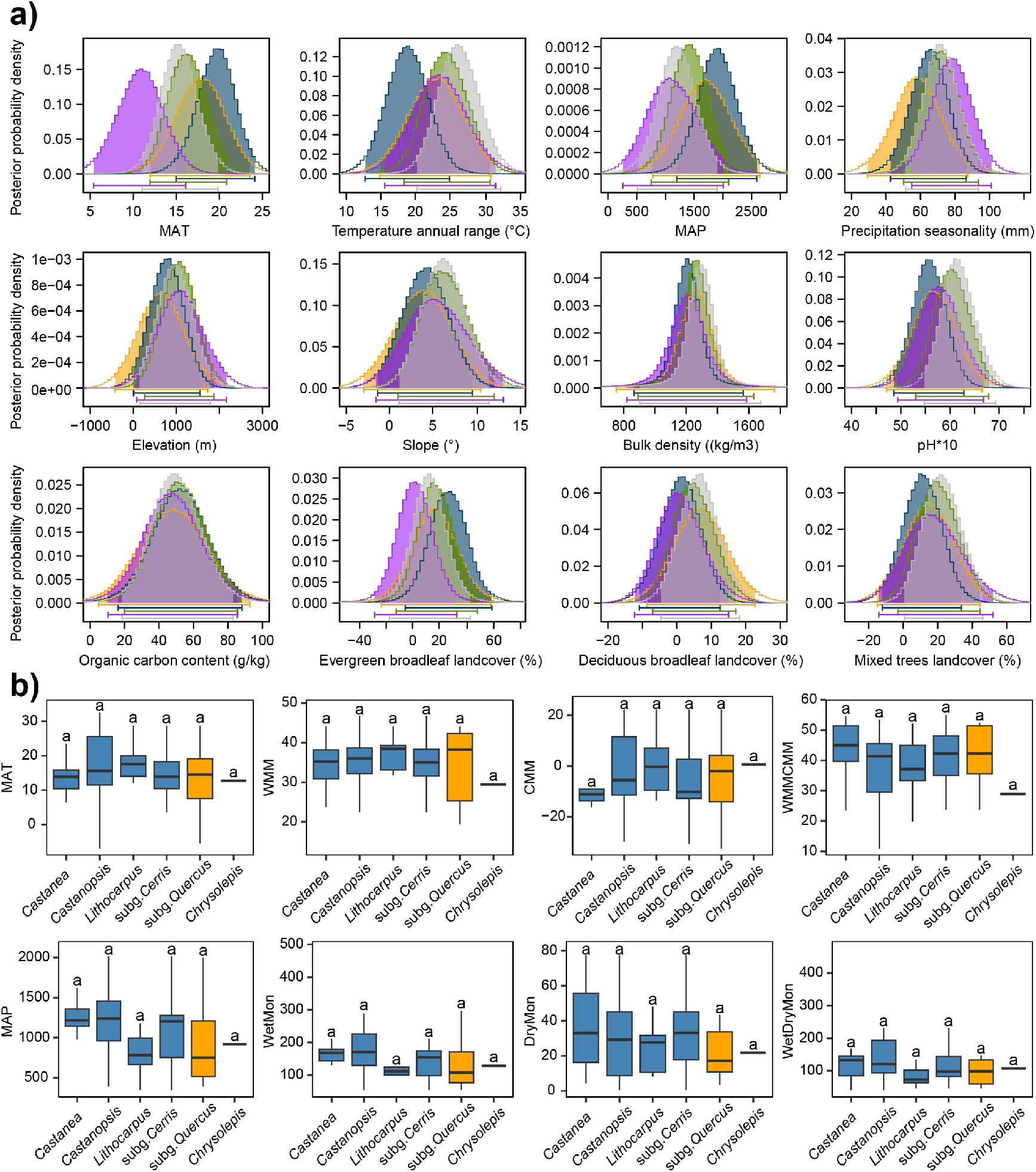
Overlapped ancestral ecological niche space between ancestors of *Quercus* lineages and other genera of Quercoideae involved in deep reticulation. **a)** Posterior probability distributions for the ancestral niche estimation across 12 environmental variables for the ancestors of *Quercus* and relatives based on extant species. The shaded regions and the bars at the bottom of the distributions denote the 95% credibility intervals. Orange, the most recent common ancestor (MRCA) of *Castanea* + *Castanopsis*; blue, the MRCA of *Lithocarpus*; green, the MRCA of the Old World oak clade (i.e., *Q.* subg. *Cerris*); purple, the MRCA of *Chrysolepis*; gray, the MRCA of the New World oak clade (i.e., *Q.* subg. *Quercus*). **b)** Boxplots showing climatic niche space of Eocene fossil taxa of *Quercus* lineages and relatives. The same letters above boxes indicate no significantly climatic difference inferred by function “LSD.test()” in the R package agricolae (de Mendiburu 2009). Blue boxes denote the Old World lineages, while orange one denote the New World lineages. Abbreviations: MAT, mean annual temperature (℃); WMM, warmest month mean surface air temperature (℃); CMM, coldest month mean surface air temperature (℃); WMMCMM, warmest month–coldest month temperature difference (℃); MAP, mean annual precipitation (mm); WetMon, wettest month precipitation (mm); DryMon, driest month precipitation (mm); WetDryMon, wettest month–driest month precipitation difference (mm).

Collectively, these analyses provide multifaceted evidence that geographic and ecological connectivity during the Eocene facilitated widespread gene flow in two major geographic centers—North America and Eurasia—during the initial radiation of Quercoideae (Fig. 2b; Fig. 3–4). These results also suggest that *Quercus* and relatives (Quercoideae) have been engaging in syngameon-type systems throughout their evolutionary history. Although the ancestors of *Quercus* and relatives appear to have had overlapping niches, multiple niche shifts (i.e., shifts in climatic and ecological preferences) in Quercoideae, and particularly in *Quercus* (Supplementary Fig. S32), coincide with post-Eocene climatic changes (Zachos et al. 2001). An early Paleogene syngameon may have facilitated the transfer of adaptive alleles for key functional traits or resulted in enhanced genetic variation; this may have significantly contributed to the evolutionary success of oaks and their relatives (Hipp et al. 2020; Kremer and Hipp 2020), allowing them to adapt to the myriad ecological challenges posed since the Oligocene, including widespread cooling and aridification and major topographical changes (Zachos et al. 2001; Dickinson 2004; Licht et al. 2014; Antonelli et al. 2018).

## CONCLUSION

This work provides new insights into the evolutionary history of *Quercus* and a basis for further studies on this well-known genus. Our results document widespread ancient reticulations between major lineages of *Quercus* and Quercoideae during the initial radiation of Fagaceae. Ancestral reconstructions based on data from extant species or the fossil record both support the plausibility of ancient reticulation events during the early to middle Eocene in two major geographic centers, North America and Eurasia, among co-occurring lineages occupying overlapping ecological niche space. Given the inherent challenges of detecting ancient gene flow, we emphasize the importance of bringing together multiple lines of evidence and analytical approaches for confident reconstruction of ancient reticulation. This includes the dissection of conflicting signals in nuclear and chloroplast datasets using summary statistics, simulations, and network methods; molecular dating of separate genomic compartments; and ancestral range and niche reconstructions leveraging signal from both extant and fossil species. These types of syntheses clarify not only the players but also the time, place, and ecological context of ancient reticulation. This work reveals that hybridization is not only an important recent and ongoing evolutionary force in *Quercus* and relatives, but also has been an important process throughout the long history of this lineage. Our work also provides a methodological roadmap for conducting similar studies in other lineages.

## SUPPLEMENTARY MATERIAL

The resulting DNA alignments, trees, and custom Python and R scripts are available on Dryad Digital Repository: https://doi.org/10.5061/dryad.m0cfxpp5b. Raw sequence data are available at the NCBI SRA (transcriptomes in PRJNA910851; Hyb-Seq data in PRJNA913201). Paleo-climate models can be accessed at the University of Bristol, BRIDGE repository (https://www.paleo.bristol.ac.uk/ummodel/scripts/papers/Valdes_et_al_2021.html)

## Supporting information

Supplementary Methods and Results

## ACKNOWLEDGEMENTS

We thank the following herbaria and their curators and staff for kindly providing herbarium or silica-dried materials: Botanical Research Institute of Texas; California Academy of Sciences; Field Museum of Natural History; Harvard University; Kunming Institute of Botany, Chinese Academy of Sciences; Missouri Botanical Garden; The Ohio State University; Smithsonian Institution; New York Botanical Garden, and University of Texas at Austin. We thank Dr. Lin Bai and Jiajin Wu for help with sampling. We thank the Germplasm Bank of Wild Species at the Kunming Institute of Botany, including the iFlora high performance computing center, for facilitating this study. We thank Dr. Isabel Sanmartín, Dr. Richard Ree, Dr. Joe Walker, and another anonymous reviewer for their helpful suggestions for this paper.

