## Supplementary Tables and Figures for "Phylogenomic Analyses Reveal Widespread Gene Flow During the Early Radiation of Oaks and Relatives (Fagaceae: Quercoideae)"

### SUPPLEMENTARY FIGURES

**FIGURE S1.** Overview of the Hyb-Seq datasets (red boxes) and analyses conducted in this study.

**FIGURE S2.** Overview of the transcriptome datasets (red boxes) and analyses conducted in this study.

**FIGURE S3.** Overview of plastome datasets (red box) and analyses conducted in this study.

**FIGURE S4.** Species tree of oaks and relatives inferred by ASTRAL-III based on the HYB-89MO dataset including 89 loci from 431 species.

**FIGURE S5.** Species tree of oaks and relatives inferred by ASTRAL-III based on the HYB-98RT dataset including 98 loci from 431 species.

**FIGURE S6.** Maximum likelihood (ML) tree of oaks and relatives inferred by RAxML based on the concatenated supermatrix (the HYB-89MO dataset) including 89 loci from 431 species under an unpartitioned GTR-GAMMA model.

**FIGURE S7.** ML tree of oaks and relatives inferred by RAxML based on the concatenated supermatrix (the HYB-98RT dataset) including 98 loci from 431 species under an unpartitioned GTR-GAMMA model.

**FIGURE S8.** Species tree of oaks and relatives inferred by ASTRAL-III based on the RNA-977MO dataset including 977 loci from 89 species.

**FIGURE S9.** Species tree of oaks and relatives inferred by ASTRAL-III based on the RNA-2821RT dataset including 2821 loci from 89 species.

**FIGURE S10.** ML tree of oaks and relatives inferred by RAxML based on the concatenated supermatrix (the RNA-977MO dataset) including 977 loci from 89 species under an unpartitioned GTR-GAMMA model.

**FIGURE S11.** ML tree of oaks and relatives inferred by RAxML based the concatenated supermatrix (the RNA-2821RT dataset) including 2821 loci from 89 species under an unpartitioned GTR-GAMMA model.

**FIGURE S12.** The distribution of gene-wise and site-wise phylogenetic signal for alternative topologies of five uncertain deep branches based on the HYB-98RT dataset.

**FIGURE S13.** The distribution of gene-wise and site-wise phylogenetic signal for alternative topologies of five uncertain deep branches based on the RNA-2821RT dataset.

**FIGURE S14.** Clade frequencies of the chloroplast trees simulated under the coalescent (with the guide tree scaled by a factor of four), mapped on the cladogram of the concatenated ML tree (the HYB-98RT dataset).

**FIGURE S15.** Clade frequencies of the chloroplast trees simulated under the coalescent (with the guide tree scaled by a factor of two), mapped on the cladogram of the concatenated ML tree (the HYB-98RT dataset).

**FIGURE S16.** Species networks from reduced-representation data sets of oaks and Fagaceae.

**FIGURE S17.** Levels of intergenic discordance and uninformativeness with respect to major clades of oaks and Quercoideae.

**FIGURE S18.** Phyparts results based on the 98 gene trees from the HYB-98RT dataset, mapped against the ASTRAL species tree.

**FIGURE S19.** Phyparts result based on the 2821 gene trees of the RNA-2821RT dataset, mapped against the ASTRAL species tree.

**FIGURE S20.** Cophylogeny showing incongruence between the nuclear ASTRAL (left; the HYB-98RT dataset) and unpartitioned chloroplast ML (right) trees with coalescent simulation results from the nuclear guide tree scaled by a factor of four.

**FIGURE S21.** Tanglegram comparing the nuclear ASTRAL (left; the RNA-2821RT dataset) and chloroplast (right) phylogenies optimized in Dendroscope, with coalescent simulation results from the nuclear guide tree scaled by a factor of four.

**FIGURE S22.** Cophylogeny showing incongruence between the nuclear ASTRAL (left; the HYB-98RT dataset) and unpartitioned chloroplast (right) ML trees with coalescent simulation results from the nuclear guide tree scaled by a factor of two.

**FIGURE S23.** Cophylogeny showing incongruence between the nuclear ASTRAL (left; the RNA-2821RT dataset) and unpartitioned chloroplast (right) ML trees with coalescent simulation results from the nuclear guide tree scaled by a factor of two.

**FIGURE S24.** Proportions of positive tests for various introgression signatures visualized on the time-calibrated tree based on the HYB-98RT dataset.

**FIGURE S25.** Proportions of positive tests for various introgression signatures visualized on the time-calibrated tree based on the RNA-2821RT dataset.

**FIGURE S26.** Plots showing the relative contributions of incomplete lineage sorting (ILS), gene flow, and gene tree estimation error to the observed gene-tree discordance, across two datasets and two taxonomic levels.

**FIGURE S27.** Cladogram of the chloroplast ML tree of oaks and relatives inferred by RAxML based on the concatenated plastome supermatrix including 223 species under an unpartitioned GTR-GAMMA model.

**FIGURE S28.** Cladogram of the chloroplast ML tree of oaks and relatives inferred by RAxML based on the concatenated plastome supermatrix including 223 species under a GTR-GAMMA model partitioned by genome region (LSC, SSC and IR).

**FIGURE S29.** Ancestral range estimation of Fagaceae under the DEC model, based on the MCC tree of the HYB-MO89 dataset.

**FIGURE S30.** Potential distribution of the major lineages of *Quercus* and Quercoideae from Late Paleocene to Late Eocene as inferred by fossil-based ecological niche modeling under PEO strategy.

**FIGURE S31.** The fossil distribution of *Chrysolepis* and *Notholithocarpus* during the Paleogene.

**FIGURE S32.** Median values of ecological niche space across Fagaceae phylogeny, based on ancestral niche reconstructions on the tree produced from the Hyb-Seq dataset.

**FIGURE S33.** Boxplots showing climatic niche space from Early Eocene to Late Oligocene of fossil taxa of *Quercus* lineages and relatives.

**FIGURE S34.** Genomic locations of the targeted loci.

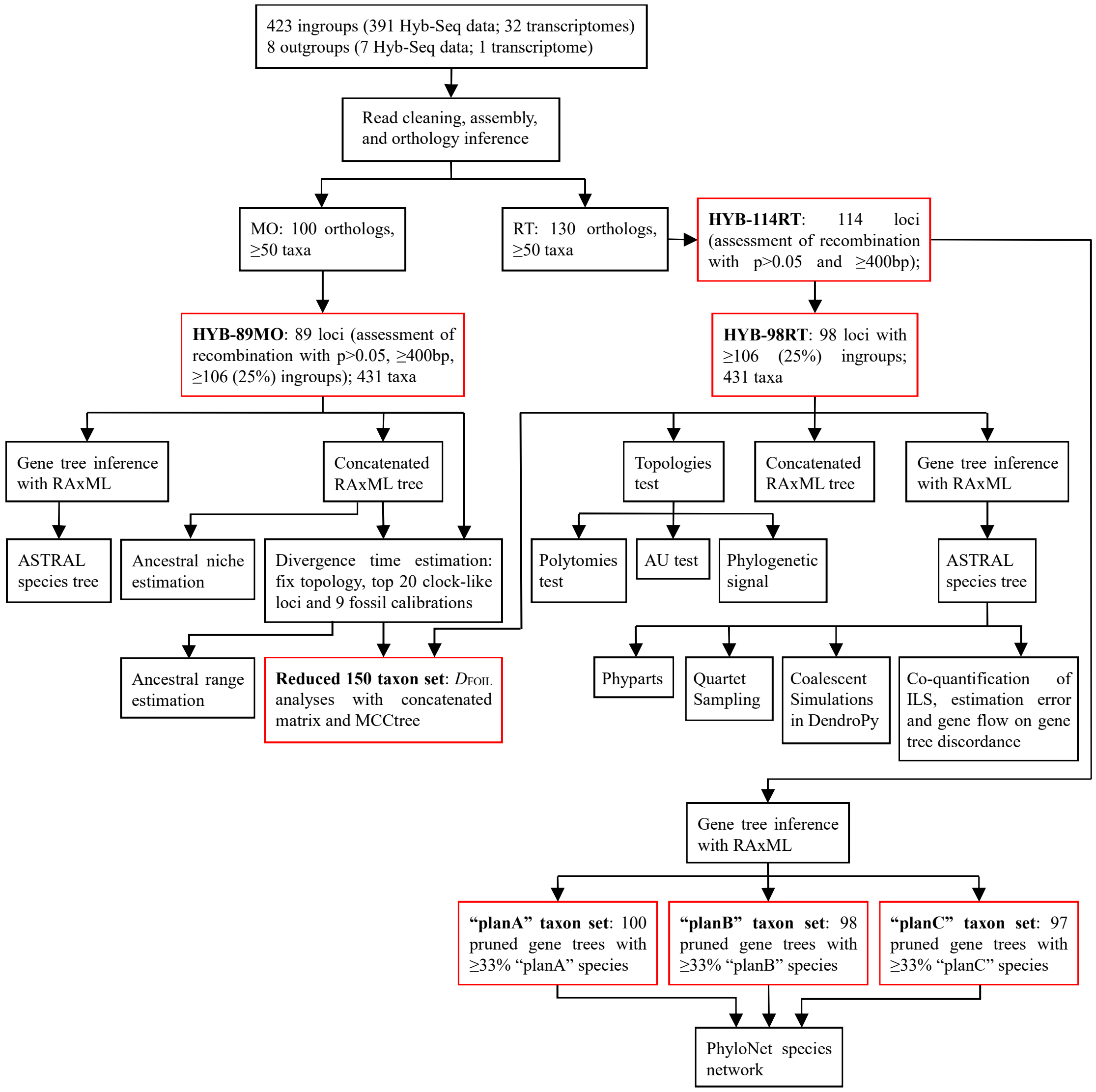


**FIGURE S1. Overview of the Hyb-Seq datasets (red boxes) and analyses conducted in this study.** MO, monophyletic outgroup approach for orthology inferences following the pipeline of Yang and Smith (2014); RT, rooted ingroup approach for orthology inferences following the pipeline of Yang and Smith (2014).

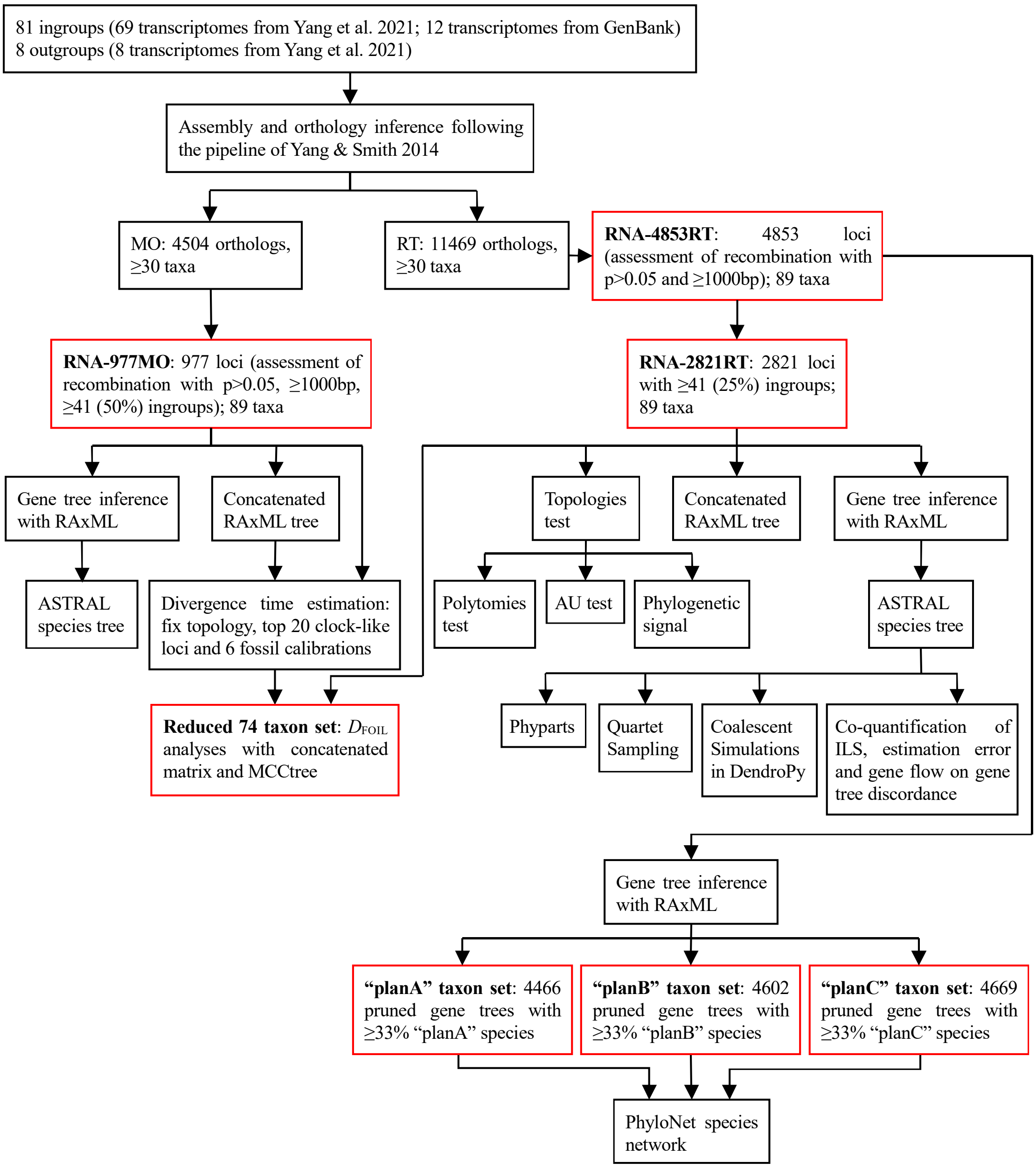


**FIGURE S2. Overview of the transcriptome datasets (red boxes) and analyses conducted in this study.**

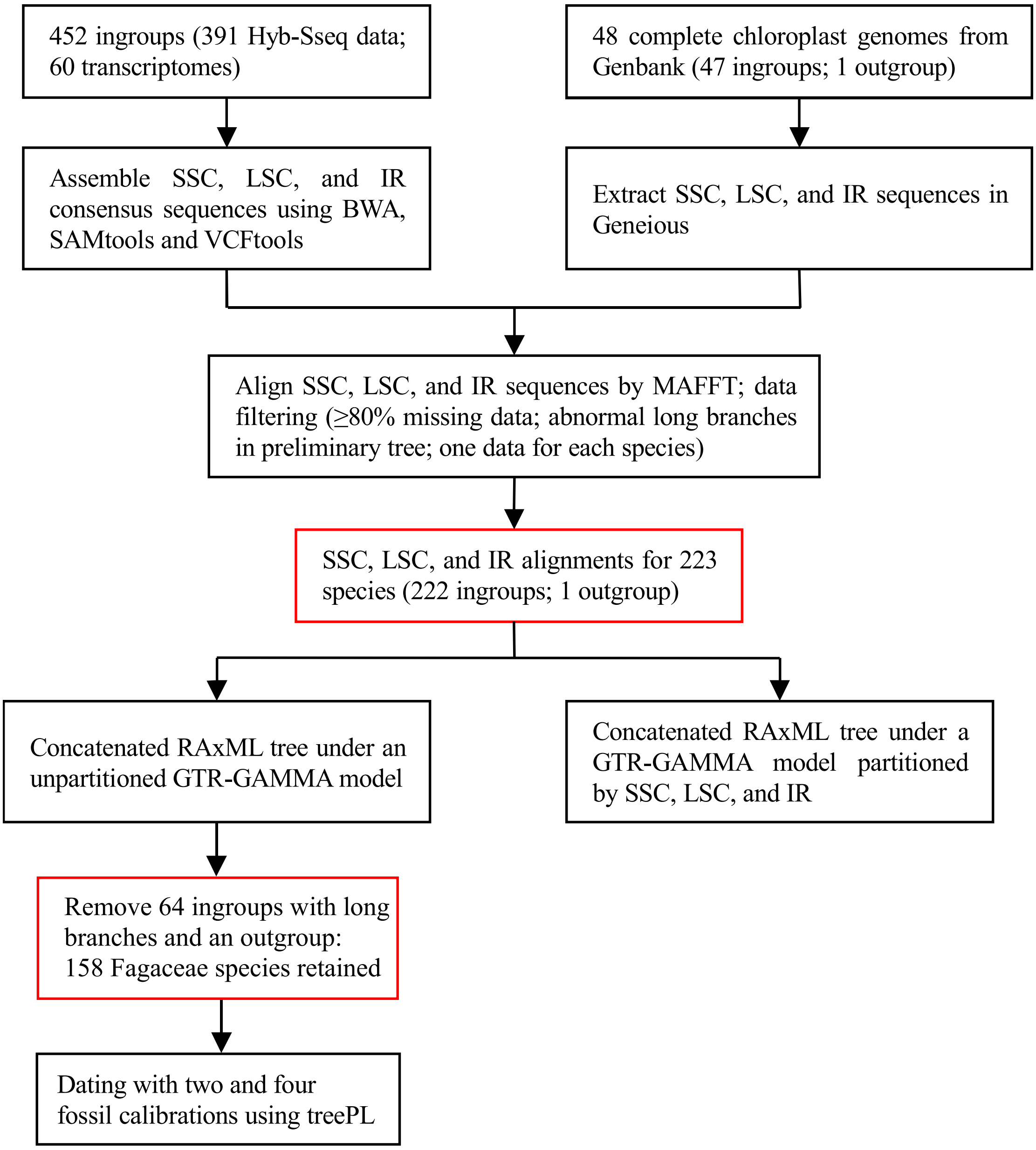


**FIGURE S3. Overview of plastome datasets (red box) and analyses conducted in this study.** LSC, SSC, and IR refer to the large-single-copy region, small-single-copy region, and inverted repeat of the complete chloroplast genome, respectively.

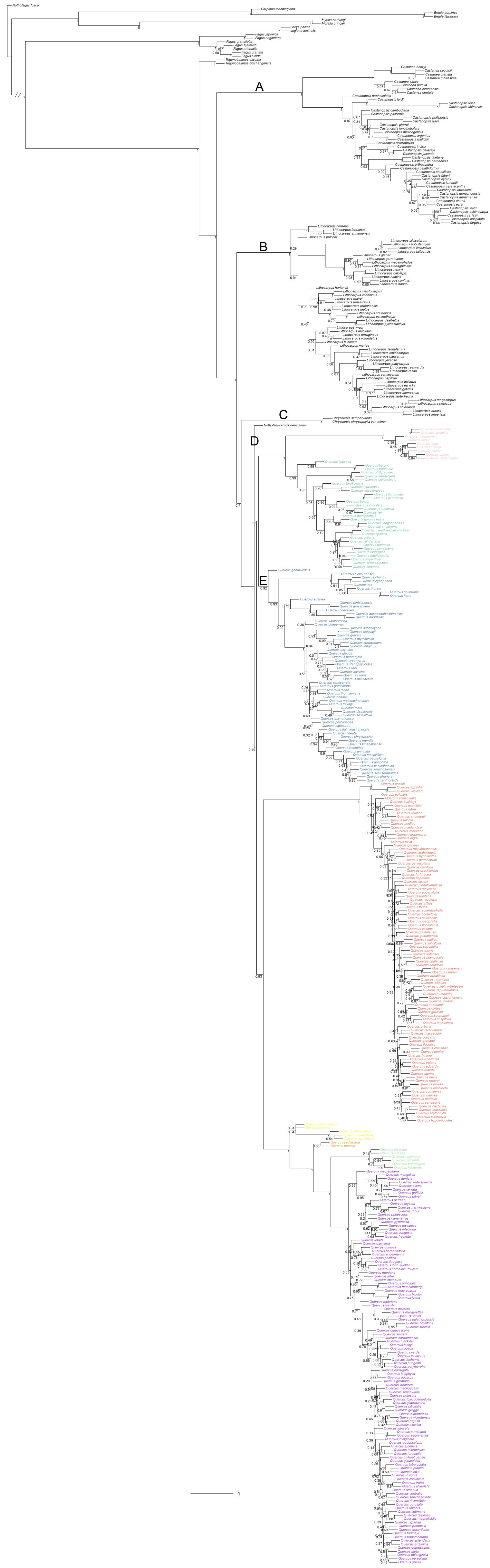
A
B
C
D
E
1

**FIGURE S4. Species tree of oaks and relatives inferred by ASTRAL-III based on the HYB-89MO dataset including 89 loci from 431 species .** Local posterior probabilities are shown above branches. A–E denote the five uncertain deep branches (Supplementary Table S5) investigated in this study. Colored tip labels denote the eight sections of oaks: pink, sect. *Cerris*; green, sect. *Ilex*; blue, sect. *Cyclobalanopsis*; red, sect. *Lobatae*; yellow, sect. *Protobalanus*; orange, sect. *Ponticae*; light green, sect. *Virentes*; and purple, sect. *Quercus*.

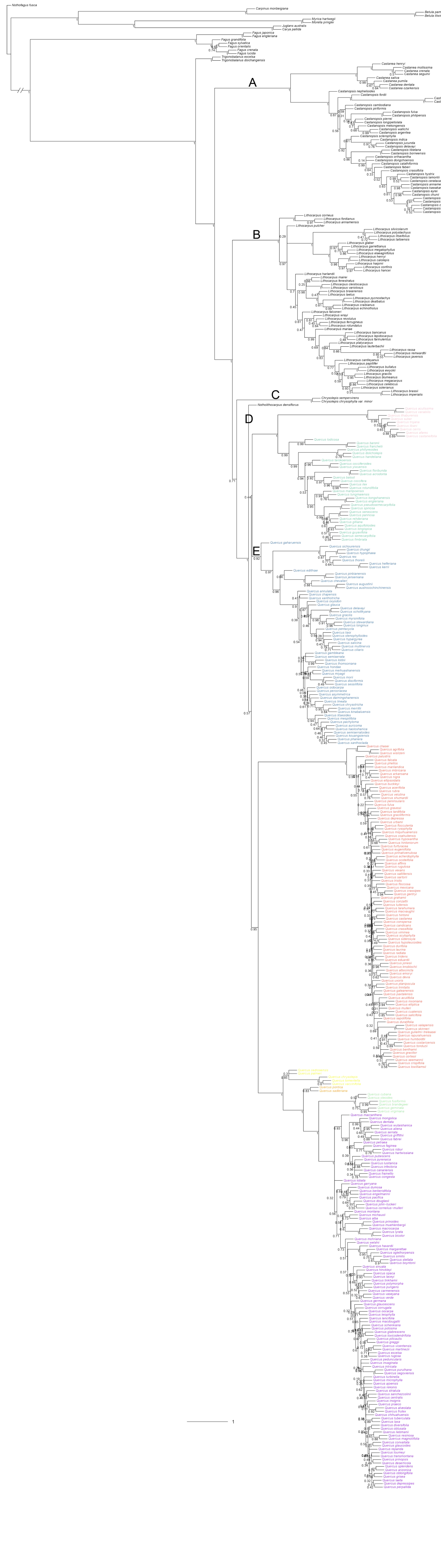


117

**FIGURE S5. Species tree of oaks and relatives inferred by ASTRAL-III based on the HYB-98RT dataset including 98 loci from 431 species.** Local posterior probabilities are shown above branches. A–E denote the five uncertain deep branches (Supplementary Table S5) investigated in this study. Tip labels are colored following the color scheme in Supplementary Figure S4.

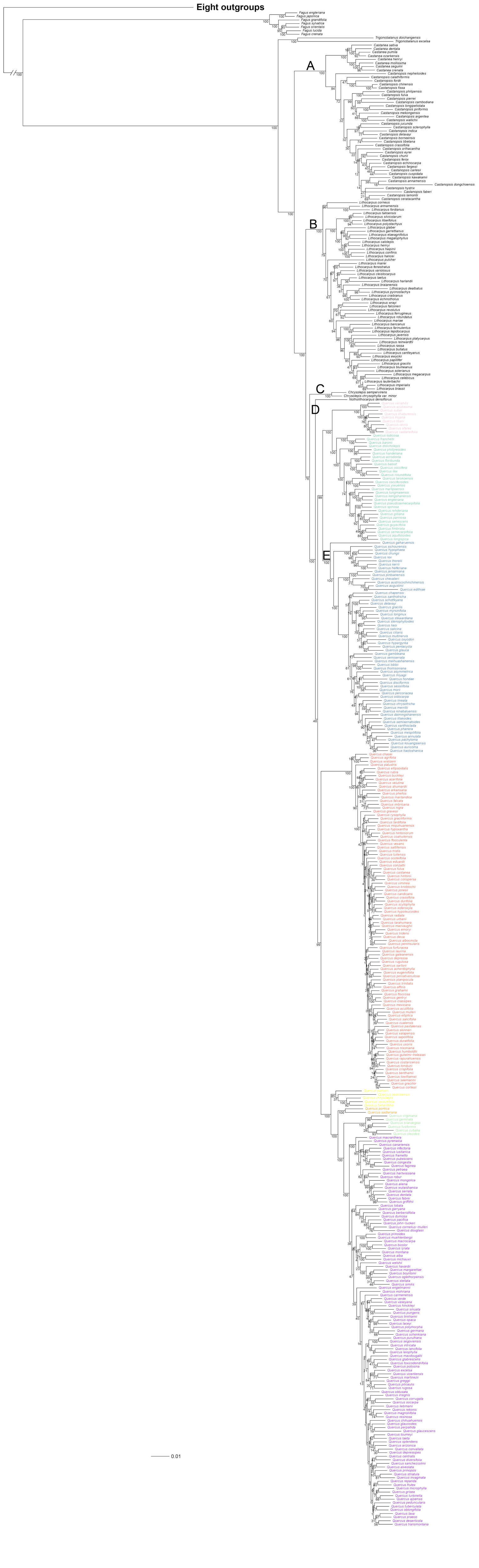
Eight outgroups
A
B
C
D
E
0.01

**FIGURE S6. Maximum likelihood (ML) tree of oaks and relatives inferred by RAxML based on the concatenated supermatrix (the HYB-89MO dataset) including 89 loci from 431 species under an unpartitioned GTR-GAMMA model.** Bootstrap values are shown above branches. A–E denote the five uncertain deep branches (Supplementary Table S5) investigated in this study. Tip labels are colored following the color scheme in Supplementary Figure S4.

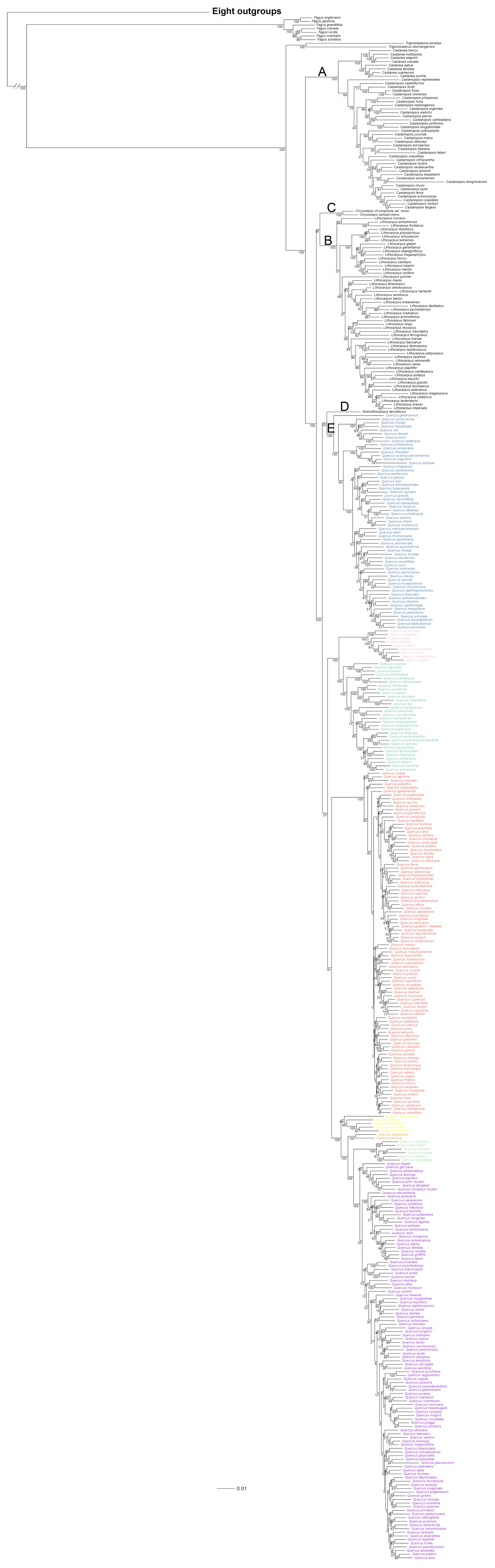
Eight outgroups
A
B
C
D
E
0.01

**FIGURE S7. ML tree of oaks and relatives inferred by RAxML based on the concatenated supermatrix (the HYB-98RT dataset) including 98 loci of 431 species and under an unpartitioned GTR-GAMMA model.** Bootstrap values are shown above branches. A–E denote the five uncertain deep branches (Supplementary Table S5) investigated in this study. Tip labels are colored following the color scheme in Supplementary Figure S4.

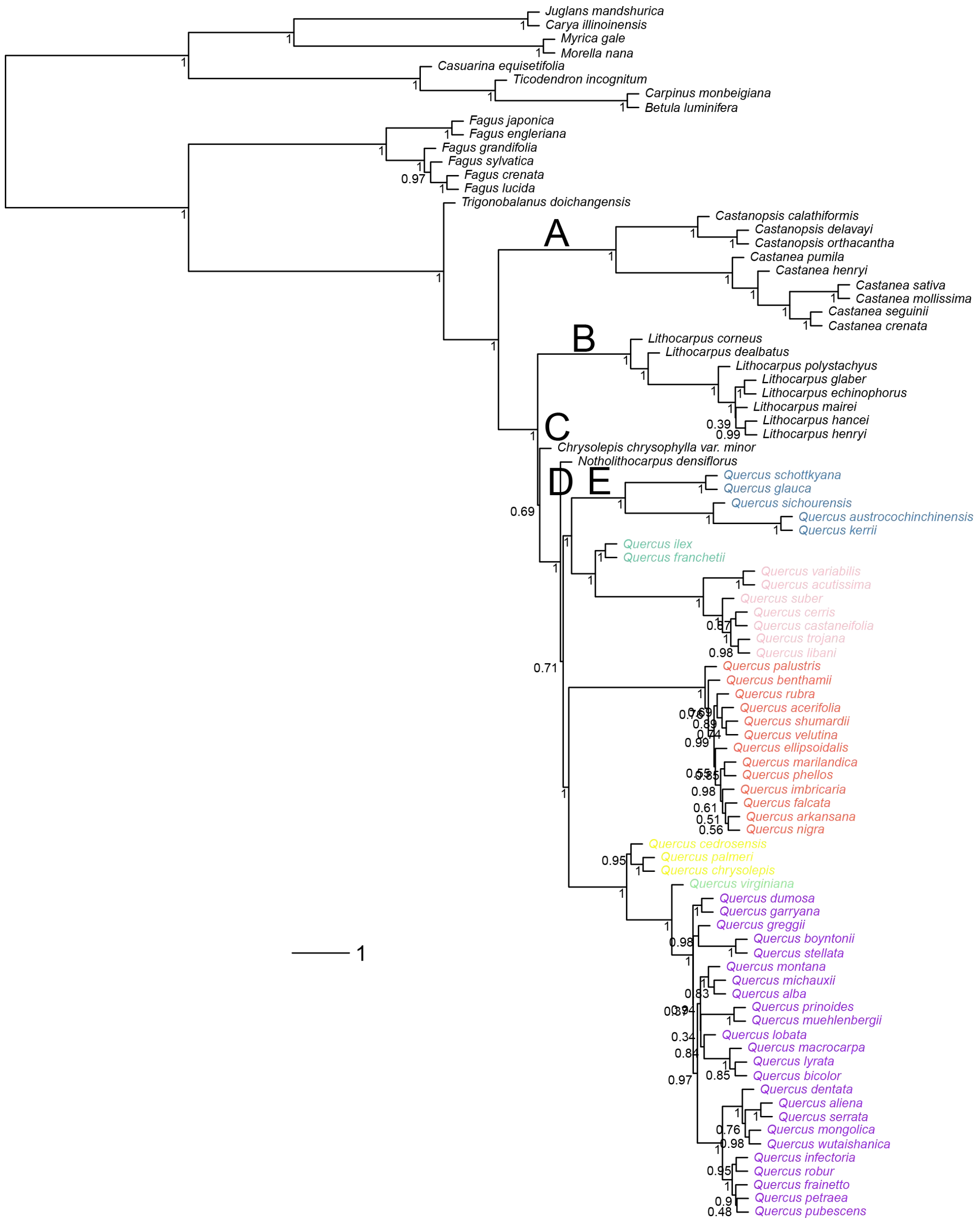


**FIGURE S8. Species tree of oaks and relatives inferred by ASTRAL-III based on the RNA-977MO dataset including 977 loci from 89 species.** Local posterior probabilities are shown above branches. A–E denote the five uncertain deep branches (Supplementary Table S5) investigated in this study. Tip labels are colored following the color scheme in Supplementary Figure S4.

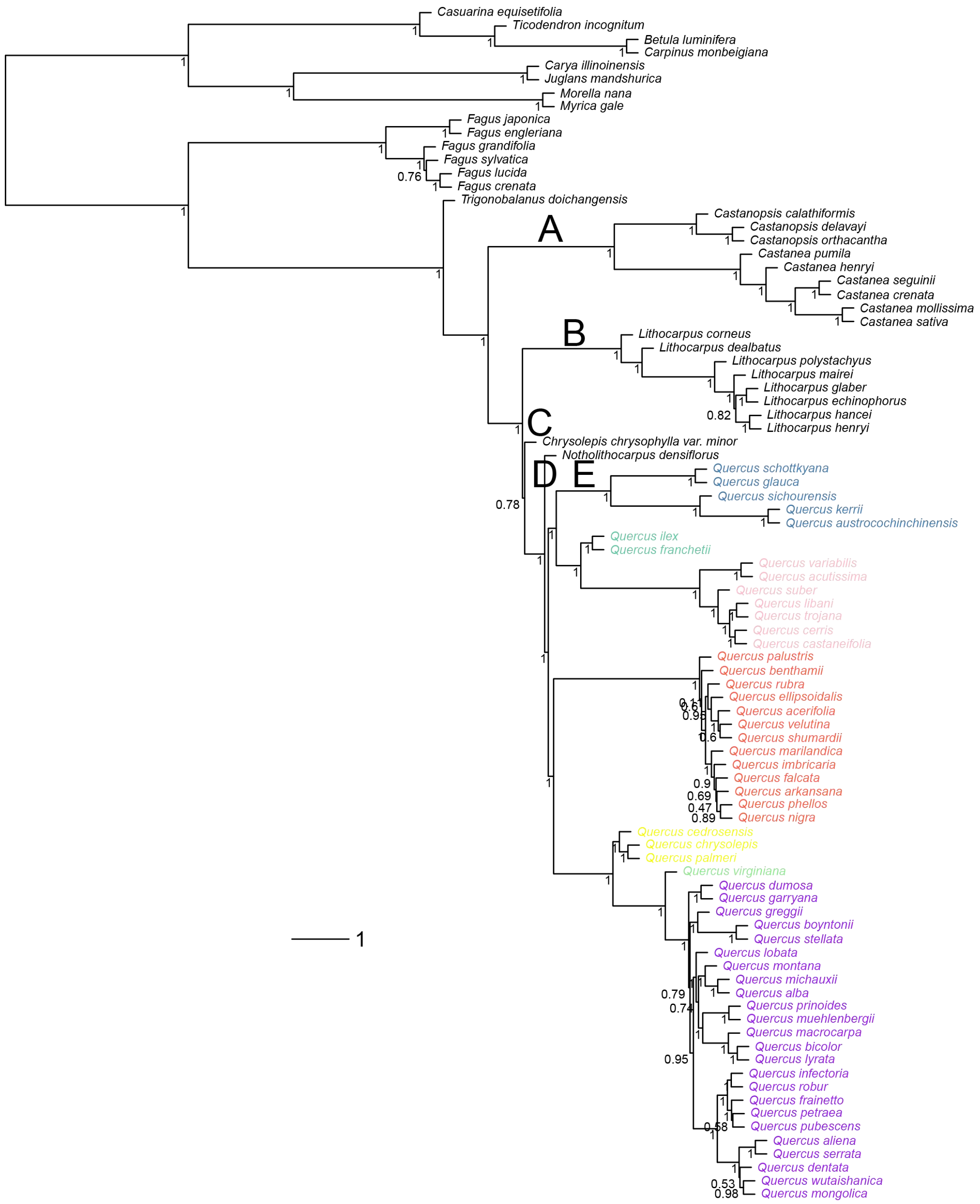


**FIGURE S9. Species tree of oaks and relatives inferred by ASTRAL-III based on the RNA-2821RT dataset including 2821 loci from 89 species.** Local posterior probabilities are shown above branches. A–E denote the five uncertain deep branches (Supplementary Table S5) investigated in this study. Tip labels are colored following the color scheme as in Supplementary Figure S4.

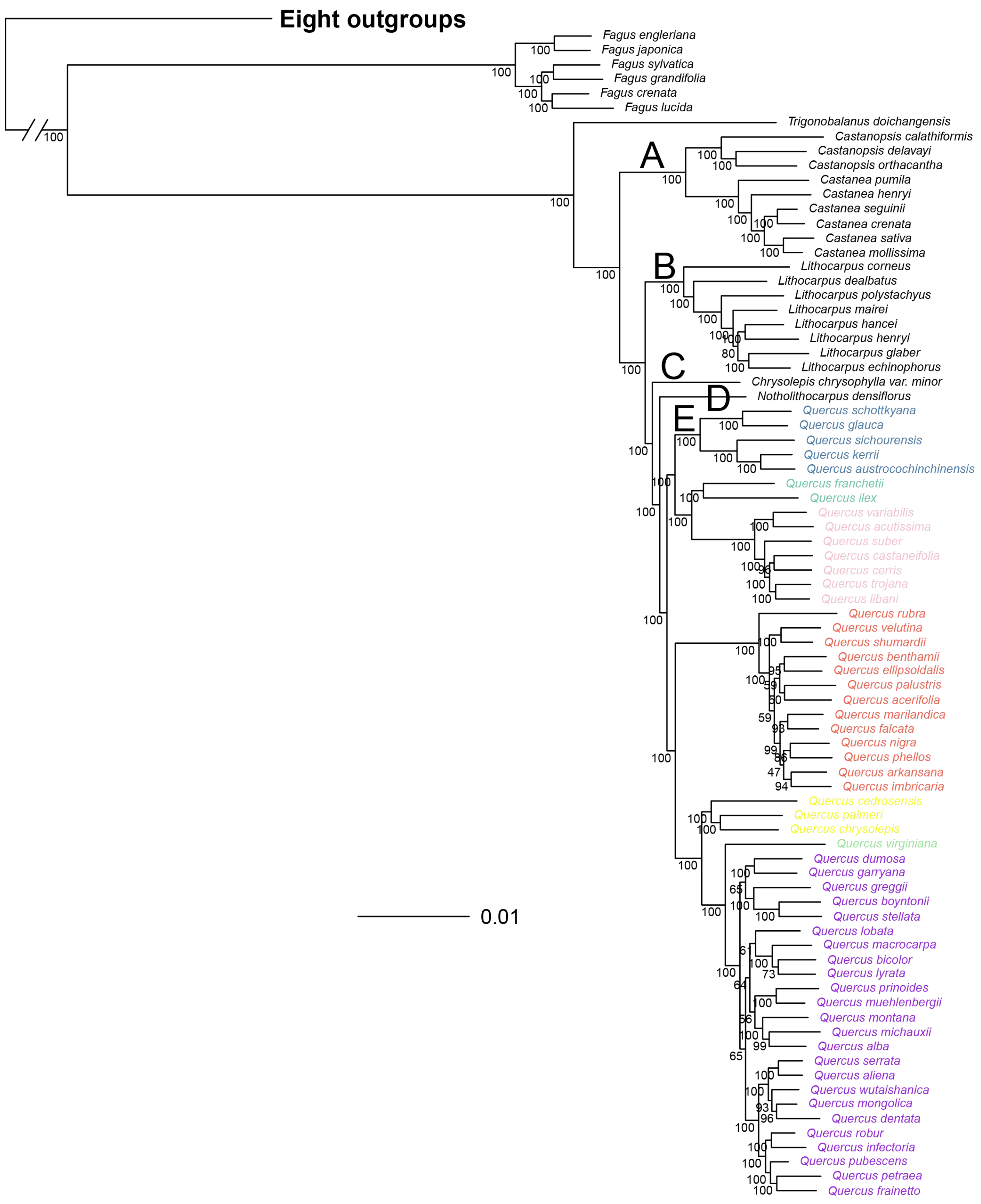


**FIGURE S10. ML tree of oaks and relatives inferred by RAxML based on the concatenated supermatrix (the RNA-977MO dataset) including 977 loci from 89 species under an unpartitioned GTR-GAMMA model.** Bootstrap values are shown above branches. A–E denote the five uncertain deep branches (Supplementary Table S5) investigated in this study. Tip labels are colored by the same color scheme as in Supplementary Figure S4.

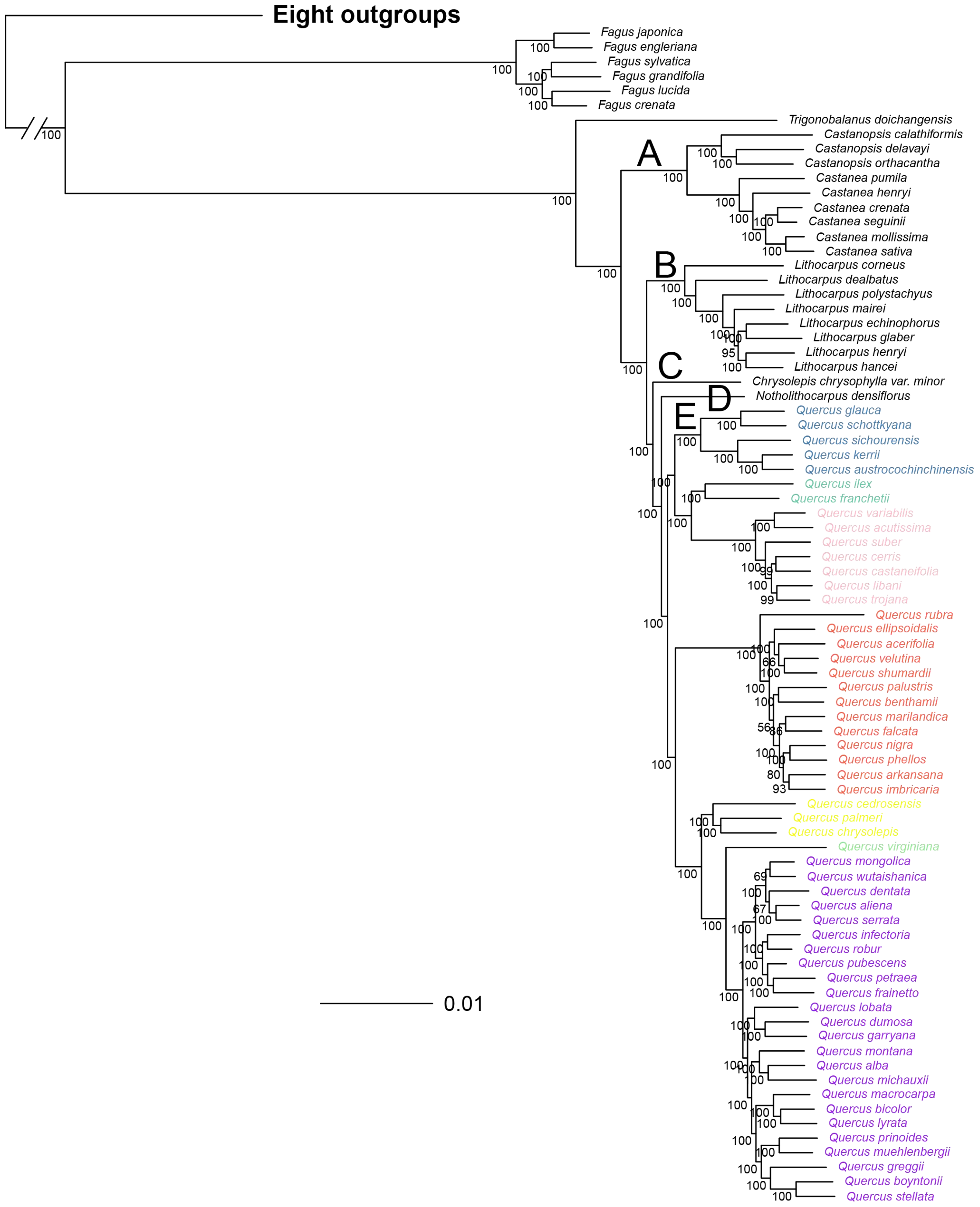


**FIGURE S11. ML tree of oaks and relatives inferred by RAxML based the concatenated supermatrix (the RNA-2821RT dataset) including 2821 loci from 89 species under an unpartitioned GTR-GAMMA model.** Bootstrap values are shown above branches. A–E denote the five uncertain deep branches (Supplementary Table S5) investigated in this study. Tip labels are colored by the same color scheme as in Supplementary Figure S4.

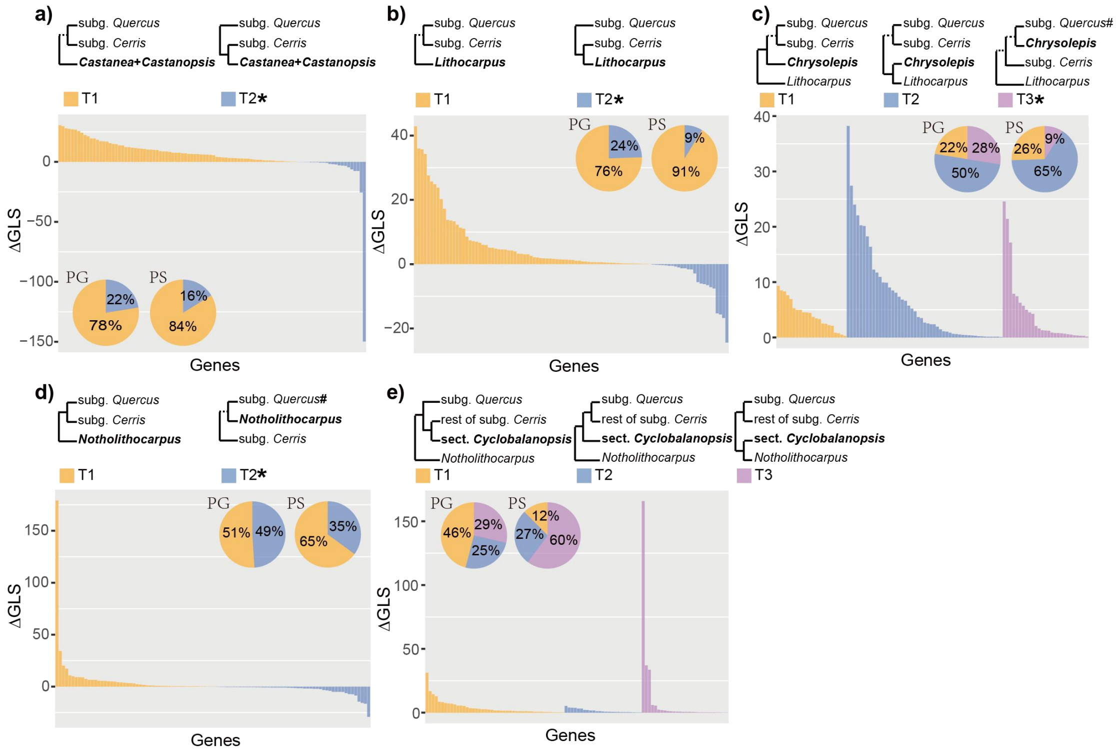


**FIGURE S12. The distribution of gene-wise and site-wise phylogenetic signal for alternative topologies of five uncertain deep branches based on the HYB-98RT dataset. a)** The position of *Castanea* + *Castanopsis*. **b)** The position of *Lithocarpus*. **c)** The position of *Chrysolepis*. **d)** The position of *Notholithocarpus*. **e)** The position of *Quercus* sect. *Cyclobalanopsis*. Topologies of T1 are extracted from the ASTRAL and concatenated ML trees (with the exception of the ML tree from HYB-98RT) in this study. The alternative topologies (T2 and T3) with an asterisk are from chloroplast trees in this and previous studies (e.g., Xiang et al. 2014; Simeone et al. 2016; Yang et al. 2021), and the remaining alternative topologies (T2 and T3) are based on the concatenated ML tree of the HYB-98RT dataset and nuclear trees of previous studies (e.g., Manos et al. 1999, 2001, 2008; Hubert et al. 2014). Dashed branches denote regions of the 3(4)-taxon tree where other lineages are not shown. The "#" means subg. *Quercus* here with sect. *Lobatae* excluded. Abbreviations: PG, percentage of genes supporting a topology; PS, percentage of sites supporting a topology.

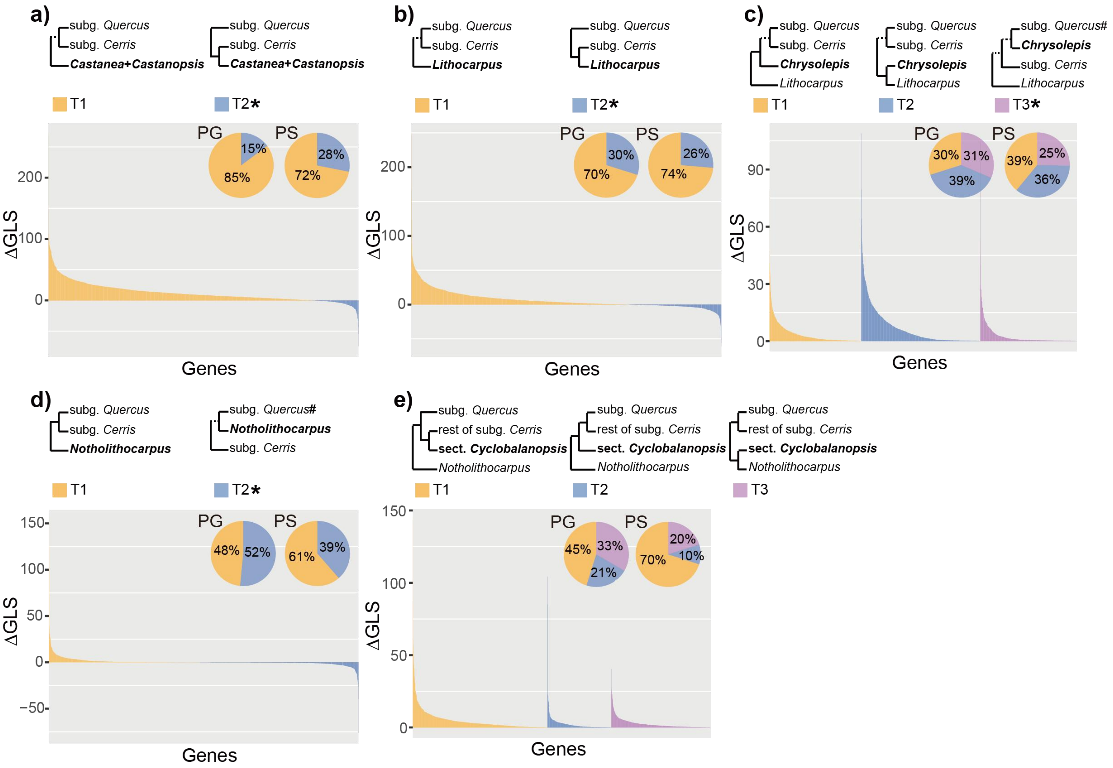


**FIGURE S13. The distribution of gene-wise and site-wise phylogenetic signal for alternative topologies of five uncertain deep branches based on the RNA-2821RT dataset. a)** The position of *Castanea* + *Castanopsis*. **b)** The position of *Lithocarpus*. **c)** The position of *Chrysolepis*. **d)** The position of *Notholithocarpus*. **e)** The position of *Quercus* subg. *Cyclobalanopsis*. Dashed branches denote other clades (which are not shown in the 3(4)-taxon tree) located at this branch. "#" means subg. *Quercus* here with sect. *Lobatae* excluded. Abbreviations: PG, percentage of genes supporting a topology; PS, percentage of sites supporting a topology.

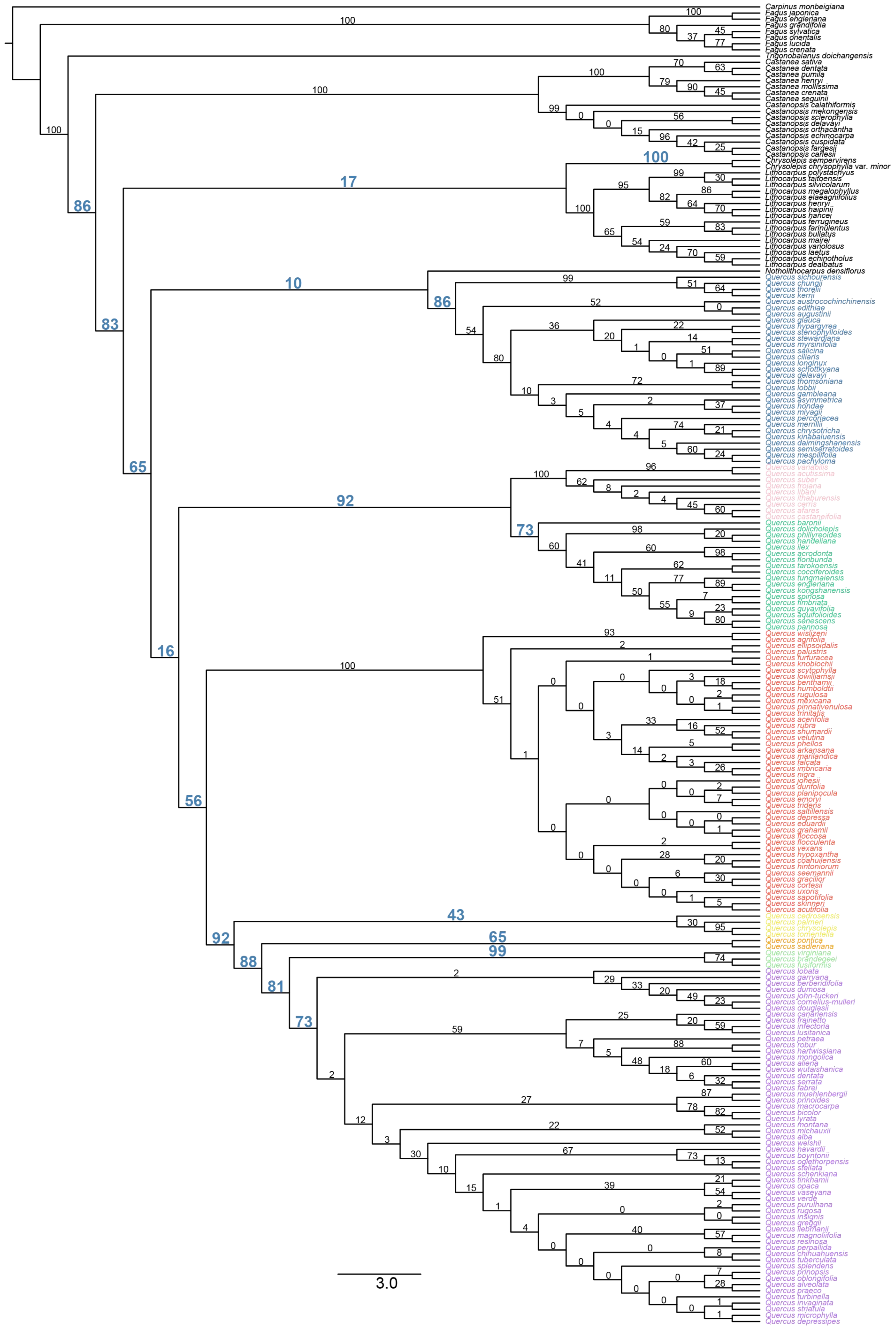


**FIGURE S14. Clade frequencies of the chloroplast trees simulated under the coalescent (with the guide tree scaled by a factor of four), mapped on the**

**cladogram of the concatenated ML tree (the HYB-98RT dataset).** Cladogram of ML tree shown here with 215 tips is pruned from the ML tree of the HYB-98RT dataset with 431 tips (Supplementary Figure S7) to match the taxon set of the chloroplast ML tree. Clade probabilities are shown above the branches. Clade probabilities associated with deep cytonuclear discordance are blue, bold, and enlarged. Tip labels are colored following the color scheme in Supplementary Figure S4.

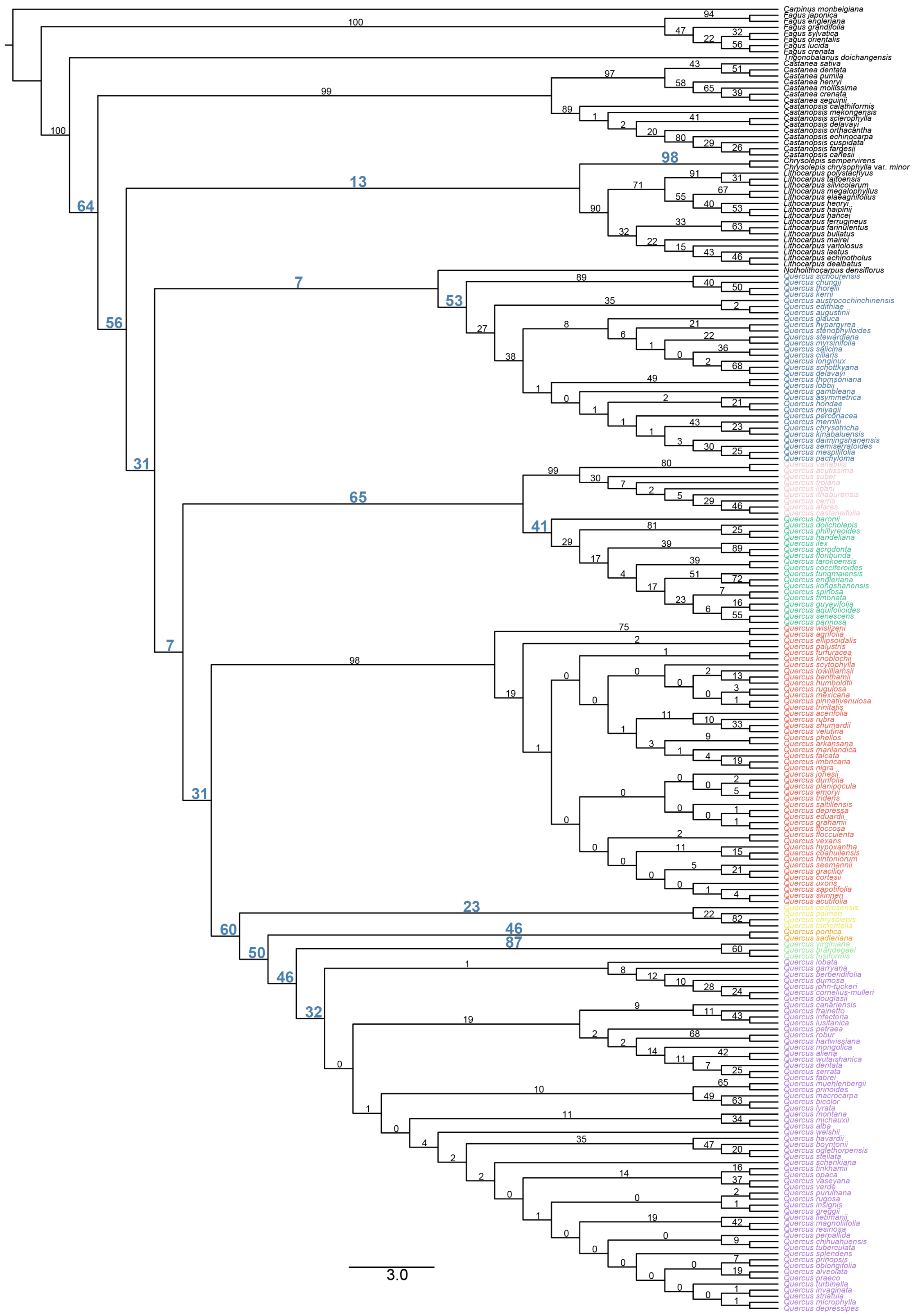

**FIGURE S15. Clade frequencies of the chloroplast trees simulated under the coalescent (with the guide tree scaled by a factor of two), mapped on the cladogram of the concatenated ML tree (the HYB-98RT dataset).** Cladogram of

ML tree shown here with 215 tips is pruned from the ML tree of the HYB-98RT dataset with 431 tips (Supplementary Figure S7) to match the taxon set of the chloroplast ML tree. Clade probabilities are shown above branches. Clade probabilities associated with deep cytonuclear discordance are blue, bold, and enlarged. Tip labels are colored following the color scheme in Supplementary Figure S4.

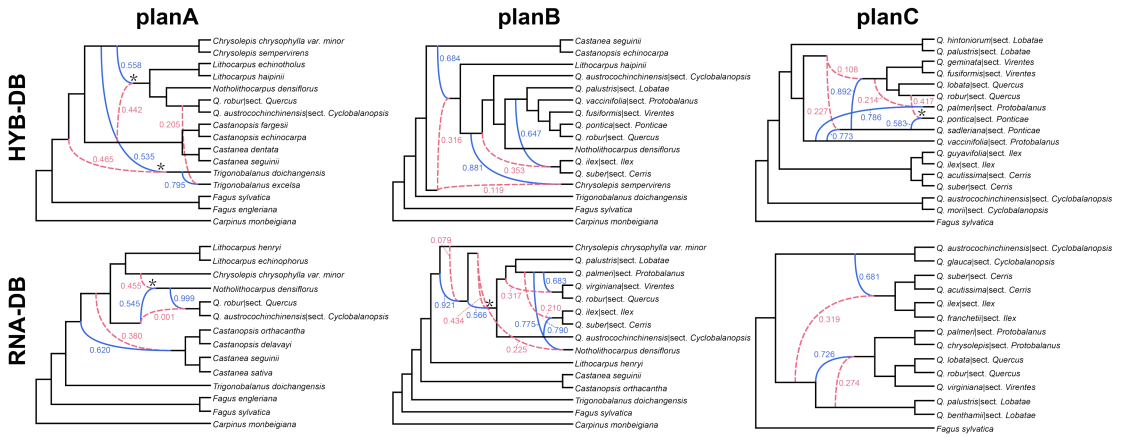


**FIGURE S16. Species networks from reduced-representation datasets of oaks and Fagaceae. a)** Optimal network under the “planA” taxon set of Fagaceae using the Hyb-Seq dataset (represented by the HYB-114RT dataset here). **b)** Optimal network under the “planB” taxon set of Fagaceae using the Hyb-Seq dataset. **c)** Optimal network under the “planC” taxon set of oaks using the Hyb-Seq dataset. **d)** Optimal network under the “planA” taxon set of Fagaceae using the transcriptome dataset (represented by the RNA-4853RT dataset here). **e)** Optimal network under the “planB” taxon set of Fagaceae using the transcriptome dataset. **f)** Optimal network under the “planC” taxon set of oaks using the transcriptome dataset. The optimal networks of the three strategies of representative sampling and two datasets (see Supplementary Methods for details) show incongruent scenarios but all basically indicate that complex reticulate evolution occurred in the early history of oaks and subfamily Quercoideae. Dashed red curved branches denote the minor edge of a reticulate node while solid blue curved branches denote the major edge of a reticulate node. Values next to curved branches denote the inheritance probabilities. A reticulate node with an asterisk denotes that its parents contributed relatively equivalent inheritance probabilities (~ 0.5 vs. 0.5). Abbreviations: sect. CY, sect. *Cyclobalanopsis*.

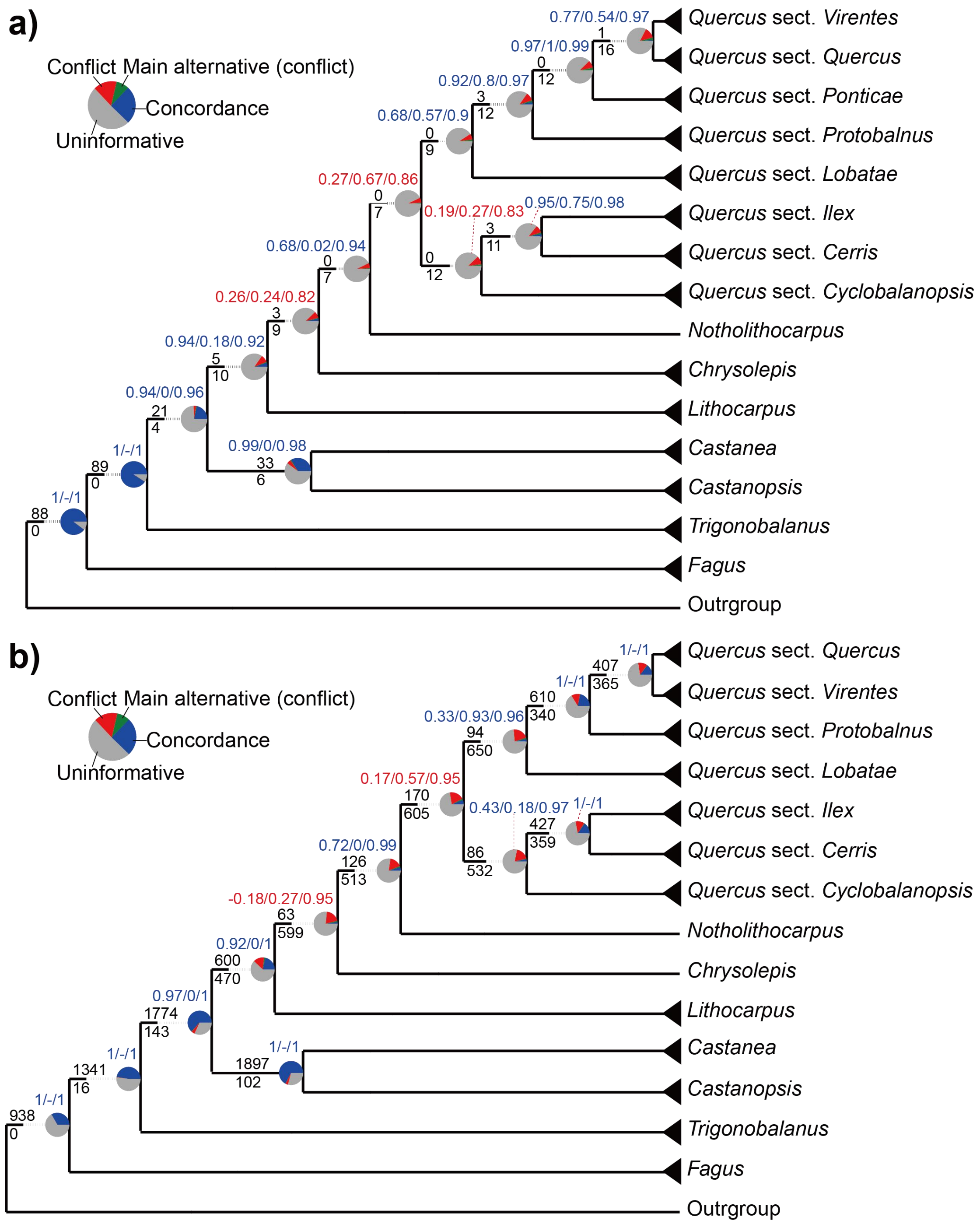


**FIGURE S17. Levels of intergenic discordance and uninformativeness with respect to major clades of oaks and Quercoideae. a)** Concordance analyses for the HYB-98RT dataset. **b)** Concordance analyses for the RNA-2821RT dataset. The ASTRAL tree is used as the mapping tree in phyparts and Quartet Sampling (QS) analyses. Only results for major clades are shown (see Supplementary Fig. S18-S19 for phyparts results of full-taxa sets). Blue, green, red, and grey in each pie chart represent the percentage of gene trees that support that clade, a main alternative topology, all of the remaining alternatives, and are uninformative (that have less than 70% bootstrap support or inadequate taxon sampling), respectively. Numbers above the branches indicate the number of gene trees concordant with each clade, and numbers below branches indicate the number of gene trees in conflict with that clade. The number at the left corner of each pie chart is the QS score (quartet concordance / quartet differential / quartet informativeness). Red QS scores indicate weakly supported branches while blue QS scores indicate strongly supported branches.

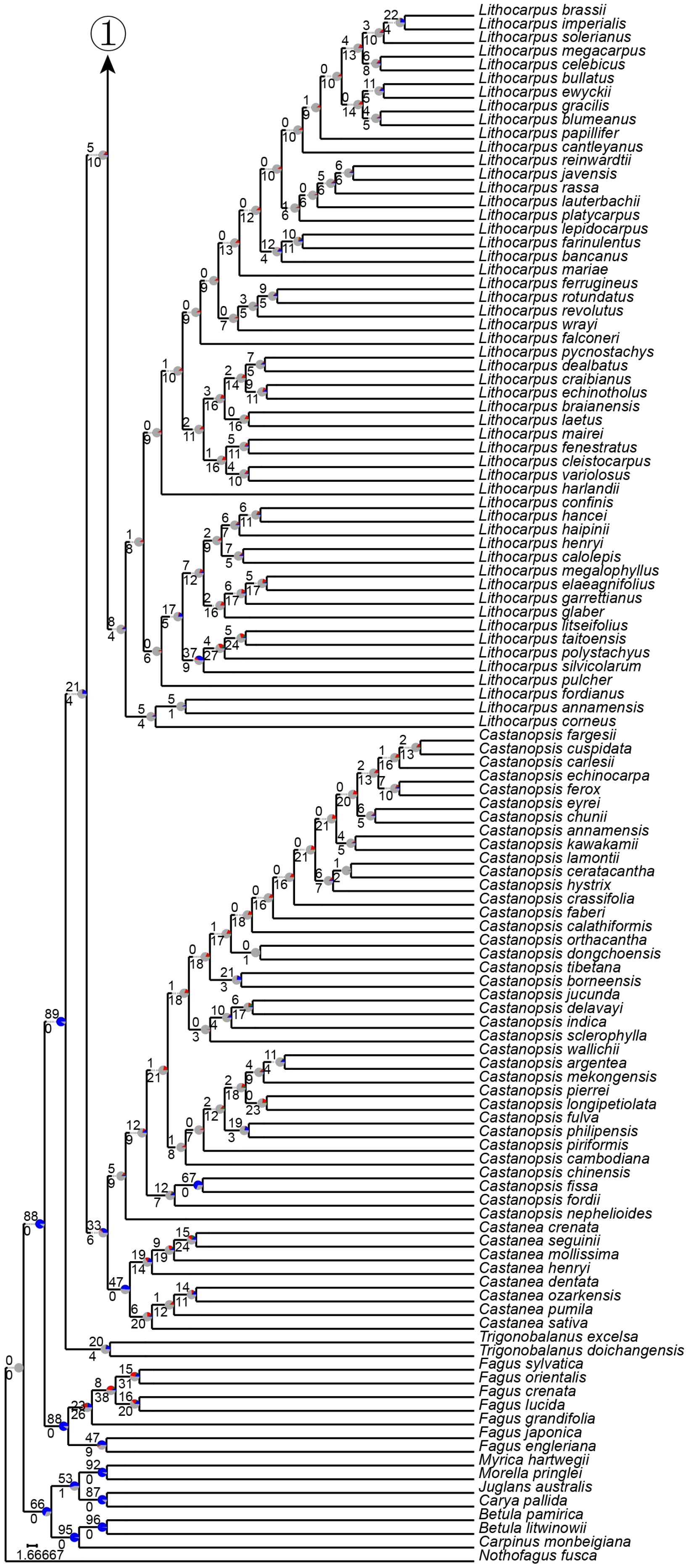
①
Lithocarpus brassii
Lithocarpus imperialis
Lithocarpus solerianus
Lithocarpus megacarpus
Lithocarpus celebicus
Lithocarpus bullatus
Lithocarpus ewyckii
Lithocarpus gracilis
Lithocarpus blumeanus
Lithocarpus papillifer
Lithocarpus cantleyanus
Lithocarpus reinwardtii
Lithocarpus javensis
Lithocarpus rassa
Lithocarpus lauterbachii
Lithocarpus platycarpus
Lithocarpus lepidocarpus
Lithocarpus farinulentus
Lithocarpus bancanus
Lithocarpus mariae
Lithocarpus ferrugineus
Lithocarpus rotundatus
Lithocarpus revolutus
Lithocarpus wrayi
Lithocarpus falconeri
Lithocarpus pycnostachys
Lithocarpus dealbatus
Lithocarpus craibianus
Lithocarpus echinotholus
Lithocarpus braianensis
Lithocarpus laetus
Lithocarpus mairei
Lithocarpus fenestratus
Lithocarpus cleistocarpus
Lithocarpus variolosus
Lithocarpus harlandii
Lithocarpus confinis
Lithocarpus hancei
Lithocarpus haipinii
Lithocarpus henryi
Lithocarpus calolepis
Lithocarpus megalophyllus
Lithocarpus elaeagnifolius
Lithocarpus garrettianus
Lithocarpus glaber
Lithocarpus litseifolius
Lithocarpus taitoensis
Lithocarpus polystachyus
Lithocarpus silvicolarum
Lithocarpus pulcher
Lithocarpus fordianus
Lithocarpus annamensis
Lithocarpus corneus
Castanopsis fargesii
Castanopsis cuspidata
Castanopsis carlesii
Castanopsis echinocarpa
Castanopsis ferox
Castanopsis eyrei
Castanopsis chunii
Castanopsis annamensis
Castanopsis kawakamii
Castanopsis lamontii
Castanopsis ceratacantha
Castanopsis hystrix
Castanopsis crassifolia
Castanopsis faberi
Castanopsis calathiformis
Castanopsis orthacantha
Castanopsis dongchoensis
Castanopsis tibetana
Castanopsis borneensis
Castanopsis jucunda
Castanopsis delavayi
Castanopsis indica
Castanopsis sclerophylla
Castanopsis wallichii
Castanopsis argentea
Castanopsis mekongensis
Castanopsis pierrei
Castanopsis longipetiolata
Castanopsis fulva
Castanopsis philipensis
Castanopsis piriformis
Castanopsis cambodiana
Castanopsis chinensis
Castanopsis fissa
Castanopsis fordii
Castanopsis nephelioides
Castanea crenata
Castanea seguinii
Castanea mollissima
Castanea henryi
Castanea dentata
Castanea ozarkensis
Castanea pumila
Castanea sativa
Trigonobalanus excelsa
Trigonobalanus doichangensis
Fagus sylvatica
Fagus orientalis
Fagus crenata
Fagus lucida
Fagus grandifolia
Fagus japonica
Fagus engleriana
Myrica hartwegii
Morella pringlei
Juglans australis
Carya pallida
Betula pamirica
Betula litwinowii
Carpinus monbeigiana
Nothofagus fusca
1.66667

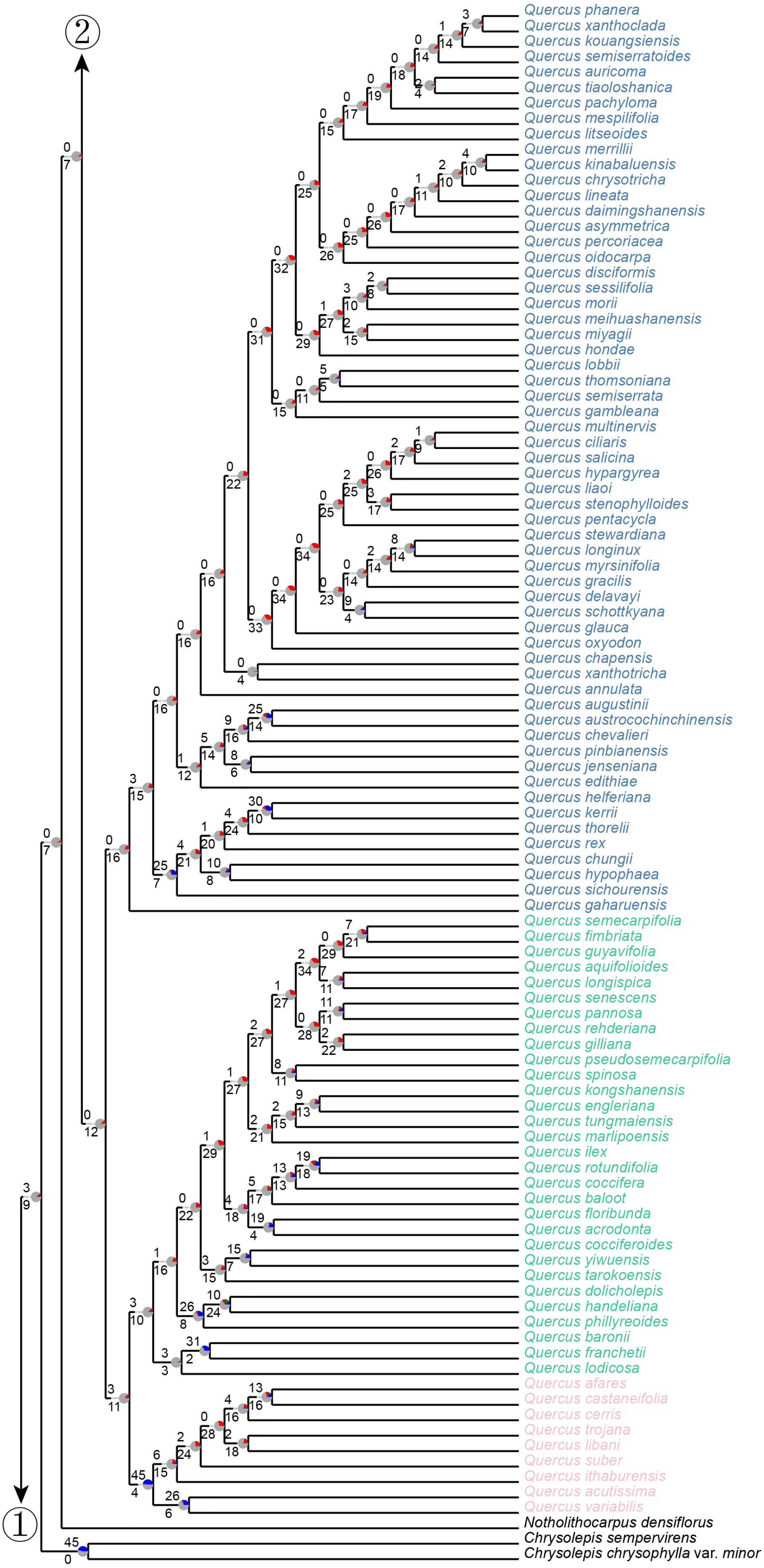

Quercus phanera
Quercus xanthoclada
Quercus kouangsiensis
Quercus semiserratoides
Quercus auricoma
Quercus tiaoloshanica
Quercus pachyloma
Quercus mespilifolia
Quercus litseoides
Quercus merrillii
Quercus kinabaluensis
Quercus chrysotricha
Quercus lineata
Quercus daimingshanensis
Quercus asymmetrica
Quercus percoriacea
Quercus oidocarpa
Quercus disciformis
Quercus sessilifolia
Quercus morii
Quercus meihuashanensis
Quercus miyagii
Quercus hondae
Quercus lobbii
Quercus thomsoniana
Quercus semiserrata
Quercus gambleana
Quercus multinervis
Quercus ciliaris
Quercus salicina
Quercus hypargyrea
Quercus liaoi
Quercus stenophylloides
Quercus pentacycla
Quercus stewardiana
Quercus longinux
Quercus myrsinifolia
Quercus gracilis
Quercus delavayi
Quercus schottkyana
Quercus glauca
Quercus oxyodon
Quercus chapensis
Quercus xanthotricha
Quercus annulata
Quercus augustinii
Quercus austrocochinchinensis
Quercus chevalieri
Quercus pinbianensis
Quercus jenseniana
Quercus edithiae
Quercus helferiana
Quercus kerrii
Quercus thorelii
Quercus rex
Quercus chungii
Quercus hypophaea
Quercus sichourensis
Quercus gaharuensis
Quercus semecarpifolia
Quercus fimbriata
Quercus guyavifolia
Quercus aquifolioides
Quercus longispica
Quercus senescens
Quercus pannosa
Quercus rehderiana
Quercus gilliana
Quercus pseudosemecarpifolia
Quercus spinosa
Quercus kongshanensis
Quercus engleriana
Quercus tungmaiensis
Quercus marlipoensis
Quercus ilex
Quercus rotundifolia
Quercus coccifera
Quercus baloot
Quercus floribunda
Quercus acrodonta
Quercus cocciferoides
Quercus yiwuensis
Quercus tarokoensis
Quercus dolicholepis
Quercus handeliana
Quercus phillyreoides
Quercus baronii
Quercus franchetii
Quercus lodicosa
Quercus afares
Quercus castaneifolia
Quercus cerris
Quercus trojana
Quercus libani
Quercus suber
Quercus ithaburensis
Quercus acutissima
Quercus variabilis
Notholithocarpus densiflorus
Chrysolepis sempervirens
Chrysolepis chrysophylla var. minor
①
②

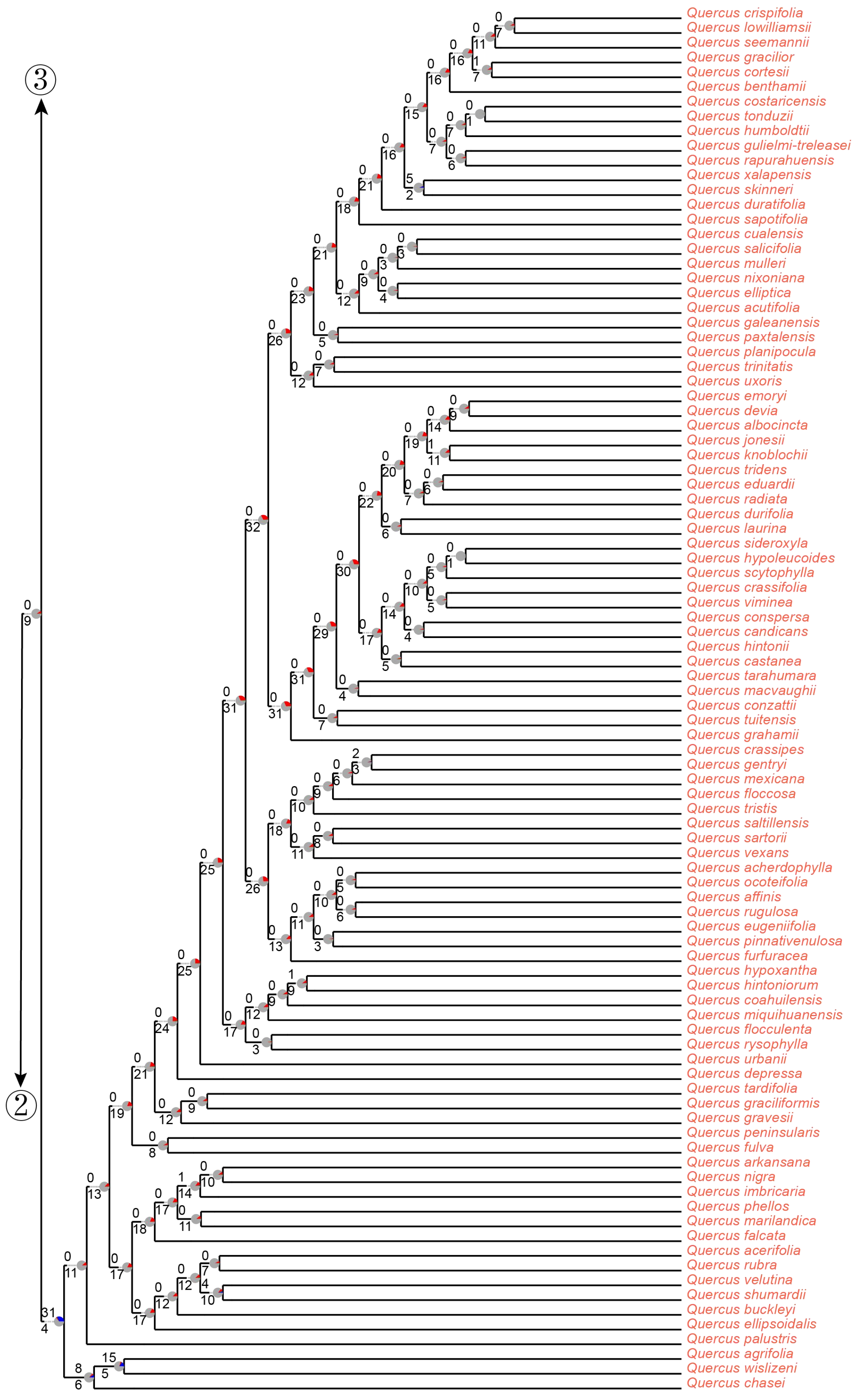

3
2
Quercus crispifolia
Quercus lowilliamsii
Quercus seemannii
Quercus gracilior
Quercus cortesii
Quercus benthamii
Quercus costaricensis
Quercus tonduzii
Quercus humboldtii
Quercus gulielmi-treleasei
Quercus rapurahuensis
Quercus xalapensis
Quercus skinneri
Quercus duratifolia
Quercus sapotifolia
Quercus cualensis
Quercus salicifolia
Quercus mulleri
Quercus nixoniana
Quercus elliptica
Quercus acutifolia
Quercus galeanensis
Quercus paxtalensis
Quercus planipocula
Quercus trinitatis
Quercus uxoris
Quercus emoryi
Quercus devia
Quercus albocincta
Quercus jonesii
Quercus knoblochii
Quercus tridens
Quercus eduardii
Quercus radiata
Quercus durifolia
Quercus laurina
Quercus sideroxyla
Quercus hypoleucoides
Quercus scytophylla
Quercus crassifolia
Quercus viminea
Quercus conspersa
Quercus candicans
Quercus hintonii
Quercus castanea
Quercus tarahumara
Quercus macvaughii
Quercus conzattii
Quercus tuitensis
Quercus grahamii
Quercus crassipes
Quercus gentryi
Quercus mexicana
Quercus floccosa
Quercus tristis
Quercus saltillensis
Quercus sartorii
Quercus vexans
Quercus acherdophylla
Quercus ocoteifolia
Quercus affinis
Quercus rugulosa
Quercus eugeniifolia
Quercus pinnativenulosa
Quercus furfuracea
Quercus hypoxantha
Quercus hintoniorum
Quercus coahuilensis
Quercus miquihuanensis
Quercus flocculenta
Quercus rysophylla
Quercus urbanii
Quercus depressa
Quercus tardifolia
Quercus graciliformis
Quercus gravesii
Quercus peninsularis
Quercus fulva
Quercus arkansana
Quercus nigra
Quercus imbricaria
Quercus phellos
Quercus marilandica
Quercus falcata
Quercus acerifolia
Quercus rubra
Quercus velutina
Quercus shumardii
Quercus buckleyi
Quercus ellipsoidalis
Quercus palustris
Quercus agrifolia
Quercus wislizeni
Quercus chasei

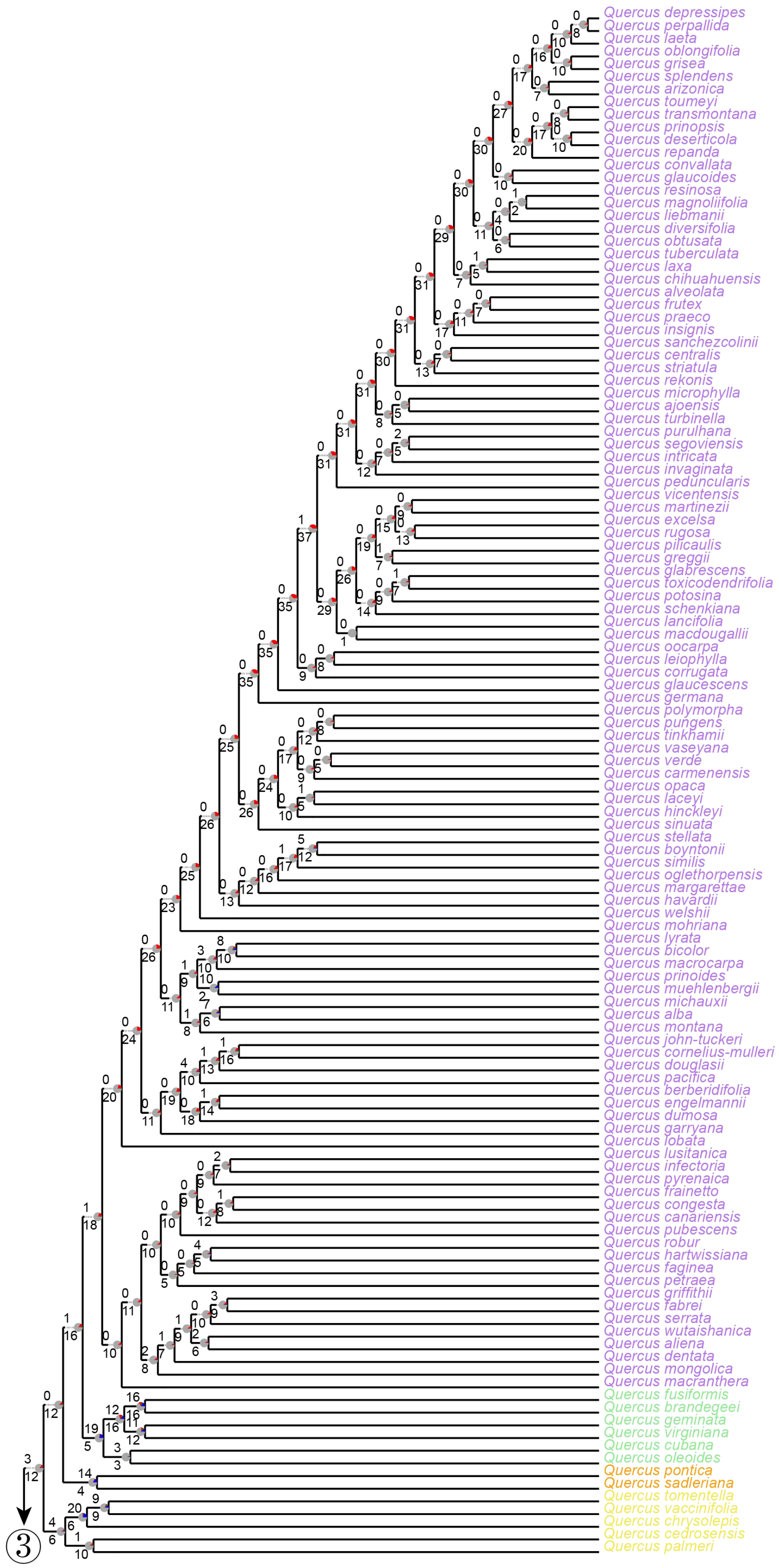
Quercus depressipes
Quercus perpallida
Quercus laeta
Quercus oblongifolia
Quercus grisea
Quercus splendens
Quercus arizonica
Quercus toumeyi
Quercus transmontana
Quercus prinopsis
Quercus deserticola
Quercus repanda
Quercus convallata
Quercus glaucoides
Quercus resinosa
Quercus magnoliifolia
Quercus liebmanii
Quercus diversifolia
Quercus obtusata
Quercus tuberculata
Quercus laxa
Quercus chihuahuensis
Quercus alveolata
Quercus frutex
Quercus praeco
Quercus insignis
Quercus sanchezcolinii
Quercus centralis
Quercus striatula
Quercus rekonis
Quercus microphylla
Quercus ajoensis
Quercus turbinella
Quercus purulhana
Quercus segoviensis
Quercus intricata
Quercus invaginata
Quercus peduncularis
Quercus vicentensis
Quercus martinezii
Quercus excelsa
Quercus rugosa
Quercus pilicaulis
Quercus greggii
Quercus glabrescens
Quercus toxicodendrifolia
Quercus potosina
Quercus schenkiana
Quercus lancifolia
Quercus macdougallii
Quercus oocarpa
Quercus leiophylla
Quercus corrugata
Quercus glaucescens
Quercus germana
Quercus polymorpha
Quercus pungens
Quercus tinkhamii
Quercus vaseyana
Quercus verde
Quercus carmenensis
Quercus opaca
Quercus laceyi
Quercus hinckleyi
Quercus sinuata
Quercus stellata
Quercus boyntonii
Quercus similis
Quercus oglethorpensis
Quercus margarettae
Quercus havardii
Quercus welshii
Quercus mohriana
Quercus lyrata
Quercus bicolor
Quercus macrocarpa
Quercus prinoides
Quercus muehlenbergii
Quercus michauxii
Quercus alba
Quercus montana
Quercus john-tuckeri
Quercus cornelius-mulleri
Quercus douglasii
Quercus pacifica
Quercus berberidifolia
Quercus engelmannii
Quercus dumosa
Quercus garryana
Quercus lobata
Quercus lusitanica
Quercus infectoria
Quercus pyrenaica
Quercus frainetto
Quercus congesta
Quercus canariensis
Quercus pubescens
Quercus robur
Quercus hartwissiana
Quercus faginea
Quercus petraea
Quercus griffithii
Quercus fabrei
Quercus serrata
Quercus wutaishanica
Quercus aliena
Quercus dentata
Quercus mongolica
Quercus macranthera
Quercus fusiformis
Quercus brandegeei
Quercus geminata
Quercus virginiana
Quercus cubana
Quercus oleoides
Quercus pontica
Quercus sadleriana
Quercus tomentella
Quercus vaccinifolia
Quercus chrysolepis
Quercus cedrosensis
Quercus palmeri
3

**FIGURE S18. Phyparts results based on the 98 gene trees from the HYB-98RT dataset, mapped against the ASTRAL species tree.** Bootstrap contraction threshold for phyparts analysis is set as 70%. Numbers next to branches represent the number of concordant (above) and conflicting (below) gene trees with that clade. The color scheme of pie charts next to nodes is the same as in Supplementary Figure S17. Tip labels are colored by the same color scheme as in Supplementary Figure S4.

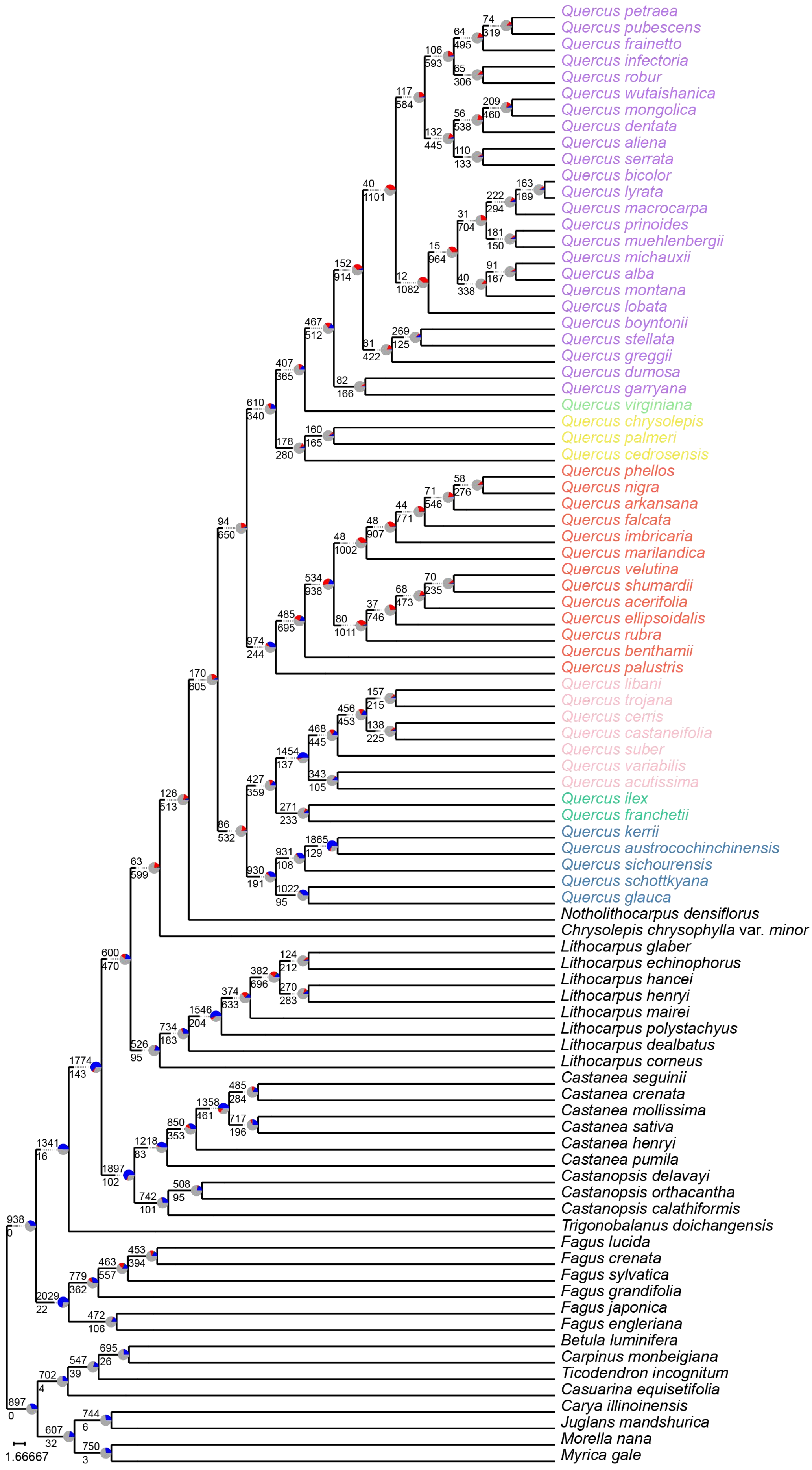
Quercus petraea
Quercus pubescens
Quercus frainetto
Quercus infectoria
Quercus robur
Quercus wutaishanica
Quercus mongolica
Quercus dentata
Quercus aliena
Quercus serrata
Quercus bicolor
Quercus lyrata
Quercus macrocarpa
Quercus prinoides
Quercus muehlenbergii
Quercus michauxii
Quercus alba
Quercus montana
Quercus lobata
Quercus boyntonii
Quercus stellata
Quercus greggii
Quercus dumosa
Quercus garryana
Quercus virginiana
Quercus chrysolepis
Quercus palmeri
Quercus cedrosensis
Quercus phellos
Quercus nigra
Quercus arkansana
Quercus falcata
Quercus imbricaria
Quercus marilandica
Quercus velutina
Quercus shumardii
Quercus acerifolia
Quercus ellipsoidalis
Quercus rubra
Quercus benthamii
Quercus palustris
Quercus libani
Quercus trojana
Quercus cerris
Quercus castaneifolia
Quercus suber
Quercus variabilis
Quercus acutissima
Quercus ilex
Quercus franchetii
Quercus kerrii
Quercus austrocochinchinensis
Quercus sichourensis
Quercus schottkyana
Quercus glauca
Notholithocarpus densiflorus
Chrysolepis chrysophylla var. minor
Lithocarpus glaber
Lithocarpus echinophorus
Lithocarpus hancei
Lithocarpus henryi
Lithocarpus mairei
Lithocarpus polystachyus
Lithocarpus dealbatus
Lithocarpus corneus
Castanea seguinii
Castanea crenata
Castanea mollissima
Castanea sativa
Castanea henryi
Castanea pumila
Castanopsis delavayi
Castanopsis orthacantha
Castanopsis calathiformis
Trigonobalanus doichangensis
Fagus lucida
Fagus crenata
Fagus sylvatica
Fagus grandifolia
Fagus japonica
Fagus engleriana
Betula luminifera
Carpinus monbeigiana
Ticodendron incognitum
Casuarina equisetifolia
Carya illinoinensis
Juglans mandshurica
Morella nana
Myrica gale
1.66667

**FIGURE S19. Phyparts results based on the 2821 gene trees in the RNA-2821RT dataset, mapped against the ASTRAL species tree.** Bootstrap contraction threshold for phyparts analysis is set as 70%. The color scheme of pie charts next to nodes is the same as in Supplementary Figure S17. Tip labels are colored by the same color scheme as in Supplementary Figure S4.

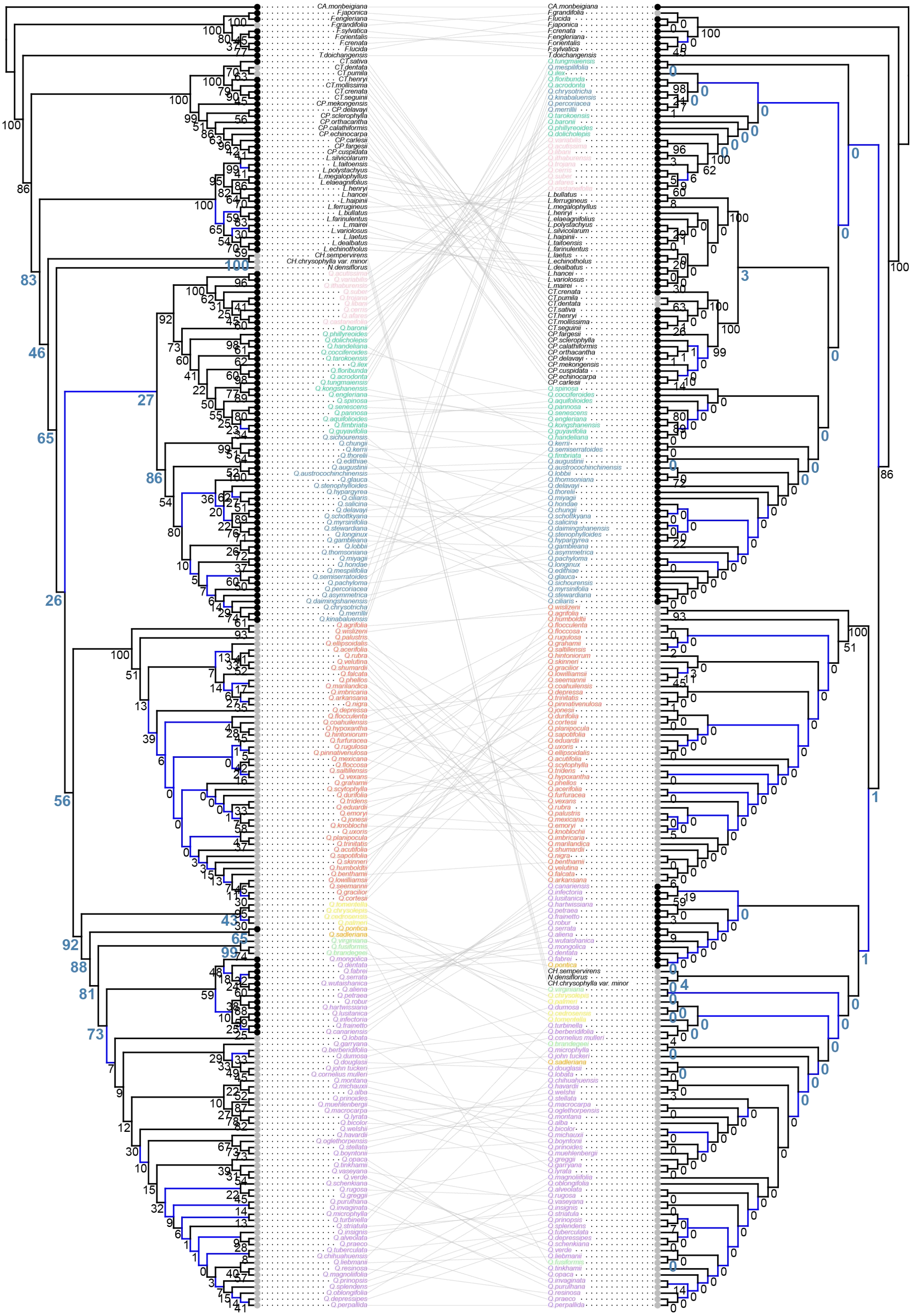

**FIGURE S20. Cophylogeny showing incongruence between the nuclear ASTRAL (left; the HYB-98RT dataset) and unpartitioned chloroplast ML (right) trees with coalescent simulation results from the nuclear guide tree scaled by a factor**

**of four.** Branches with low support value (LPP < 0.5; BS < 50%) are colored with blue. Clade frequencies from the gene trees simulated under the coalescent are shown near nodes; clade frequencies associated with deep cytonuclear discordances are blue, bold, and enlarged. Black dots at the tips denote an Old World distribution, while gray dots denote a New World distribution. Tip labels are colored by the same color scheme as in Supplementary Figure S4. This plot was generated using the function "cophylo()" in the R package phytools (Revell 2012). Abbreviations: CA., *Carpinus*; F., *Fagus*; T., *Trigonobalanus*; CP., *Castanopsis*; CT., *Castanea*; L., *Lithocarpus*; CH., *Chrysolepis*; N., *Notholithocarpus*; Q., *Quercus*.

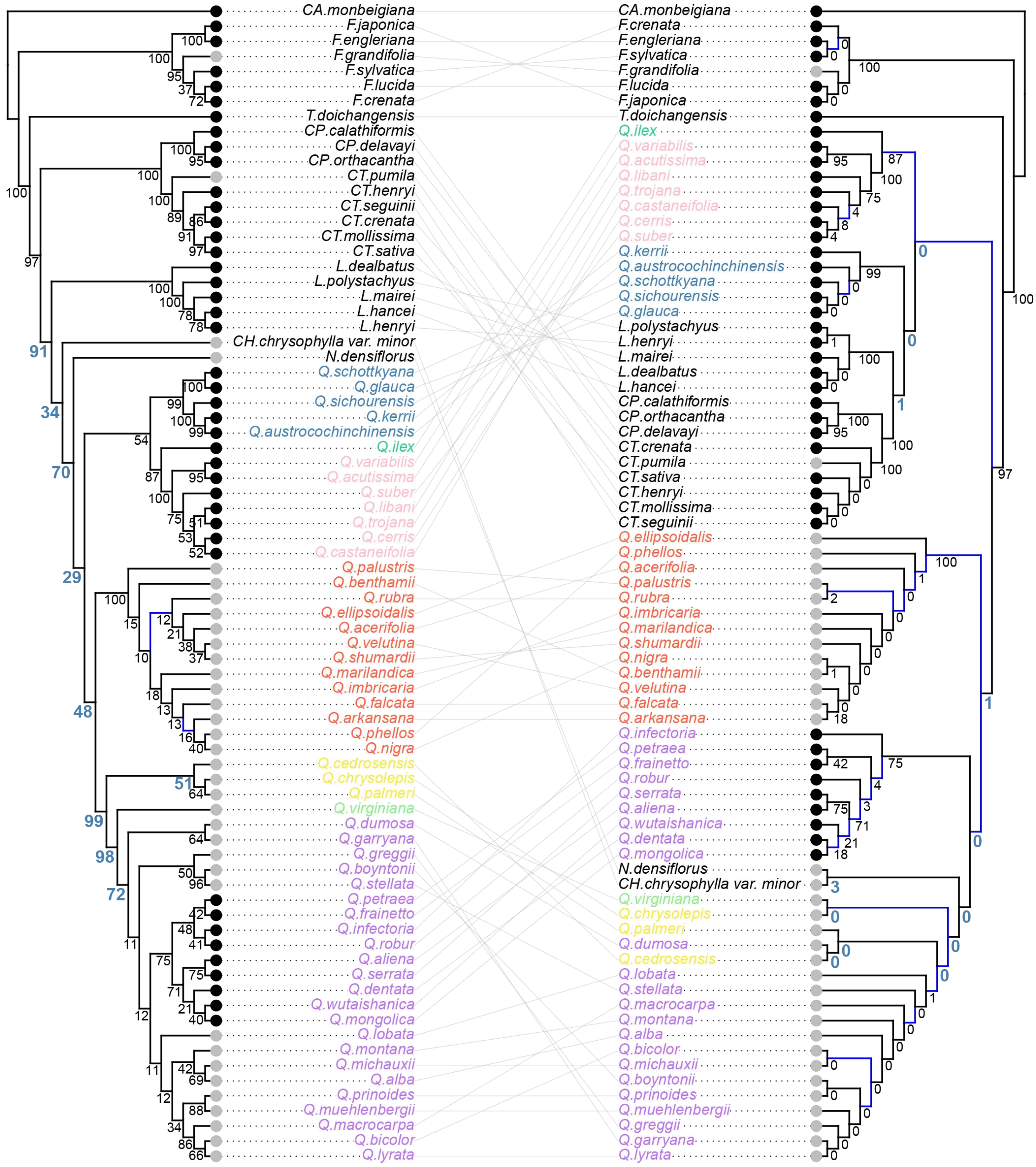


**FIGURE S21. Tanglegram comparing the nuclear ASTRAL (left; the RNA-2821RT dataset) and chloroplast ML (right) trees optimized in Dendroscope, with coalescent simulation results from the nuclear guide tree scaled by a factor of four.** The two trees shown here with 77 tips are pruned from the ASTRAL tree of the RNA-2821RT dataset (see Supplementary Fig. S9 for tree with 89 tips) and the ML tree of plastome dataset inferred with an unpartitioned model (see Supplementary Fig. S27 for tree with 223 tips). Branches with low support value (LPP

< 0.5; BS < 50%) are colored with blue. Clade frequencies from the gene trees simulated under the coalescent are shown near nodes; clade frequencies associated with deep cytonuclear discordances are blue, bold, and enlarged. Black dots at the tips denote an Old World distribution, while gray dots denote a New World distribution. Tip labels are colored by the same color scheme as in Supplementary Figure S4. Abbreviations: CA., *Carpinus*; F., *Fagus*; T., *Trigonobalanus*; CP., *Castanopsis*; CT., *Castanea*; L., *Lithocarpus*; CH., *Chrysolepis*; N., *Notholithocarpus*; Q., *Quercus*.

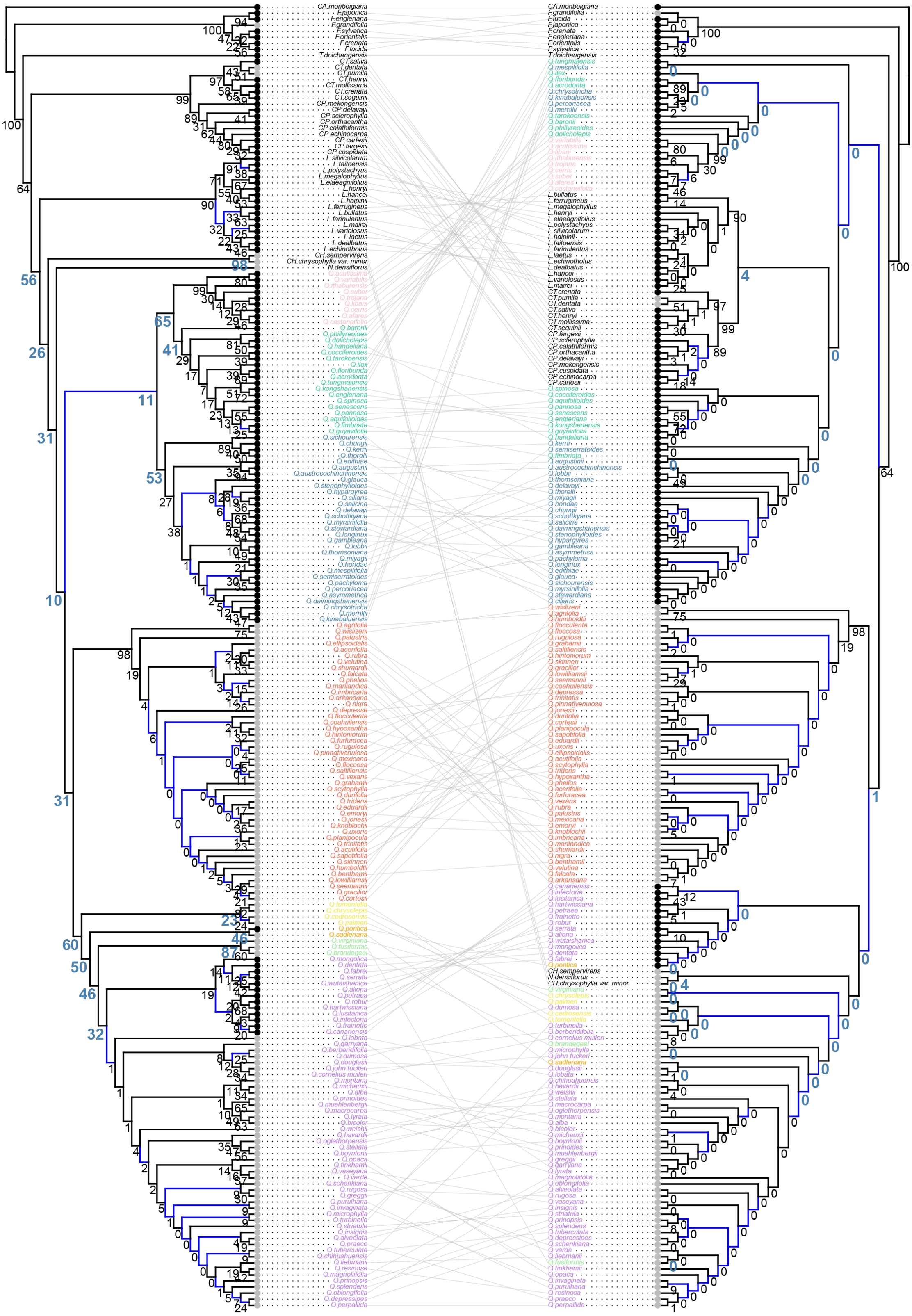

**FIGURE S22. Cophylogeny showing incongruence between the nuclear ASTRAL (left; the HYB-98RT dataset) and unpartitioned chloroplast ML (right) trees with coalescent simulation results from the nuclear guide tree scaled by a factor**

**of two.** The nuclear and chloroplast trees shown here, each with 215 tips, are pruned from the ASTRAL tree of the HYB-98RT dataset and the concatenated ML tree of the plastome dataset under the unpartitioned model, respectively. The complete nuclear tree with 431 tips and chloroplast tree with 223 tips are provided in Supplementary Figures S5 and S27, respectively. Branches with low support value ($LPP < 0.5$; $BS < 50\%$) are colored with blue. Clade probabilities from the gene tree distribution predicted under the coalescent are shown near nodes; clade frequencies associated with deep cytonuclear discordances are blue, bold, and enlarged. Black dots at the tips denote an Old World distribution, while gray dots denote a New World distribution. Tip labels are colored by the same color scheme as in Supplementary Figure S4. Abbreviations: CA., *Carpinus*; F., *Fagus*; T., *Trigonobalanus*; CP., *Castanopsis*; CT., *Castanea*; L., *Lithocarpus*; CH., *Chrysolepis*; N., *Notholithocarpus*; Q., *Quercus*.

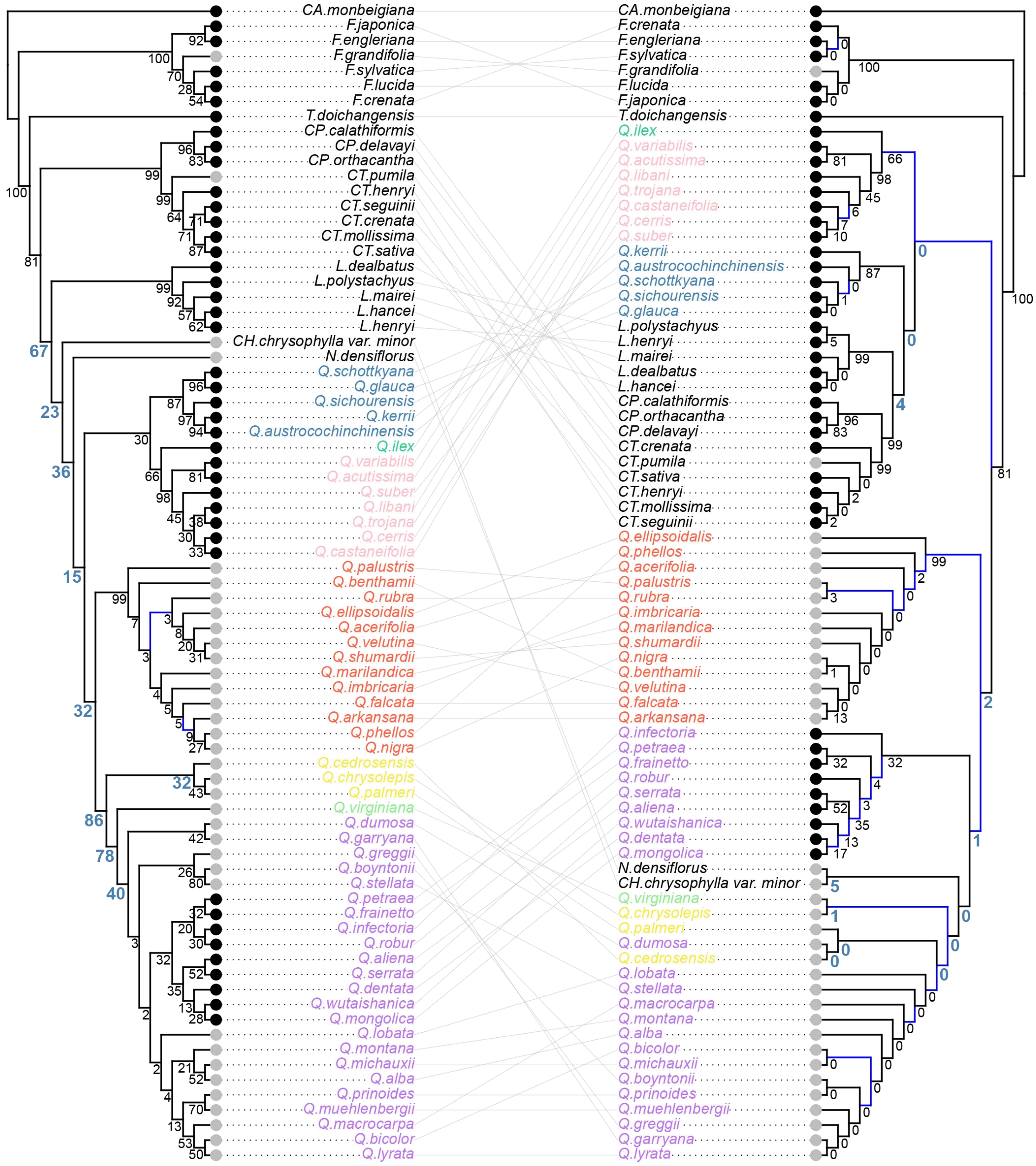


**FIGURE S23. Cophylogeny showing incongruence between the nuclear ASTRAL (left; the RNA-2821RT dataset) and unpartitioned chloroplast ML (right) trees with coalescent simulation results from the nuclear guide tree scaled by a factor of two.** The nuclear and chloroplast trees shown here, each with 77 tips, are pruned from the ASTRAL tree of the RNA-2821RT dataset and the concatenated ML tree of the plastome dataset under the unpartitioned model, respectively. The complete nuclear tree with 89 tips and chloroplast tree with 223 tips are provided in

Supplementary Figures S9 and S27, respectively. Branches with low support value (LPP < 0.5; BS < 50%) are colored with blue. Clade probabilities from the gene tree distribution predicted under the coalescent are shown near nodes; clade frequencies associated with deep cytonuclear discordances are blue, bold, and enlarged. Black dots at the tips denote an Old World distribution, while gray dots denote a New World distribution. Tip labels are colored by the same color scheme as in Supplementary Figure S4. Abbreviations: CA., *Carpinus*; F., *Fagus*; T., *Trigonobalanus*; CP., *Castanopsis*; CT., *Castanea*; L., *Lithocarpus*; CH., *Chrysolepis*; N., *Notholithocarpus*; Q., *Quercus*.

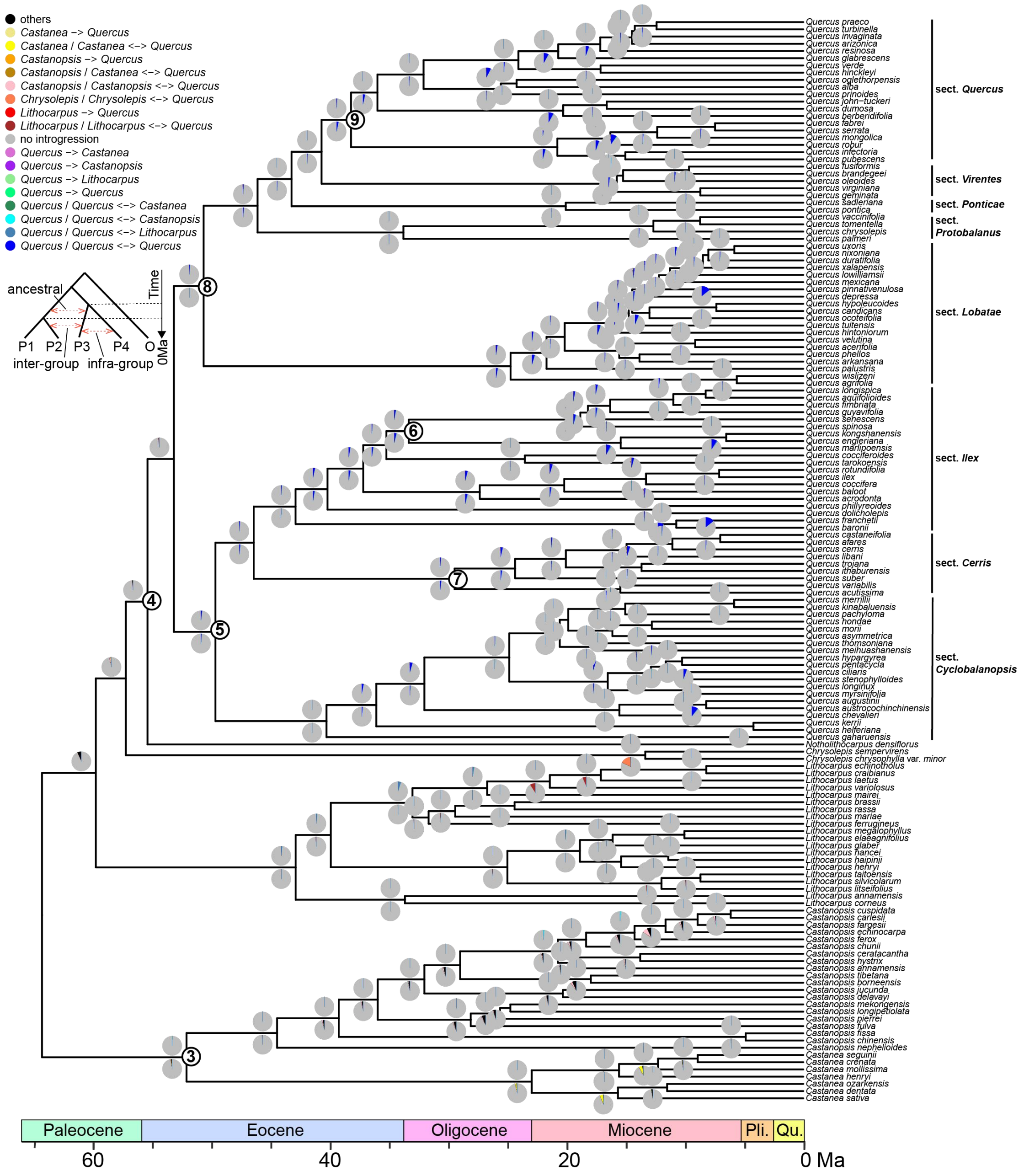


**FIGURE S24. Proportions of positive tests for various introgression signatures visualized on the time-calibrated tree based on the HYB-98RT dataset.** Results here are from a reduced sample of 150 taxa from the HYB-98RT dataset. Signatures of introgression involving oaks are shown in color (according to the key) while the introgression signatures between two non-*Quercus* taxa are shown in black (see Supplementary Table S7 for detailed results). The five-taxon tree in the top left corner shows the four ingroup taxa (P1–P4) and the outgroup (O) used for the $D_{\mathrm{FOIL}}$ test, in which two introgression types can be detected: (1) “ancestral” introgression between the ancestral taxon of one subgroup (comprising P1 and P2) and P3 (or P4) without considering direction (i.e., P1/P2<–>P3 or P1/P2<–>P4), and (2) “inter-group” introgression between taxa in different subgroups with determined direction (e.g., P1–>P3 represents introgression from taxon P1 to taxon P3). Pie charts above the nodes represent the proportions of positive tests for various introgression signatures (shown by the legend) when that node was the most recent common ancestor (MRCA) of taxa P3 and P4, while pie charts below the nodes represent the results when that node was the MRCA of taxa P1 and P2. The numbers in circles show the placements of the fossil calibrations used (see Supplementary Table S6 for details).

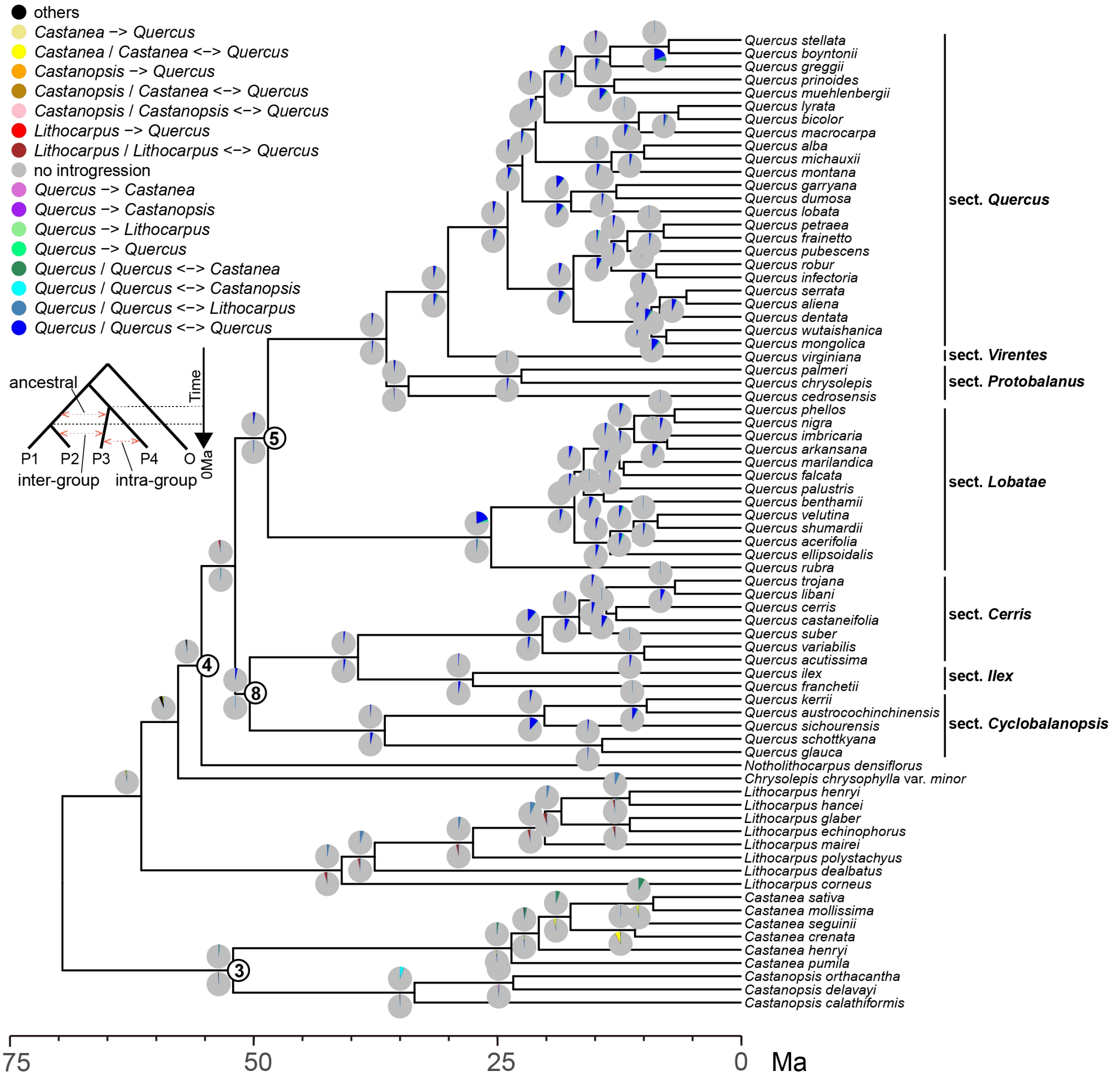


**FIGURE S25. Proportions of positive tests for various introgression signatures visualized on the time-calibrated tree based on the RNA-2821RT dataset.** The numbers in circles show the placements of the fossil calibrations used (see Supplementary Table S6 for details).

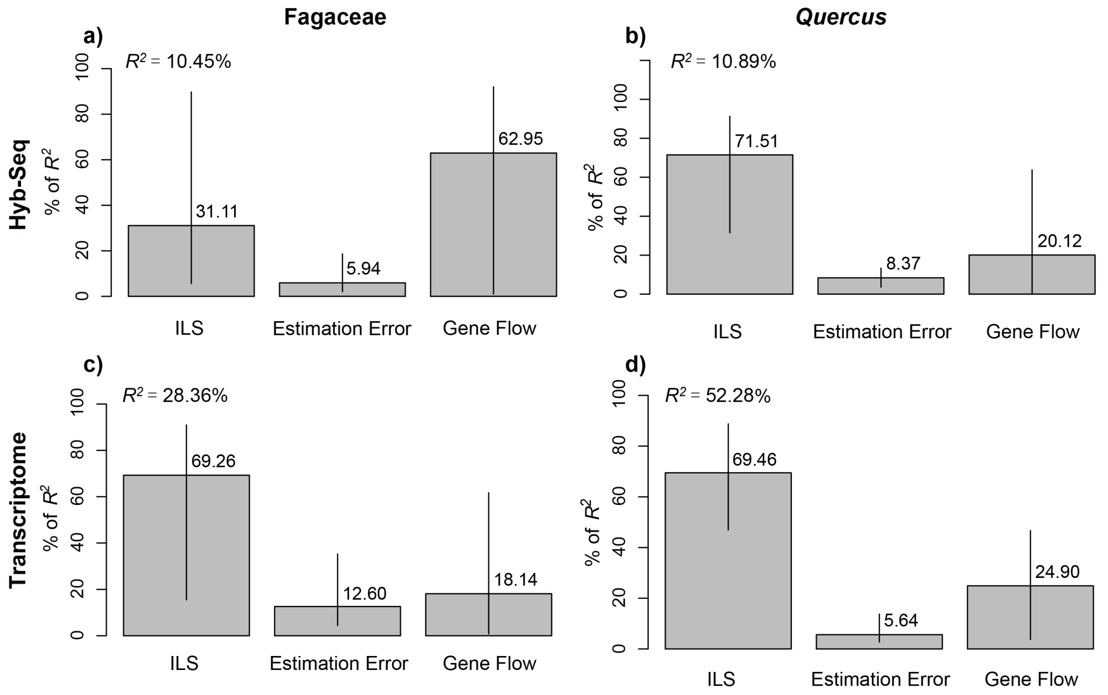


**FIGURE S26. Plots showing the relative contributions of incomplete lineage sorting (ILS), gene flow, and gene tree estimation error to the observed gene-tree discordance, across two datasets and two taxonomic levels. a)** Fagaceae in the Hyb-Seq dataset (represented here by the HYB-98RT dataset). **b)** *Quercus* in the Hyb-Seq dataset. **c)** Fagaceae in the transcriptome dataset (represented here by the RNA-2821RT dataset). **d)** *Quercus* in the transcriptome dataset. $R^2$ denotes the total proportion of gene tree variation explained by the model. Relative importance with 95% bootstrap confidence intervals are decomposed with the “lmg” method and regressors of log transformations and summed to 100%.

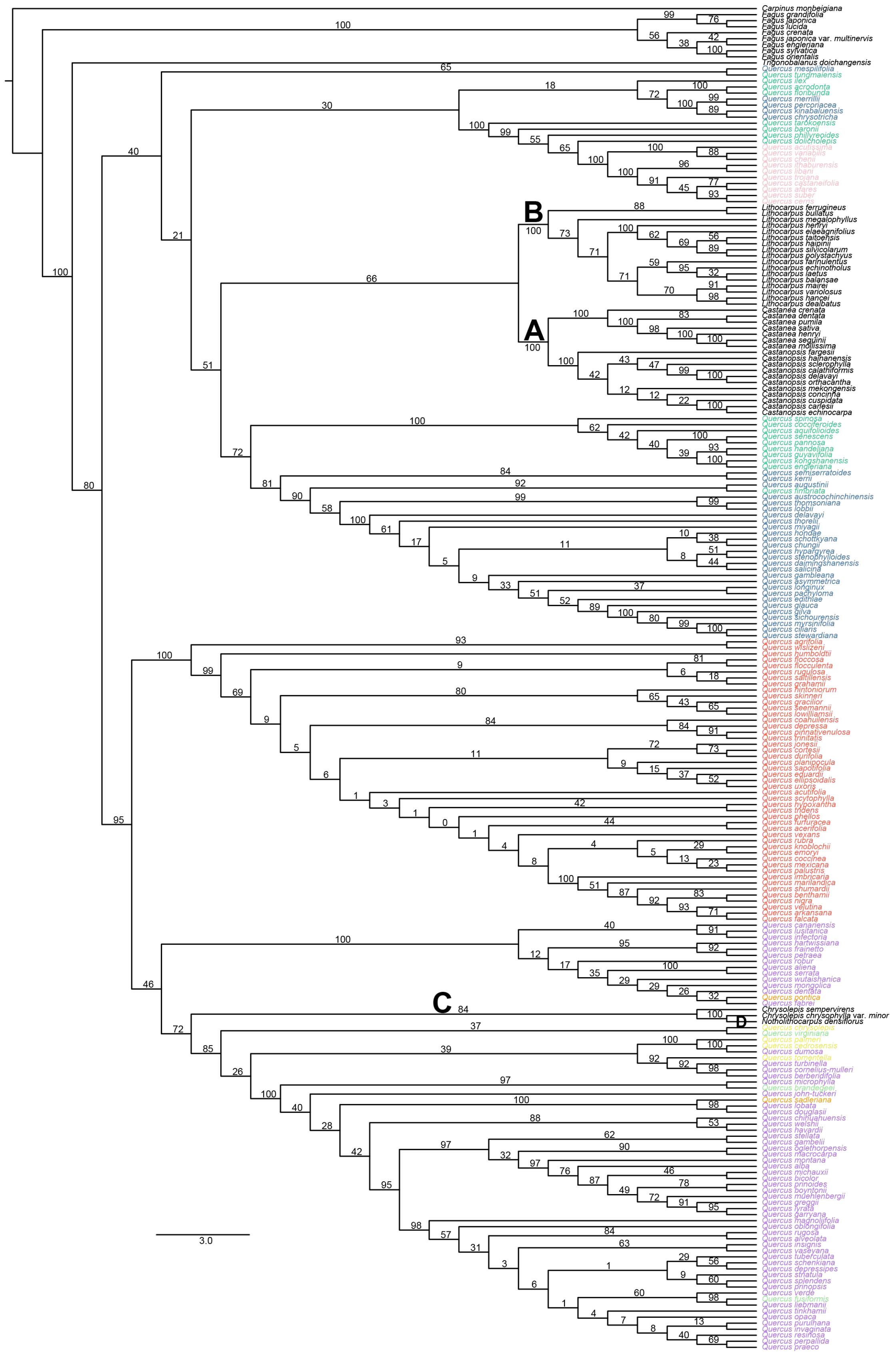

**FIGURE S27. Cladogram of the chloroplast ML tree of oaks and relatives inferred by RAxML based on the concatenated plastome supermatrix including**

**223 species under an unpartitioned GTR-GAMMA model.** Bootstrap values are shown above branches. A–D denote the deep uncertain branches (Supplementary Table S5) investigated in this study. Tip labels are colored following the color scheme in Supplementary Figure S4.

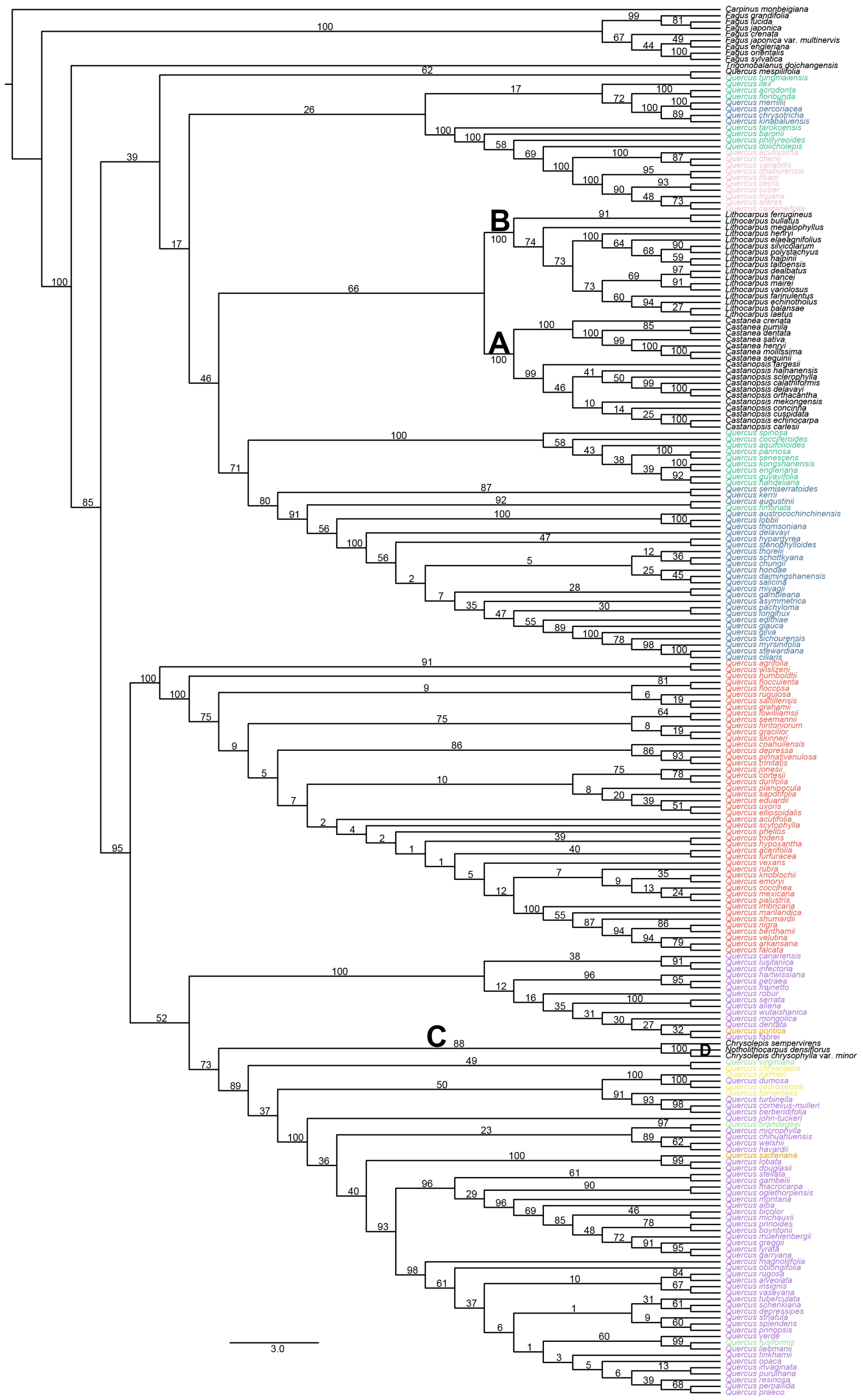

Carpinus monbeigiana
Fagus grandifolia
Fagus lucida
Fagus japonica
Fagus crenata
Fagus japonica var. multinervis
Fagus engleriana
Fagus orientalis
Fagus sylvatica
Trigonobalanus doichangensis
Quercus mespilifolia
Quercus tungmaiensis
Quercus ilex
Quercus acrodonta
Quercus floribunda
Quercus merrillii
Quercus percoriacea
Quercus chrysotricha
Quercus kinabaluensis
Quercus tarokoensis
Quercus baronii
Quercus phillyreoides
Quercus dolicholepis
Quercus acutissima
Quercus chenii
Quercus variabilis
Quercus ithaburensis
Quercus libani
Quercus cerris
Quercus suber
Quercus trojana
Quercus afares
Quercus castaneifolia
B
Lithocarpus ferrugineus
Lithocarpus bullatus
Lithocarpus megalophyllus
Lithocarpus henryi
Lithocarpus elaeagnifolius
Lithocarpus silvicolarum
Lithocarpus polystachyus
Lithocarpus haipinii
Lithocarpus taitoensis
Lithocarpus dealbatus
Lithocarpus hancei
Lithocarpus mairei
Lithocarpus variolosus
Lithocarpus farinulentus
Lithocarpus echinotholus
Lithocarpus balansae
Lithocarpus laetus
A
Castanea crenata
Castanea pumila
Castanea dentata
Castanea sativa
Castanea henryi
Castanea mollissima
Castanea seguinii
Castanopsis fargesii
Castanopsis hainanensis
Castanopsis sclerophylla
Castanopsis calathiformis
Castanopsis delavayi
Castanopsis orthacantha
Castanopsis mekongensis
Castanopsis concinna
Castanopsis cuspidata
Castanopsis echinocarpa
Castanopsis carlesii
Quercus spinosa
Quercus cocciferoides
Quercus aquifolioides
Quercus pannosa
Quercus senescens
Quercus kongshanensis
Quercus engleriana
Quercus guyavifolia
Quercus handeliana
Quercus semiserratoides
Quercus kerrii
Quercus augustinii
Quercus fimbriata
Quercus austrocochinchinensis
Quercus lobbii
Quercus thomsoniana
Quercus delavayi
Quercus hypargyrea
Quercus stenophylloides
Quercus thorelii
Quercus schottkyana
Quercus chungii
Quercus hondae
Quercus daimingshanensis
Quercus salicina
Quercus miyagii
Quercus gambleana
Quercus asymmetrica
Quercus pachyloma
Quercus longinux
Quercus edithiae
Quercus glauca
Quercus gilva
Quercus sichourensis
Quercus myrsinifolia
Quercus stewardiana
Quercus ciliaris
Quercus agrifolia
Quercus wislizeni
Quercus humboldtii
Quercus flocculenta
Quercus floccosa
Quercus rugulosa
Quercus saltillensis
Quercus grahamii
Quercus lowilliamsii
Quercus seemannii
Quercus hintoniorum
Quercus gracilior
Quercus skinneri
Quercus coahuilensis
Quercus depressa
Quercus pinnativenulosa
Quercus trinitatis
Quercus jonesii
Quercus cortesii
Quercus durifolia
Quercus planipocula
Quercus sapotifolia
Quercus eduardii
Quercus uxoris
Quercus ellipsoidalis
Quercus acutifolia
Quercus scytophylla
Quercus phellos
Quercus tridens
Quercus hypoxantha
Quercus acerifolia
Quercus furfuracea
Quercus vexans
Quercus rubra
Quercus knoblochii
Quercus emoryi
Quercus coccinea
Quercus mexicana
Quercus palustris
Quercus imbricaria
Quercus marilandica
Quercus shumardii
Quercus nigra
Quercus benthamii
Quercus velutina
Quercus arkansana
Quercus falcata
Quercus canariensis
Quercus lusitanica
Quercus infectoria
Quercus hartwissiana
Quercus petraea
Quercus frainetto
Quercus robur
Quercus serrata
Quercus aliena
Quercus wutaishanica
Quercus mongolica
Quercus dentata
Quercus pontica
Quercus fabrei
C
D
Chrysolepis sempervirens
Notholithocarpus densiflorus
Chrysolepis chrysophylla var. minor
Quercus virginiana
Quercus chrysolepis
Quercus palmeri
Quercus dumosa
Quercus cedrosensis
Quercus tomentella
Quercus turbinella
Quercus cornelius-mulleri
Quercus berberidifolia
Quercus john-tuckeri
Quercus brandegeei
Quercus microphylla
Quercus chihuahuensis
Quercus welshii
Quercus havardii
Quercus sadleriana
Quercus lobata
Quercus douglasii
Quercus stellata
Quercus gambelii
Quercus macrocarpa
Quercus oglethorpensis
Quercus montana
Quercus alba
Quercus bicolor
Quercus michauxii
Quercus prinoides
Quercus boyntonii
Quercus muehlenbergii
Quercus greggii
Quercus lyrata
Quercus garryana
Quercus magnoliifolia
Quercus oblongifolia
Quercus rugosa
Quercus alveolata
Quercus insignis
Quercus vaseyana
Quercus tuberculata
Quercus schenkiana
Quercus depressipes
Quercus striatula
Quercus splendens
Quercus prinopsis
Quercus verde
Quercus fusiformis
Quercus liebmanii
Quercus tinkhamii
Quercus opaca
Quercus invaginata
Quercus purulhana
Quercus resinosa
Quercus perpallida
Quercus praeco
3.0

**FIGURE S28. Cladogram of the chloroplast ML tree of oaks and relatives inferred by RAxML based on the concatenated plastome supermatrix including 223 species under a GTR-GAMMA model partitioned by genome region (LSC, SSC, and IR).** Bootstrap values are shown above branches. A–D denote the deep uncertain branches (Supplementary Table S5) investigated in this study. Tip labels are colored by the same color scheme as in Supplementary Figure S4.

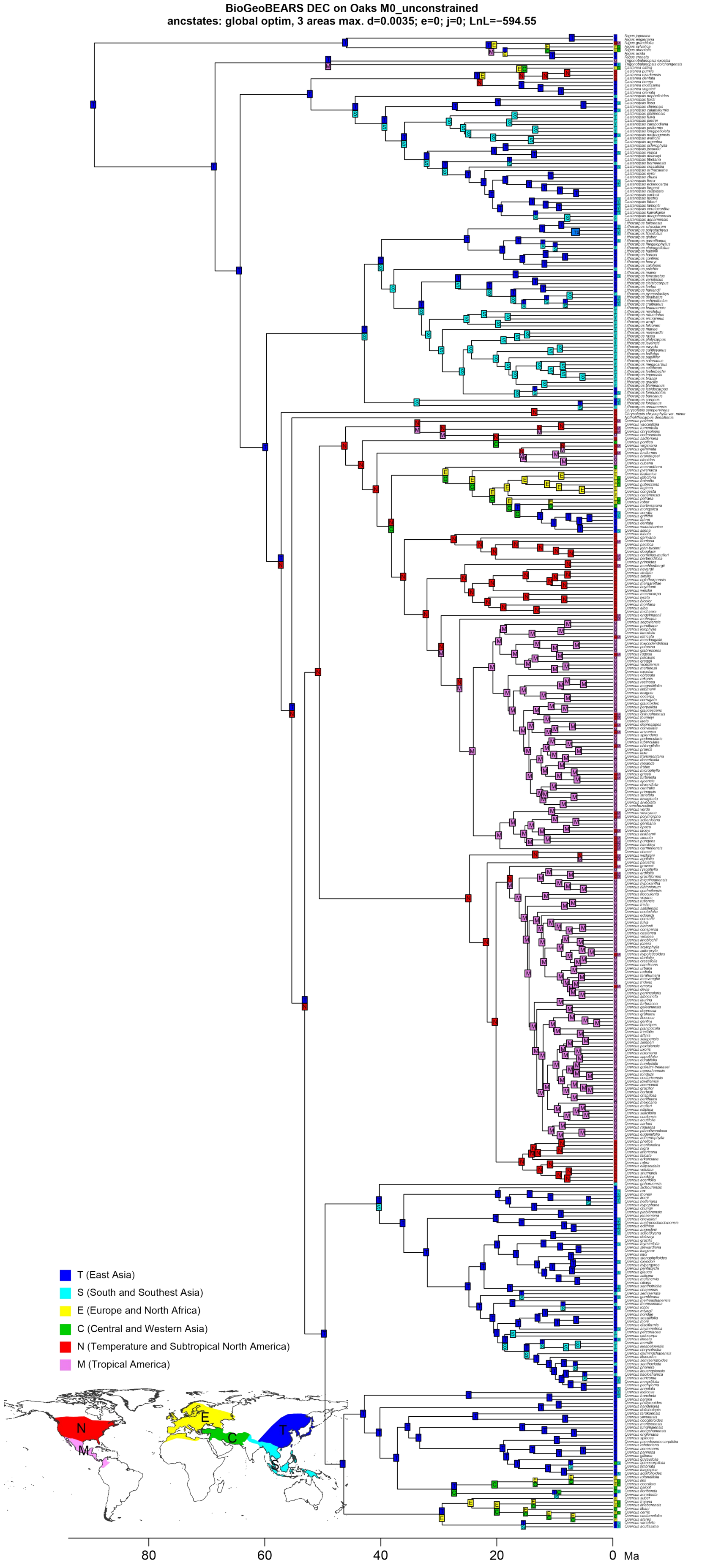
BioGeoBEARS DEC on Oaks M0_unconstrained
ancstates: global optim, 3 areas max. d=0.0035; e=0; j=0; LnL=−594.55
T (East Asia)
S (South and Southeast Asia)
E (Europe and North Africa)
C (Central and Western Asia)
N (Temperature and Subtropical North America)
M (Tropical America)
80
60
40
20
0
Ma

389

**FIGURE S29. Ancestral range estimation of Fagaceae under the DEC model, based on the MCC tree of the HYB-MO89 dataset.**

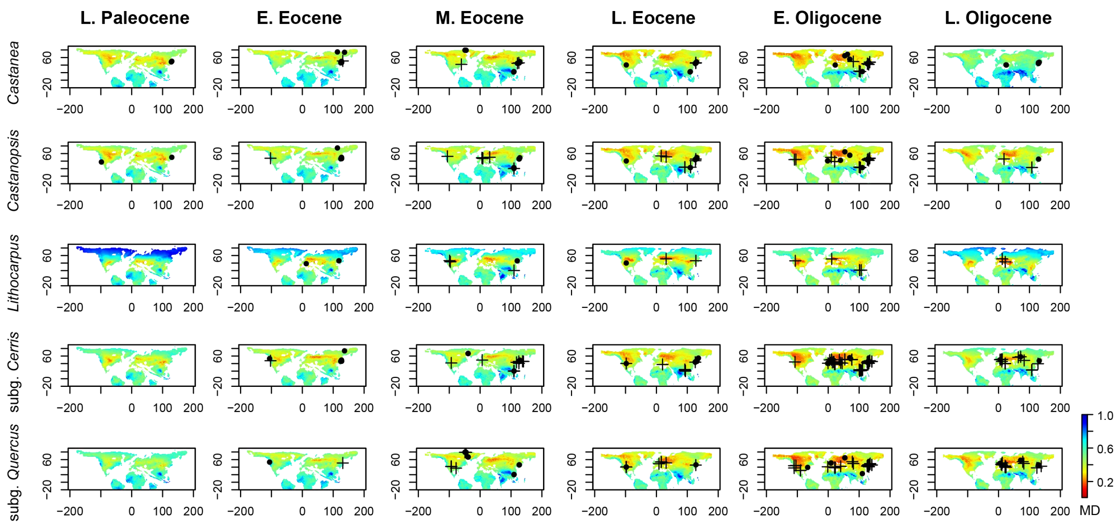


**FIGURE S30. Potential distribution of the major lineages of *Quercus* and Quercoideae from Late Paleocene to Late Eocene as inferred by fossil-based ecological niche modeling under PEO strategy.** These maps were generated by projecting the climatic tolerances of Paleocene, Eocene, and Oligocene fossil taxa (i.e., PEO strategy) onto six paleo-climate scenarios. The fossil distribution of *Chrysolepis* and *Notholithocarpus* during the Paleogene is provided in Supplementary Figure S31. A MD (mahalanobis distance) score of < 0.3 (red) corresponds with highly suitable region for each Quercoideae lineages and otherwise climatically unsuitable region. Dots and crosses denote the location of pollen and macrofossil taxa, respectively. Abbreviations: E., Early; M., Middle; L., Late.

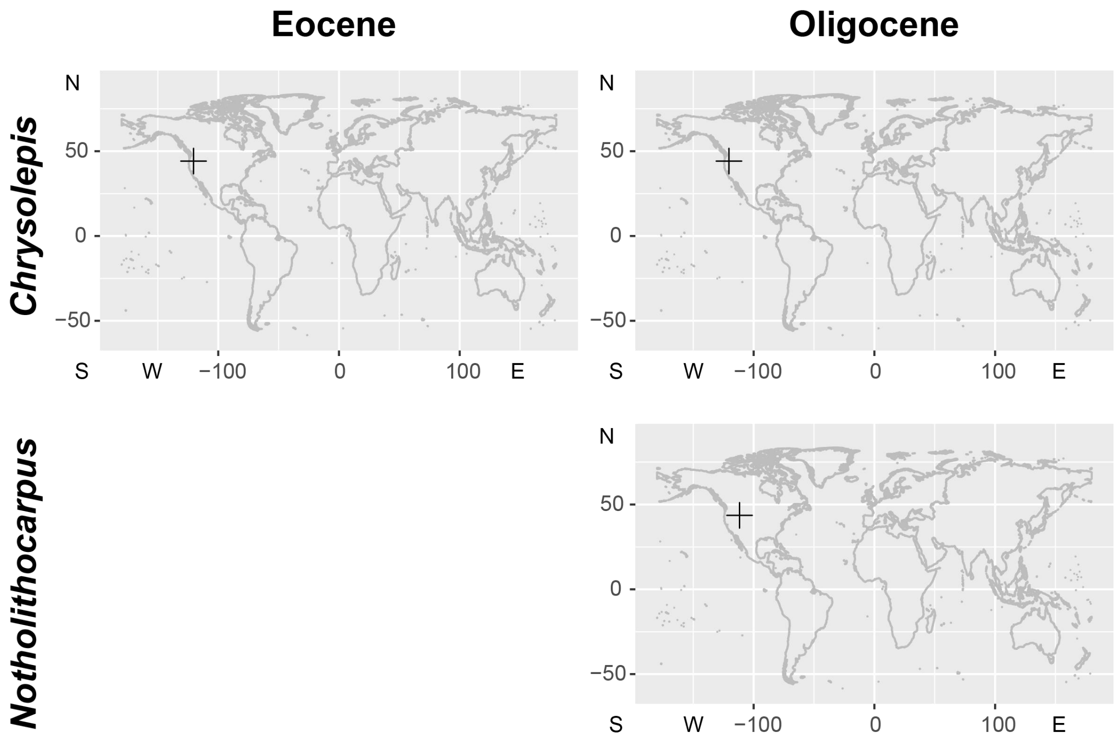


**FIGURE S31. The fossil distribution of *Chrysolepis* and *Notholithocarpus* during the Paleogene.** The cross mark denotes macrofossil taxon.

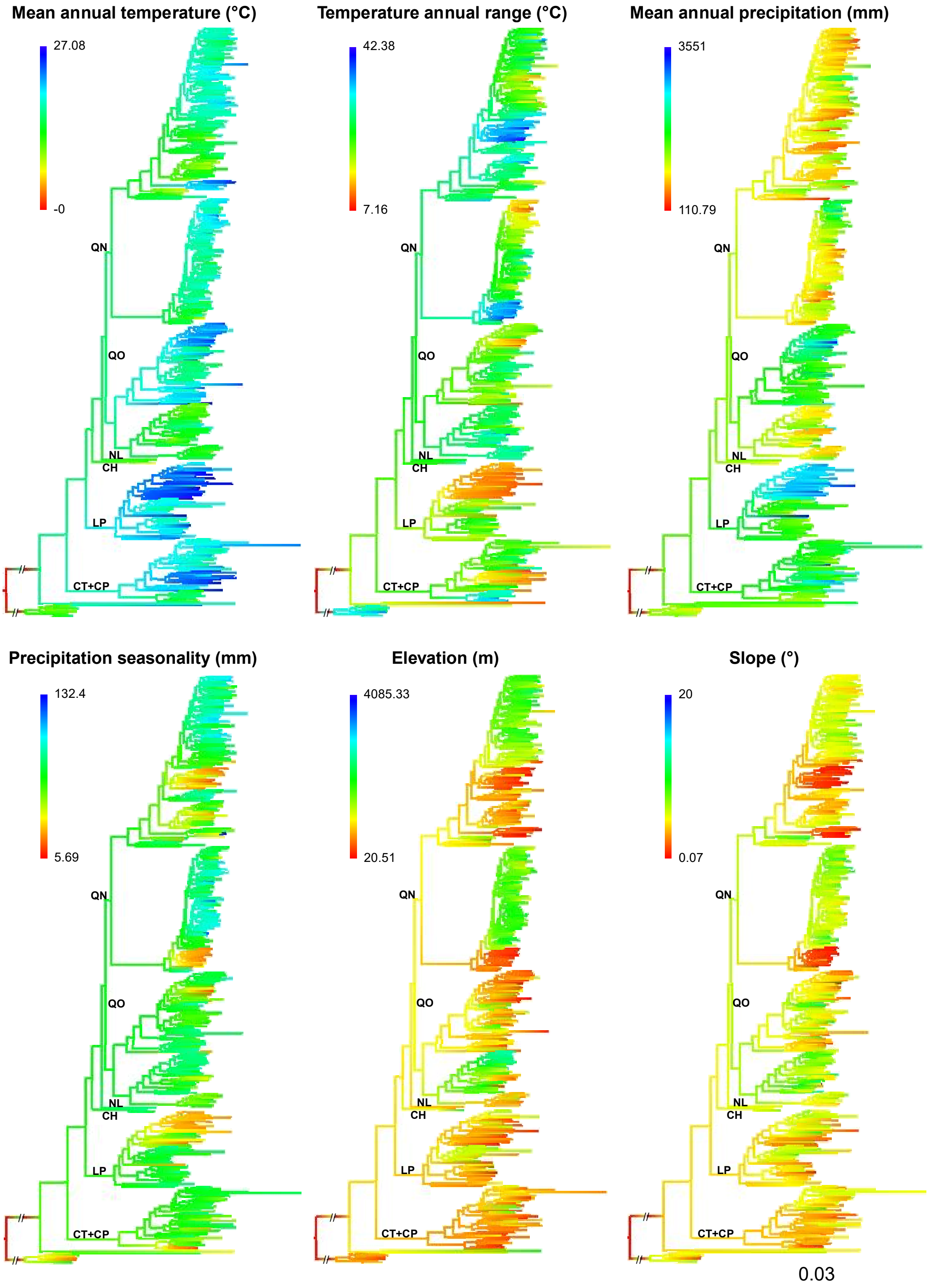
Mean annual temperature (°C)
27.08
-0
Temperature annual range (°C)
42.38
7.16
Mean annual precipitation (mm)
3551
110.79
Precipitation seasonality (mm)
132.4
5.69
Elevation (m)
4085.33
20.51
Slope (°)
20
0.07
QN
QO
NL
CH
LP
CT+CP
0.03

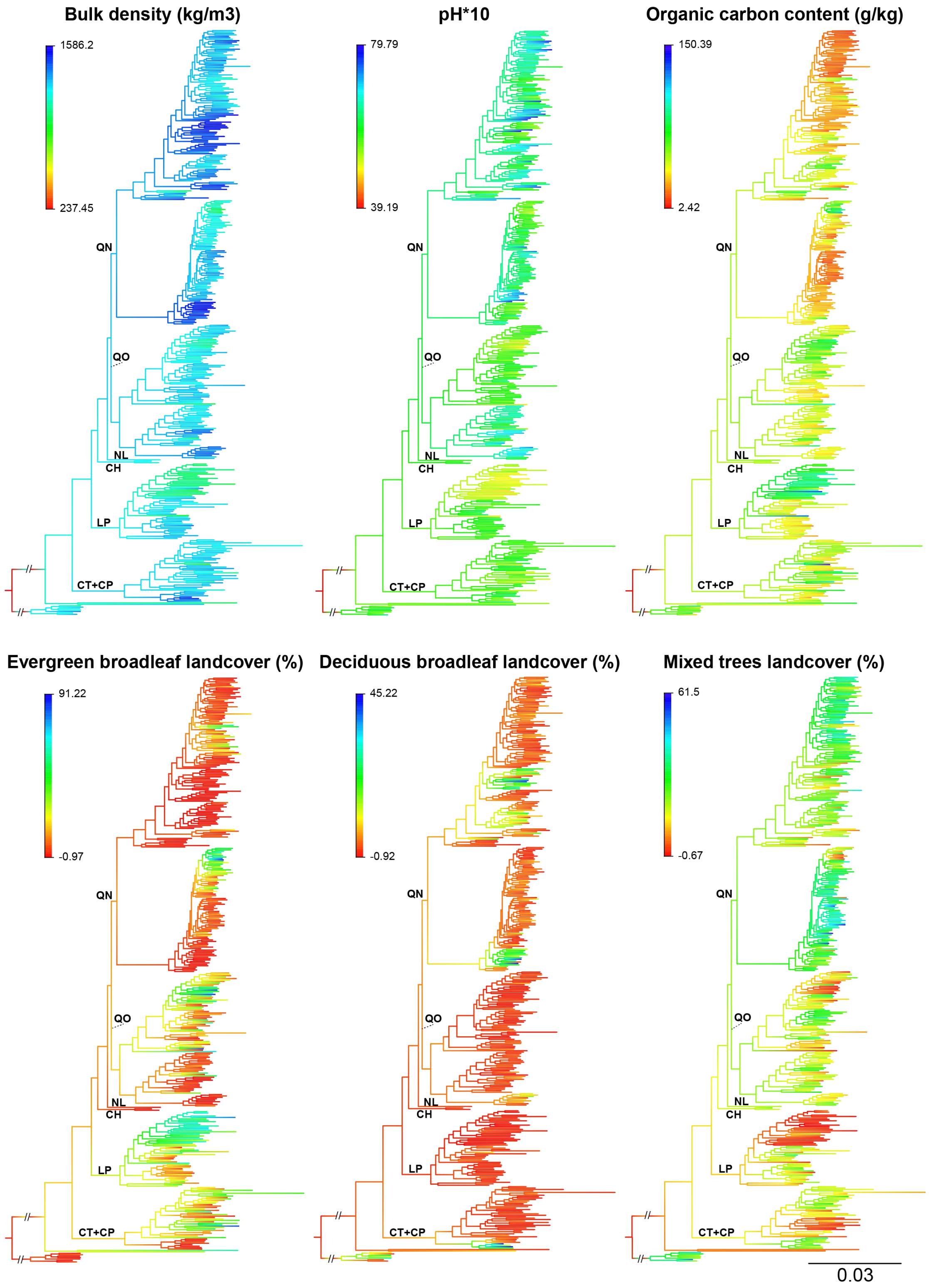


**FIGURE S32. Median values of ecological niche space across Fagaceae phylogeny, based on ancestral niche reconstructions on the tree produced from the Hyb-Seq dataset.** Tip labels (excluded here to save space) can be found in

Supplementary Figure S6. Abbreviations: QN, the New World oak clade; QO, the Old World oak clade; NL, *Notholithocarpus*; CH, *Chrysolepis*; LP, *Lithocarpus*; CT+CP, *Castanea* + *Castanopsis*.

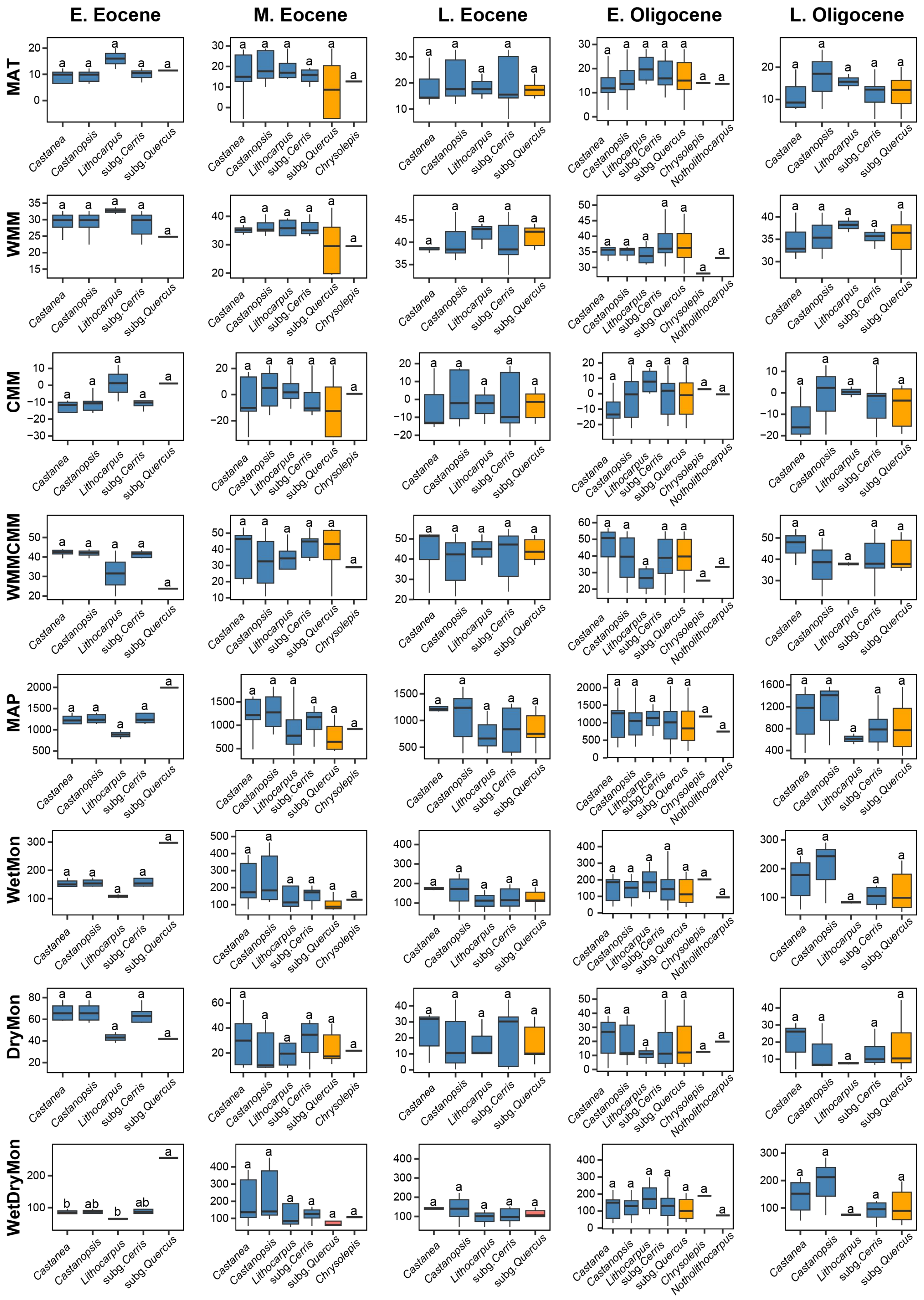


FIGURE S33. Boxplots showing climatic niche space from Early Eocene to Late Oligocene of fossil taxa of *Quercus* lineages and relatives. The same letters above boxes indicate no significantly climatic difference inferred by function “LSD.test()”

in the R package agricolae (de Mendiburu 2009). Blue boxes denote the Old World lineages, while orange boxes denote the New World lineages. Abbreviations: MAT, mean annual temperature (°C); WMM, warmest month mean surface air temperature (°C); CMM, coldest month mean surface air temperature (°C); WMMCMM, warmest month–coldest month temperature difference (°C); MAP, mean annual precipitation (mm); WetMon, wettest month precipitation (mm); DryMon, driest month precipitation (mm); WetDryMon, wettest month–driest month precipitation difference (mm).

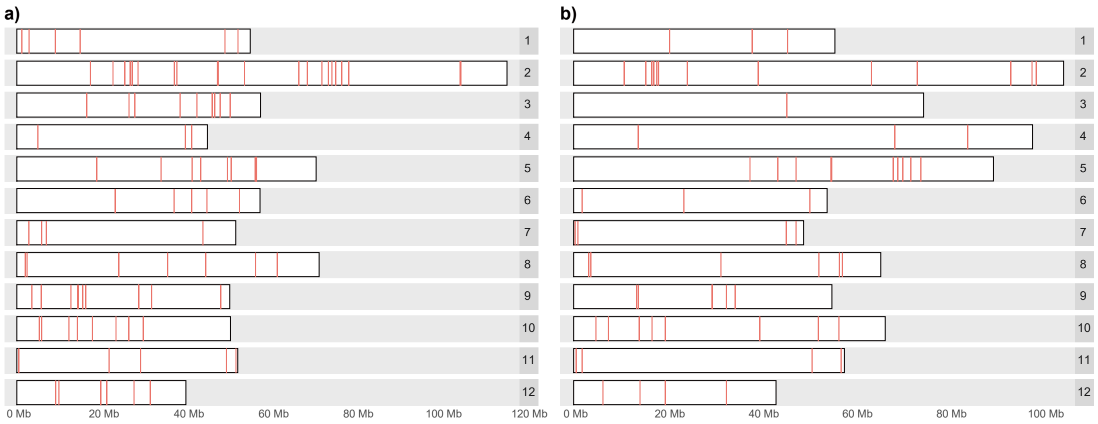


**FIGURE S34. Genomic locations of the targeted loci.** The red bands denote the genomic location of targeted genes. **a)** Distribution of targeted loci on the 12 chromosomes of the genome of *Quercus lobata*. **b)** Distribution of targeted loci on the 12 chromosomes of the genome of *Q. robur*.

### SUPPLEMENTARY TABLES

**TABLE S1.** Taxon sampling percentage for each genus of Fagaceae and each section of oaks in the Hyb-Seq, transcriptome, and plastome datasets.

**TABLE S2.** Information for taxon sampling and Hyb-Seq and transcriptome assembly.

**TABLE S3.** A summary of all datasets used in this study.

**TABLE S4.** Information for taxon sampling and plastome assembly.

**TABLE S5.** The five uncertain deep nodes and their alternative topologies as well as AU tests and polytomy tests.

**TABLE S6.** Prior settings and estimated ages of nine calibrated nodes in BEAST analysis.

**TABLE S7.** Number of positive tests from the $D_{\mathrm{FOIL}}$ tests categorized by genera in Quercoideae.

**TABLE S8.** Numbers of positive $D_{\mathrm{FOIL}}$ introgression tests per genus (outside *Quercus*) or *Quercus* subgenus based on the HYB-98RT dataset.

**TABLE S9.** The 95% confidence interval of 12 observed environmental variables values of *Notholithocarpus*.

**TABLE S10.** Number of positive $D_{\mathrm{FOIL}}$ introgression tests per genus (outside *Quercus*) or *Quercus* subgenus based on the RNA-2821RT dataset.

**TABLE S11.** Model selection among species networks recovered for oaks and their relatives based on the Hyb-Seq dataset.

**TABLE S12.** Model selection among species networks recovered for oaks and their relatives based on the transcriptome dataset.

**TABLE S13.** Comparison of three biogeographical models based on the Hyb-Seq dataset.

**TABLE S1. Taxon sampling percentage for each genus of Fagaceae and each section of oaks in the Hyb-Seq, transcriptome, and plastome datasets.** "a", The species numbers of *Quercus* and eight sections are from the main text in Denk et al. (2017), the numbers in parentheses are from the accepted number based on the "World Checklist" column in the species list of an appended XLSX file in Denk et al. (2017), and the numbers of the other genera of Fagaceae are from the website World Checklist of Selected Plant Families (https://wcsp.science.kew.org/qsearch.do; accessed 10 March 2020). "b", "Plastome" here means chloroplast dataset which includes chloroplast genomes consisting of assembled sequences of one inverted repeat region, small single-copy region, and large single-copy region. "c", This number has been widely used in recent oak studies (e.g., Deng et al. 2018; Hipp et al. 2018; Kremer and Hipp 2020). The numbers in parentheses of the third, fourth, and fifth columns denote the sampling percent of a particular group. HYB-DB, Hyb-Seq dataset; and RNA-DB, transcriptome dataset.

| Section | Species[a] | HYB-DB | RNA-DB | Plastome[b] |
|---|---|---|---|---|
| Genus *Quercus* | ~481 (524; 435[c]) | 314 (65/60/72%) | 55 (11/10/13%) | 175 (36/33/40%) |
| *Quercus* section | | | | |
| *Cerris* | ~13 (11) | 9 (69%) | 7 (54%) | 10 (77%) |
| *Ilex* | ~32 (39) | 30 (94%) | 2 (6%) | 18 (56%) |
| *Cyclobalanopsis* | ~101 (116) | 59 (58%) | 5 (5%) | 32 (32%) |
| *Lobatae* | ~162 (160) | 94 (58%) | 13 (8%) | 47 (29%) |
| *Protobalanus* | 5 (5) | 5 (100%) | 3 (60%) | 4 (80%) |
| *Ponticae* | 2 (2) | 2 (100%) | 0 (0%) | 2 (100%) |
| *Virentes* | 6 (6) | 6 (100%) | 1 (17%) | 3 (50%) |
| *Quercus* | ~160 (155) | 109 (68%) | 24 (15%) | 59 (37%) |
| Genus *Notholithocarpus* | 1 | 1 (100%) | 1 (100%) | 1 (100%) |
| Genus *Chrysolepis* | 2 | 2 (100%) | 1 (50%) | 2 (100%) |
| Genus *Lithocarpus* | 340 | 53 (16%) | 8 (2%) | 17 (5%) |
| Genus *Castanopsis* | 143 | 36 (25%) | 3 (2%) | 11 (8%) |
| Genus *Castanea* | 8 | 8 (100%) | 6(75%) | 7 (88%) |
| Genus *Trigonobalanus* | 3 | 2 (67%) | 1 (33%) | 1 (33%) |
| Genus *Fagus* | 11 | 7 (64%) | 6 (55%) | 8 (73%) |
| Outgroup | - | 8 | 8 | 1 |
| **Total** | **-** | **431** | **89** | **223** |

**TABLE S3. A summary of all datasets used in this study.** Ingroup here means Fagaceae species. The seventh, eighth, and ninth columns are the characteristics of the concatenated alignment of each dataset. "*" means that this dataset is also used in other analyses, as detailed in Supplementary Figures S1 and S2.

| **Dataset** | **Orthology inference** | **Locus length (bp)** | **φ test** | **Ingroups for each locus** | **Loci number** | **Alignment length (bp)** | **Informative sites (bp)** | **Gaps (%)** | **Analysis** |
|---|---|---|---|---|---|---|---|---|---|
| HYB-DB | | | | | | | | | |
| HYB-89MO | MO | ≥400 | $P>0.05$ | ≥106 | 89 | 96,122 | 59,886 | 21.78 | RAxML; ASTRAL-III |
| HYB-98RT | RT | ≥400 | $P>0.05$ | ≥106 | 98 | 105,564 | 64,772 | 25.86 | RAxML; ASTRAL-III; * |
| HYB-114RT | RT | ≥400 | $P>0.05$ | ≥50 | 114 | - | - | - | PhyloNet |
| RNA-DB | | | | | | | | | |
| RNA-977MO | MO | ≥1000 | $P>0.05$ | ≥41 | 977 | 2,229,405 | 904,917 | 52.03 | RAxML; ASTRAL-III |
| RNA-2821RT | RT | ≥1000 | $P>0.05$ | ≥41 | 2821 | 6,050,182 | 1,704,829 | 55.25 | RAxML; ASTRAL-III;* |
| RNA-4853RT | RT | ≥1000 | $P>0.05$ | ≥30 | 4853 | - | - | - | PhyloNet |
| Plastome | - | - | - | - | - | 135,243 | 116,579 | 28.71 | RAxML |

**TABLE S5. The five uncertain deep nodes and their alternative topologies as well as AU tests and polytomy tests.** Dashed branches denote other clades (which are not shown in the 3(4)-taxon tree) located at this branch. “#” means subg. *Quercus* with sect. *Lobatae* excluded. “a” means that values in parentheses are the *P* value from the RNA-2821RT dataset and otherwise from the HYB-98RT dataset, and the bold numbers denote *P* < 0.05.

| No. | Uncertain Nodes | T1 | T2 | T3 | Polytomy test[a] | AU test[a] |
|---|---|---|---|---|---|---|
| A | Position of *Castanea* + *Castanopsis* | 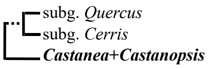 | 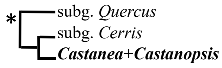 | | **0** (**0**) | T1/**T2** (T1/**T2**) |
| B | Position of *Lithocarpus* | 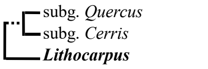 | 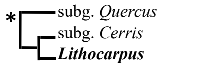 | | **0** (**0**) | T1/**T2** (T1/**T2**) |
| C | Position of *Chrysolepis* | 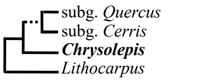 | 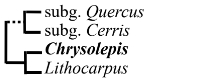 | 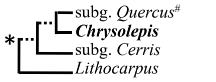 | 0.174 (**0.004**) | T1/T2/**T3** (T1/**T2**/**T3**) |
| D | Position of *Notholithocarpus* | 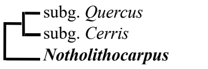 | 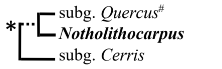 | | **0.004** (**0**) | T1/**T2** (T1/**T2**) |
| E | Position of *Quercus* sect. *Cyclobalanopsis* | 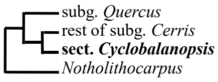 | 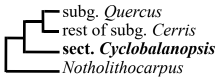 | 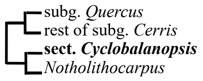 | 0.913 (**0**) | **T1**/T2/T3 (T1/**T2**/**T3**) |

**TABLE S6. Prior settings and estimated ages of nine calibrated nodes in BEAST analysis.** All nodes are calibrated in the Hyb-Seq dataset while only No. 1–5 and 8 are calibrated in the transcriptome dataset for incomplete taxon sampling within sections of *Quercus*. CG, crown group; SG, stem group. U, uniform distribution; L, lognormal distribution. "#" denotes uniform distribution with two fossil ages as the maximum and minimum age and lognormal distribution with fossil age as the median age and a standard deviation of 0.5 (the number between parentheses is the 95% highest posterior density [HPD] interval). "*" denotes the mean and 95% HPD interval of node age extracted from maximum clade credibility trees based on the Hyb-Seq dataset (left of symbol "/") and the transcriptome dataset (right of symbol "/"). Abbreviations in column Refs: a, Takahashi et al. 2008; b, Grímsson et al. 2016; c, Manchester and Dillhoff 2004; d, Wilf et al. 2019; e, Hofmann 2010; f, Manchester 1994; g, Chen et al. 2021; h, Su et al. 2019; i, Pavlyutkin et al. 2014; and j, McIver and Basinger 1999. All units of age are Ma.

| No. | Nodes | Fossils | Type | Fossil Age | Priors[#] | Estimated Age[*] | Refs |
|---|---|---|---|---|---|---|---|
| 1 | CG Fagaceae | *Archaefagacea futabensis* Takahashi, Friis, Herendeen et Crane gen. et sp. nov.; Fagoideae PT 1 | flowers and fruits; pollen | early Coniacian; 82–81 | U: 81.0-89.8 | 89.37 (88.52-89.81) / 88.29 (85.36-89.81) | a; b |
| 2 | CG genus *Fagus* | *Fagus langevinii* Manchester & Dilhoff | foliage, pollen, cupule/nut | 47.0–53.0 | L: 47.0 (45.3-51.5) | 45.95 (44.83-47.25) / 46.44 (44.91-48.32) | c |
| 3 | SG genus *Castanopsis* | *Castanopsis rothwellii* Wilf, Nixon, Gandolfo et Cúneo | two infructescences | 52.22± 0.22 | L: 52.2 (50.5-56.7) | 52.05 (50.22-54.33) / 52.05 (49.98-54.7) | d |
| 4 | SG genus *Quercus* | *Quercus* type 1A | pollen | Late Thanetian to Early Ypresian age | L: 56.0 (54.3-60.5) | 55.34 (54.00-56.87) / 55.29 (53.93-57.00) | e |
| 5 | SG sect. *Cyclobalanopsis* | *Quercus paleocarpa* Manchester | fruit | 48.32±0.11 | L: 48.3 (46.6-52.8) | 49.61 (47.24-51.96) / 50.34 (47.23-53.42) | f |
| 6 | SG the Himalayan alpine clade in sect. *Ilex* | *Quercus* cf. *presenescens* Z. K. Zhou | foliage | 34.6±0.8 – 35.5±0.3 | L: 34.6 (32.9-39.1) | 33.30 (32.42-34.35) / - | g; h |
| 7 | SG the East Asian clade in sect. *Cerris* | *Quercus gracilis* (Pavlyutkin) Pavlyutkin | foliage | 34-30 | L: 30 (28.3-34.5) | 29.43 (27.95-31.24) / - | i |
| 8 | SG sect. *Lobatae* | *Quercus* PT 1 aff. Group *Lobatae* | pollen | 47.8 | L: 48.7 (47-53.2) | 50.61 (48.06-53.05) / 48.47 (46.78-50.57) | b |
| 9 | CG sect. *Quercus* | *Quercus* L. | foliage | middle to late Eocene | L: 39.4 (37.7-43.9) | 38.17 (38.73-42.69) / - | j |

**TABLE S7. Number of positive tests from the $D_{\mathrm{FOIL}}$ tests categorized by genera in Quercoideae.** The directionality of introgression signature is indicated by the arrow.

| No. | Signatures | Number of positive tests | |
|---|---|---|---|
| | | HYB-98RT | RNA-2821RT |
| 1 | *Castanea* –> Chrysolepis | - | 1 |
| 2 | *Castanea* –> *Lithocarpus* | 1 | 1 |
| 3 | *Castanea* –> *Quercus* | 98 | 24 |
| 4 | *Castanea* / *Castanea* <–> *Chrysolepis* | - | 1 |
| 5 | *Castanea* / *Castanea* <–> *Castanopsis* | 10 | - |
| 6 | *Castanea* / *Castanea* <–> *Lithocarpus* | 1,918 | 26 |
| 7 | *Castanea* / *Castanea* <–> *Notholithocarpus* | - | 1 |
| 8 | *Castanea* / *Castanea* <–> *Quercus* | 3,995 | 657 |
| 9 | *Castanopsis* –> *Lithocarpus* | 234 | 1 |
| 10 | *Castanopsis* –> *Quercus* | 145 | 3 |
| 11 | *Castanopsis* / *Castanea* <–> *Chrysolepis* | - | 9 |
| 12 | *Castanopsis* / *Castanea* <–> *Lithocarpus* | 9,666 | 39 |
| 13 | *Castanopsis* / *Castanea* <–> *Notholithocarpus* | - | 8 |
| 14 | *Castanopsis* / *Castanea* <–> *Quercus* | 8,253 | 41 |
| 15 | *Castanopsis* / *Castanopsis* <–> *Castanea* | 1 | - |
| 16 | *Castanopsis* / *Castanopsis* <–> *Lithocarpus* | 37,738 | 8 |
| 17 | *Castanopsis* / *Castanopsis* <–> *Notholithocarpus* | 18 | 1 |
| 18 | *Castanopsis* / *Castanopsis* <–> *Quercus* | 19,889 | 45 |
| 19 | *Lithocarpus* –> *Castanea* | 14 | 1 |
| 20 | *Lithocarpus* –> *Castanopsis* | 45 | 3 |
| 21 | *Lithocarpus* –> *Chrysolepis* | - | 8 |
| 22 | *Lithocarpus* –> *Notholithocarpus* | 1 | 1 |
| 23 | *Lithocarpus* –> *Quercus* | 169 | 39 |
| 24 | *Lithocarpus* / *Lithocarpus* <–> *Castanea* | 26 | 12 |
| 25 | *Lithocarpus* / *Lithocarpus* <–> *Castanopsis* | 16 | 7 |
| 26 | *Lithocarpus* / *Lithocarpus* <–> *Chrysolepis* | 63 | 140 |
| 27 | *Lithocarpus* / *Lithocarpus* <–> *Notholithocarpus* | 306 | 62 |
| 28 | *Lithocarpus* / *Lithocarpus* <–> *Quercus* | 10,998 | 1163 |
| 29 | *Chrysolepis* / *Chrysolepis* <–> *Castanopsis* | 2 | - |
| 30 | *Chrysolepis* / *Chrysolepis* <–> *Lithocarpus* | 2 | - |
| 31 | *Chrysolepis* / *Chrysolepis* <–> *Notholithocarpus* | 58 | - |
| 32 | *Chrysolepis* / *Chrysolepis* <–> *Quercus* | 931 | - |
| 33 | *Notholithocarpus* –> *Castanea* | - | 1 |
| 34 | *Notholithocarpus* –> *Lithocarpus* | - | 4 |
| 35 | *Quercus* –> *Castanea* | 66 | 13 |

| | | | |
|---|---|---|---|
| 36 | *Quercus* –> *Castanopsis* | 95 | 15 |
| 37 | *Quercus* –> *Lithocarpus* | 577 | 56 |
| 38 | *Quercus* –> *Quercus* | 1,220 | 451 |
| 39 | *Quercus* / *Quercus* <–> *Castanea* | 892 | 393 |
| 40 | *Quercus* / *Quercus* <–> *Castanopsis* | 790 | 130 |
| 41 | *Quercus* / *Quercus* <–> *Lithocarpus* | 2,370 | 353 |
| 42 | *Quercus* / *Quercus* <–> *Quercus* | 31,828 | 4,340 |
| 43 | no introgression | 5,184,264 | 216,940 |

**TABLE S8. Numbers of positive $D_{\mathrm{FOIL}}$ introgression tests per genus (outside *Quercus*) or *Quercus* subgenus based on the HYB-98RT dataset.** Results here are from a reduced 150-taxon set of the HYB-98RT dataset including 49 species of *Quercus* subg. *Cerris*, 51 species of *Quercus* subg. *Quercus*, and 50 species of closely related genera. "X" means *Quercus* subg. *Cerris* or *Quercus* subg. *Quercus* (i.e., the second or third column). The directionality of the introgression signature is indicated by the arrow. Grey row indicates signature of ancestral introgression, while white row indicates signature of intergroup introgression. Group with "#" only (or nearly) distributed in the New World, while other groups only (or nearly) distributed in the Old World.

| Introgression Signatures | *Quercus* subg. *Cerris* | *Quercus* subg. *Quercus*# | Total |
|---|---|---|---|
| *Castanea* –> X | 92 | 6 | 98 |
| *Castanea* / *Castanea* <–> X | 2,931 | 1,064 | 3,995 |
| *Castanopsis* –> X | 145 | 0 | 145 |
| *Castanopsis* / *Castanea* <–> X | 7,253 | 1,000 | 8,253 |
| *Castanopsis* / *Castanopsis* <–> X | 19,355 | 534 | 19,889 |
| *Lithocarpus* –> X | 154 | 15 | 169 |
| *Lithocarpus* / *Lithocarpus* <–> X | 7,147 | 3,851 | 10,998 |
| *Chrysolepis*# / *Chrysolepis*# <–> X | 201 | 730 | 931 |
| X –> *Castanea* | 65 | 1 | 66 |
| X –> *Castanopsis* | 95 | 0 | 95 |
| X –> *Lithocarpus* | 476 | 101 | 577 |
| X –> *Quercus* subg. *Cerris* | 39 | 175 | 214 |
| X –> *Quercus* subg. *Quercus*# | 994 | 12 | 1,006 |
| X/X <–> *Castanea* | 483 | 409 | 892 |
| X/X <–> *Castanopsis* | 700 | 90 | 790 |
| X/X <–> *Lithocarpus* | 1,258 | 1,112 | 2,370 |
| X/X <–> *Quercus* subg. *Cerris* | 1,677 | 16,905 | 18,582 |
| X/X <–> *Quercus* subg. *Quercus*# | 12,421 | 825 | 13,246 |

**TABLE S9. The 95% confidence interval of 12 observed environmental variables values of *Notholithocarpus*.**

| **Environmental variables** | **2.5% quantiles** | **97.5% quantiles** |
|---|---|---|
| Mean annual temperature (°C) | 6.77 | 14.62 |
| Temperature annual range (°C) | 15.78 | 33.04 |
| Mean annual precipitation (mm) | 549.80 | 2152.00 |
| Precipitation seasonality (mm) | 67.77 | 95.32 |
| Elevation (m) | 59.60 | 1642.40 |
| Slope (°) | 1 | 15 |
| Bulk density (kg/m$^3$) | 956.20 | 1480.20 |
| pH*10 | 52 | 65 |
| Organic carbon content (g/kg) | 29 | 97 |
| Evergreen broadleaf landcover (%) | 0 | 0 |
| Deciduous broadleaf landcover (%) | 0 | 24 |
| Mixed trees landcover (%) | 0.00 | 54.15 |

**TABLE S10. Number of positive $D_{\mathrm{FOIL}}$ introgression tests per genus (outside Quercus) or Quercus subgenus based on the RNA-2821RT dataset.** Results here are from a reduced 74-taxon set of the RNA-2821RT dataset including 14 species of *Quercus* subg. *Cerris*, 41 species of *Quercus* subg. *Quercus*, and 19 species of closely related genera. "X" means *Quercus* subg. *Cerris* or *Quercus* subg. *Quercus* (i.e., the second or third column). The directionality of the introgression signature is indicated by the arrow. Grey row indicates signature of ancestral introgression, while white row indicates signature of intergroup introgression. Group with "#" only (or nearly) distributed in the New World, while other groups only (or nearly) distributed in the Old World.

| **Introgression Signatures** | ***Quercus* subg. *Cerris*** | ***Quercus* subg. *Quercus*[#]** | **Total** |
|---|---|---|---|
| *Castanea* –> X | 5 | 19 | 24 |
| *Castanea* / *Castanea* <–> X | 170 | 487 | 657 |
| *Castanopsis* –> X | 3 | 0 | 3 |
| *Castanopsis* / *Castanea* <–> X | 5 | 36 | 41 |
| *Castanopsis* / *Castanopsis* <–> X | 12 | 33 | 45 |
| *Lithocarpus* –> X | 22 | 17 | 39 |
| *Lithocarpus* / *Lithocarpus* <–> X | 649 | 514 | 1163 |
| X –> *Castanea* | 1 | 12 | 13 |
| X –> *Castanopsis* | 2 | 13 | 15 |
| X –> *Lithocarpus* | 15 | 41 | 56 |
| X –> *Quercus* subg. *Cerris* | 4 | 125 | 129 |
| X –> *Quercus* subg. *Quercus*[#] | 89 | 233 | 322 |
| X/X <–> *Castanea* | 15 | 330 | 345 |
| X/X <–> *Castanopsis* | 13 | 107 | 120 |
| X/X <–> *Lithocarpus* | 14 | 339 | 353 |
| X/X <–> *Quercus* subg. *Cerris* | 18 | 1440 | 1458 |
| X/X <–> *Quercus* subg. *Quercus*[#] | 1117 | 1765 | 2882 |
| *Quercus* subg. *Cerris* / *Quercus* subg. *Quercus*[#] <–> *Castanea* | 48 | - | 48 |
| *Quercus* subg. *Cerris* / *Quercus* subg. *Quercus*[#] <–> *Castanopsis* | 10 | - | 10 |

**TABLE S11. Model selection among species networks recovered for oaks and their relatives based on the Hyb-Seq dataset.** The best model with the lowest information criterion under each taxon set is highlighted in bold. Network with "-" denotes that we did not obtain results of the PhyloNet analysis under the specified setting even after running for over two months.

| Taxon set | Topology | Maximum number of reticulations added | Number of detected reticulations | Ln(L) | Parameters | Number of loci | AICc | ΔAICc | BIC | ΔBIC |
|---|---|---|---|---|---|---|---|---|---|---|
| **planA** | ASTRAL | NA | NA | -1226.093 | 30 | 100 | 2539.142 | 342.437 | 2590.341 | 342.135 |
| | Network 1 | 1 | 1 | -1104.665 | 32 | 100 | 2304.853 | 108.148 | 2356.696 | 108.490 |
| | Network 2 | 2 | 1 | -1109.009 | 32 | 100 | 2313.540 | 116.834 | 2365.383 | 117.177 |
| | Network 3 | 3 | 2 | -1103.301 | 34 | 100 | 2311.218 | 114.513 | 2363.179 | 114.973 |
| | **Network 4** | **4** | **3** | **-1041.210** | **36** | **100** | **2196.706** | **0** | **2248.206** | **0** |
| | Network 5 | 5 | 3 | -1102.638 | 36 | 100 | 2319.562 | 122.857 | 2371.063 | 122.857 |
| | Network 6 | 6 | 4 | -1101.783 | 38 | 100 | 2328.156 | 131.450 | 2378.562 | 130.356 |
| **planB** | ASTRAL | NA | NA | -2148.805 | 30 | 98 | 4385.371 | 509.732 | 4435.159 | 510.133 |
| | Network 1 | 1 | 1 | -1945.627 | 32 | 98 | 3987.747 | 112.108 | 4037.973 | 112.948 |
| | Network 2 | 2 | 2 | -1910.570 | 34 | 98 | 3926.918 | 51.279 | 3977.029 | 52.004 |
| | Network 3 | 3 | 3 | -1933.154 | 36 | 98 | 3981.980 | 106.341 | 4031.366 | 106.341 |
| | Network 4 | 4 | 3 | -1931.121 | 36 | 98 | 3977.913 | 102.274 | 4027.300 | 102.274 |
| | Network 5 | 5 | 5 | -1929.031 | 40 | 98 | 3995.606 | 119.967 | 4041.461 | 116.435 |
| | **Network 6** | **6** | **3** | **-1879.983** | **36** | **98** | **3875.639** | **0** | **3925.026** | **0** |
| **planC** | ASTRAL | NA | NA | -1910.046 | 32 | 97 | 3917.092 | 201.476 | 3966.483 | 205.461 |
| | Network 1 | 1 | 1 | -1822.477 | 34 | 97 | 3751.341 | 35.725 | 3800.494 | 39.472 |
| | Network 2 | 2 | 2 | -1799.608 | 36 | 97 | 3715.616 | 0 | 3763.905 | 2.884 |
| | Network 3 | 3 | 3 | -1804.395 | 38 | 97 | 3735.894 | 20.278 | 3782.629 | 21.608 |
| | **Network 4** | **4** | **4** | **-1789.017** | **40** | **97** | **3716.605** | **0.989** | **3761.022** | **0** |
| | Network 5 | 5 | - | - | - | - | - | - | - | - |
| | Network 6 | 6 | 4 | -1796.176 | 40 | 97 | 3730.923 | 15.308 | 3775.340 | 14.319 |

**TABLE S12. Model selection among species networks recovered for oaks and their relatives based on the transcriptome dataset.** The best model with the lowest information criterion under each taxon set is highlighted in bold. Network with "-" denotes that we did not obtain results of the PhyloNet analysis under the specified setting even after running for over two months.

| Taxon set | Topology | Maximum number of reticulations added | Number of detected reticulations | Ln(L) | Parameters | Number of loci | AICc | ΔAICc | BIC | ΔBIC |
|---|---|---|---|---|---|---|---|---|---|---|
| **planA** | ASTRAL | NA | NA | -22753.573 | 26 | 4466 | 45559.462 | 2833.302 | 45725.656 | 2795.037 |
| | Network 1 | 1 | 0 | -21911.068 | 26 | 4466 | 43874.451 | 1148.292 | 44040.645 | 1110.026 |
| | Network 2 | 2 | 1 | -21670.087 | 28 | 4466 | 43396.539 | 670.379 | 43575.492 | 644.873 |
| | Network 3 | 3 | 1 | -21898.780 | 28 | 4466 | 43853.926 | 1127.767 | 44032.879 | 1102.260 |
| | Network 4 | 4 | 1 | -21870.472 | 28 | 4466 | 43797.310 | 1071.150 | 43976.263 | 1045.643 |
| | **Network 5** | **5** | **3** | **-21330.842** | **32** | **4466** | **42726.160** | **0** | **42930.619** | **0** |
| | Network 6 | 6 | 2 | -21567.527 | 30 | 4466 | 43195.474 | 43195.474 | 43387.182 | 43387.182 |
| **planB** | ASTRAL | NA | NA | -38788.631 | 28 | 4602 | 77633.617 | 4926.615 | 77813.421 | 4862.567 |
| | Network 1 | 1 | 1 | -36804.499 | 30 | 4602 | 73669.405 | 962.402 | 73862.025 | 911.171 |
| | Network 2 | 2 | 2 | -36592.473 | 32 | 4602 | 73249.407 | 542.405 | 73454.841 | 503.986 |
| | Network 3 | 3 | 2 | -36830.427 | 32 | 4602 | 73725.317 | 1018.314 | 73930.751 | 979.896 |
| | Network 4 | 4 | - | - | - | - | - | - | - | - |
| | **Network 5** | **5** | **5** | **-36315.177** | **38** | **4602** | **72707.003** | **0** | **72950.854** | **0** |
| | Network 6 | 6 | 3 | -36844.847 | 34 | 4602 | 73758.214 | 73758.214 | 73976.458 | 73976.458 |
| **planC** | ASTRAL | NA | NA | -37843.083 | 26 | 4669 | 75738.469 | 2313.989 | 75905.833 | 2288.293 |
| | Network 1 | 1 | 1 | -36858.988 | 28 | 4669 | 73774.327 | 349.847 | 73954.540 | 337.001 |
| | Network 2 | 2 | 2 | -36939.579 | 30 | 4669 | 73939.559 | 515.079 | 74132.619 | 515.079 |
| | Network 3 | 3 | 1 | -36856.995 | 28 | 4669 | 73770.339 | 345.860 | 73950.553 | 333.013 |
| | **Network 4** | **4** | **2** | **-36682.039** | **30** | **4669** | **73424.480** | **0** | **73617.540** | **0** |
| | Network 5 | 5 | 3 | -36787.502 | 32 | 4669 | 73639.459 | 214.979 | 73845.362 | 227.822 |
| | Network 6 | 6 | 2 | -36873.895 | 30 | 4669 | 73808.190 | 383.711 | 74001.250 | 383.711 |

**TABLE S13. Comparison of three biogeographical models based on the Hyb-Seq dataset.** The best model with the lowest information criterion is highlighted in bold.

| Model | LnL | Parameter | d | e | AIC | AIC_weight |
|---|---|---|---|---|---|---|
| **DEC** | **-594.6** | **2** | **0.0035** | **1.00E-12** | **1193** | **1.00** |
| DIVALIKE | -620.4 | 2 | 0.0042 | 1.00E-12 | 1245 | 5.11E-12 |
| BAYAREALIKE | -747.4 | 2 | 0.01 | 0.01 | 1499 | 3.57E-67 |
