## Supplementary Methods and Results for "Phylogenomic Analyses Reveal Widespread Gene Flow During the Early Radiation of Oaks and Relatives (Fagaceae: Quercoideae)"

*Taxon Sampling and Hybrid Enrichment Sequencing*

Our study includes 472 samples of Fagaceae (391 samples with Hyb-Seq data and 81 samples with transcriptomes; Supplementary Table S2) representing 314 species of *Quercus* and 109 species of its close relatives in Fagaceae (including 1 species of *Notholithocarpus*, 2 species of *Chrysolepis*, 53 species of *Lithocarpus*, 36 species of *Castanopsis*, 8 species of *Castanea*, 2 species of *Trigonobalanus*, and 7 species of *Fagus*). This sampling scheme was informed by the latest infrageneric classification of *Quercus* (Denk et al. 2017) and The Plant List (v.1.1; www.theplantlist.org). For outgroups beyond Fagaceae, we chose 15 species (including 7 species with Hyb-Seq data and 8 species with transcriptomes; Supplementary Table S2) representing the major lineages of Fagales: 4 species of Betulaceae, 1 species of Casuarinaceae, 4 species of Juglandaceae, 4 species of Myricaceae, 1 species of Nothofagaceae, and 1 species of Ticodendraceae. All Hyb-Seq data were newly generated for this study, while all transcriptomes were obtained from Yang et al. (2021) and GenBank (accessed 1 May 2021). Our samples for Hyb-Seq were collected from field and the following 10 herbaria: Botanical Research Institute of Texas; California Academy of Sciences; Field Museum of Natural History; Harvard University; Kunming Institute of Botany, Chinese Academy of Sciences; Missouri Botanical Garden; The Ohio State University; Smithsonian Institution; New York Botanical Garden; and University of Texas at Austin.

Genomic DNAs were extracted from herbarium or silica-dried materials using a modified CTAB protocol to maximize the yield of total DNA (Doyle and Doyle 1987). We used a set of exonic baits for 100 housekeeping genes designed for the nitrogen-fixing clade (which includes Fagaceae). These loci are evenly scattered across the 12 pseudo-chromosomes of *Quercus lobata* (Sork et al. 2016) and *Q. robur* (Schmid-Siegert et al. 2017) (Supplementary Fig. S34). Isolated DNAs were submitted to Rapid Genomics (Gainesville, FL, USA) for quantification, library preparation, hybrid enrichment with the NitFix loci, and multiplex sequencing on the Illumina HiSeq platform with 150-bp, paired-end reads. The sampling-to-sequencing workflow we followed, briefly described above, is described in detail by Folk et al. (2021).

*Read Processing, Assembly, and Orthology Inference*

*Hyb-Seq dataset*—Raw reads of Hyb-Seq data were cleaned using Trimmomatic v.0.36 (Bolger et al. 2014) to remove adapters and low-quality bases. The cleaned paired reads were assembled using HybPiper (Johnson et al. 2016). We also used HybPiper to assemble the 100 targeted genes from the transcriptome samples (representing 32 species of Fagaceae and 1 species of Betulaceae) in order to include these in the Hyb-Seq dataset (see below for further details on transcriptome samples). Prior to assembly, we developed a custom DNA reference including 100 DNA sequences based on the best BLAST hit from three published genomes—*Q. lobata* (Sork et al. 2016), *Q. robur* (Schmid-Siegert et al. 2017), and *Q. suber* (Ramos et al. 2018)—against the bait sequences. Using HybPiper, reads were first binned for each targeted gene using BWA v.0.7.12 (Li 2013), then assembled separately using SPAdes v.3.12.0 (Bankevich et al. 2012), and finally aligned to reference DNA sequences to obtain targeted regions using Exonerate (Slater and Birney 2005).

We detected and retrieved gene assemblies representing possible paralogs covering more than 85% of the reference sequence length using the HybPiper scripts "paralog_investigator.py" and "paralog_retriever.py". To infer orthologs, we generally followed the pipelines of Yang and Smith (2014) and Morales-Briones et al. (2021). Each gene cluster was first aligned using MAFFT v.7.305 (Katoh and Standley 2013), aligned columns with > 90% missing data were trimmed using Phyx (Brown et al. 2017), and then initial homolog trees were inferred using RAxML v.8.2.11 (Stamatakis 2014) under the GTR-CAT model with 100 fast bootstrap replicates. Spurious long tips were removed using the Python script "trim_tips.py" (https://bitbucket.org/dfmoralesb/target_enrichment_orthology/); monophyletic and paraphyletic tips belonging to the same species were collapsed (i.e., only the tip with the most unambiguous characters in the trimmed alignment was retained) using the Python script "mask_tips_by_taxonID_transcripts.py". We repeated the process of homolog tree inference and tip trimming three times to generate the final homologs. Orthology inference was first carried using the "monophyletic outgroup" (MO) approach from Yang and Smith (2014), which filters homolog trees with outgroups that are monophyletic and single-copy; for each homolog tree, the subtree (with both outgroups and ingroups) meeting a threshold of included species (in this case, ≥ 50 species) and having the most taxa among all candidate subtrees is retained. Compared with the MO approach, the "rooted ingroup" (RT) approach (also from Yang and Smith 2014) often yields more orthologs. We also applied the RT approach to orthology inference, which does not require the outgroups to be monophyletic and non-duplicated, but prunes the outgroups, leaving subtrees. Using the RT method, we also applied a threshold such that only subtrees with ≥ 50 ingroup taxa were retained. For downstream analyses, we added outgroups of the MO orthologs back to the RT orthologs from the same homolog.

The inferred orthologs were re-aligned, and columns with > 90% missing data were removed. To explore the presence of recombination in the dataset (given that most phylogenetic and species tree methods assume no recombination), we examined each ortholog using the permutation-based φ test implemented in PhiPack (Bruen et al. 2006). The orthologs that showed significant recombination ($P \leq 0.05$), were less than 400 bp in length, and had overall low-quality assemblies were excluded to avoid the inclusion of erroneous signal. Low-quality assemblies here refer to abundant abnormal regions in the alignments, when examined through follow-up visual inspections in Geneious v.8.0.2 (Kearse et al. 2012). The final Hyb-Seq dataset included 89 MO orthologs (HYB-89MO) and 114 RT orthologs (HYB-114RT). From these datasets, we generated one additional dataset that excluded loci with fewer than 106 taxa (~ 25%): 98 RT orthologs (HYB-98RT) (while all loci in HYB-89MO included over 106 taxa).

*Transcriptome dataset*—We applied the pipeline of Yang and Smith (2014) (https://bitbucket.org/yanglab/phylogenomic_dataset_construction/) to all 89 transcriptomes for read processing, assembly, transcript processing, and homology and orthology inference. Raw reads were first corrected for random sequencing error using Rcorrector (Song and Florea 2015), trimmed for adapters and low-quality bases using Trimmomatic, and filtered for organelle reads using Bowtie v.2.4.2 (Langmead and Salzberg 2012). *De novo* assembly was conducted using Trinity v.2.10.0 (Grabherr et al. 2011). We accessed the assembly quality using Transrate v.1.0.3 (Smith-Unna et al. 2016) and then removed the low-quality and chimeric transcripts. Cleaned transcripts were clustered as putative genes using Corset v.1.07 (Davidson and Oshlack 2014), and the longest transcript within a cluster was extracted as the representative transcript of each putative gene. All representative transcripts were translated using TransDecoder v.5.3.0 (Haas et al. 2013), and the redundancy of coding sequences (CDS) was reduced using CD-HIT v.4.7 (Fu et al. 2012).

To obtain the initial homologs, we performed an all-by-all BLASTN search on CDS, filtered the raw BLAST results by a hit fraction of 0.4, and then generated the putative gene clusters using MCL v.14-137 (van Dongen 2000). Each resulting cluster was aligned using MAFFT, and columns with > 70% missing data were removed using Phyx. The process of homolog tree inference and tip trimming was the same as that for the Hyb-Seq dataset, except that we also cut long internal branches possibly representing deep paralogs, with subclades including ≥ 30 tips retained. The steps from homolog tree inferences to deep-paralog cutting were repeated three times to obtain the final homologs. We used the MO and RT approaches for orthology inference, keeping the orthologs with ≥ 30 taxa. The resulting orthologs were re-aligned individually, and columns with > 70% missing data were removed. Cleaned alignments with significant recombination ($P \leq 0.05$) and fewer than 1000 bp were excluded. The final transcriptome datasets include 2150 MO orthologs (RNA-2150MO) and 4853 RT orthologs (RNA-4853RT); two additional datasets were generated from these after excluding loci with fewer than 41 taxa (~ 50%): 977 MO orthologs (RNA-977MO) and 2821 RT orthologs (RNA-2821RT).

*Plastome Assembly*—We newly assembled plastomes from 391 Hyb-Seq samples and 60 transcriptomes (Supplementary Table S4) with a modified bash script (https://github.com/ryanafolk/Assembly-tools/). Specifically, we extracted the large single-copy region (LSC), small single-copy region (SSC), and inverted repeat (IR) of 11 annotated plastomes of Fagaceae species from GenBank (Supplementary Table S4) for use as references. Cleaned reads of each sample were mapped to the closest reference using BWA, and the LSC, SSC, and IR consensus sequences were then called using Samtools v.1.9 (Li et al. 2009) and Bcftools v.1.9 (Li 2011). In addition, 47 complete chloroplast genomes of Fagaceae and one of Betulaceae (used as an outgroup) were downloaded from GenBank (Supplementary Table S4; accessed 5 April 2021), and the LSC, SSC, and IR sequences of these plastomes were extracted in Geneious. LSC, SSC, and IR sequences were aligned with MAFFT. The columns of alignments with > 70% missing data were stripped in Geneious. Samples with > 80% missing data in the plastome assemblies (here LSC + SSC + one IR) and showing outlier long branches (based on visualization of a preliminary chloroplast phylogeny) were removed. We then removed extra accessions such that the final dataset only included one representative for each species. The final dataset included 222 ingroups (representing 175 species of *Quercus*, 1 species of *Notholithocarpus*, 2 species of *Chrysolepis*, 17 species of *Lithocarpus*, 11 species of *Castanopsis*, 7 species of *Castanea*, 1 species of *Trigonobalanus*, and 8 species of *Fagus*) and one outgroup from outside Fagaceae (*Carpinus monbeigiana*, Betulaceae, used to root the chloroplast tree). Additionally, 64 additional samples with long branches were removed before subsequent dating analyses (158 Fagaceae species were retained). Detailed sampling and processing information of chloroplast assemblies is given in Supplementary Table S4.

*Concordance Analyses*

We performed a phyparts analysis (Smith et al. 2015) using gene trees with low-supported branches (i.e., < 70% bootstrap support) collapsed; these gene trees were mapped against the ASTRAL species tree. For each bipartition in the species tree, phyparts quantifies the number of genes supporting that bipartition, the number supporting a main alternative, the number supporting other alternatives, and the number of uninformative genes (due to low bootstrap support or inadequate taxon sampling). Visualization of results was carried out using the script “phypartspiecharts.py” (https://github.com/mossmatters/phyloscripts/). Moreover, to distinguish weakly supported branches from those with strong conflict, we conducted

Quartet Sampling analysis (QS) (Pease et al. 2018) with 1000 replicates and using the ASTRAL species tree and concatenated alignment as the inputs. The QS method provides scores for three different metrics—quartet concordance, quartet differential, and quartet informativeness—to estimate the consistency with the given branch, the presence of gene flow, and the amount of information, using the sampled frequency of three possible quartet topologies at every given branch.

*Tree Topology Tests*

We identified five uncertain and conflicting deep nodes involving nuclear-nuclear and/or cytonuclear discordance, with some of these also evident in previous studies (e.g., Manos et al. 1999, 2008; Hubert et al. 2014; Simeone et al. 2016; Yang et al. 2021). These nodes concern the phylogenetic positions of (1) *Castanea* + *Castanopsis*, (2) *Lithocarpus*, (3) *Chrysolepis*, (4) *Notholithocarpus*, and (5) *Quercus* sect. *Cyclobalanopsis* (Supplementary Fig. S4–11; Table S5). First, we performed a polytomy test (Sayyari and Mirarab 2018) to assess whether hard polytomies could be rejected at these uncertain nodes in ASTRAL-III with –t 10. Second, we used approximately unbiased (AU) tests (Shimodaira 2002) to examine whether a tree topology could be significantly rejected by data in IQ-TREE v.2.1.2 (Minh et al. 2020) with 10000 RELL replicates. Third, we quantified the phylogenetic signal of each topology using the pipeline of Shen et al. (2017). Phylogenetic signal here is defined as the difference in log-likelihood scores between two (or three) alternative resolutions (T1 and T2/T3) of a given node (branch or clade) in a tree. In this study, we calculated the difference in the site-wise log-likelihood scores (ΔSLS) and in the gene-wise log-likelihood scores (ΔGLS). Specifically, we estimated the site-wise log-likelihood scores for T1 and T2 (and T3) using the concatenated nuclear matrix and RAxML with –f G. ΔSLS was then calculated by the difference in the site-wise log-likelihood scores of T1 vs. T2 (vs. T3) for every site in the data matrix. ΔGLS was calculated by the sum of ΔSLS of all sites in a given gene.

*Coalescent Simulations to Examine Cytonuclear Discordance*

We applied coalescent simulations to investigate whether the observed deep cytonuclear discordance was likely due to ancient gene flow or incomplete lineage sorting (ILS), following several studies (e.g., Folk et al. 2017; García et al. 2017; Morales-Briones et al. 2018). In this approach, 1000 gene trees were simulated under the coalescent in the package DendroPy (Sukumaran and Holder 2010) with the ASTRAL species tree as the guide tree. The guide tree was pruned to match the taxon set of the chloroplast tree and scaled by a factor of two or four to approximate the branch lengths expected under organellar inheritance (with hermaphrodite and dioecious flowers respectively). We then summarized the frequency at which each clade was observed in the simulated chloroplast trees by mapping the simulated trees against the chloroplast and nuclear species trees using RAxML with -f b. Nuclear branches with high frequencies of alternative topologies (including the empirical chloroplast topology) are consistent with expectations under ILS. However, if the empirical chloroplast topology is observed rarely (or never) in the simulated chloroplast trees (vs. the nuclear species tree), this suggests that the empirical chloroplast topology is a result of gene flow (García et al. 2017; Morales-Briones et al. 2018; Stull et al. 2020).

*Quantifying Gene Tree Discordance Due to ILS, Estimation Error, and Gene Flow*

A high level of gene tree discordance is observed across the whole phylogeny (except several deep nodes) even at the generic and sectional levels (see Supplementary Results). Such conflict may be explained by the presence of ILS, gene tree estimation error, and gene flow (e.g., Maddison 1997; Galtier and Daubin 2008; Knowles et al. 2018; Rose et al. 2021). Newly developed approaches have enabled us to quantify these three causes across the internodes of species trees, and then estimate their relative contributions to gene tree discordance. We used the pipeline of Cai et al. (2021) (https://github.com/lmcai/Coalescent_simulation_and_gene_flow_detection/) to estimate the relative contributions of these sources of conflict to the observed gene tree discordance. Briefly, gene tree discordance is estimated by the gene concordance factor (gCF) using all bootstrap trees from gene trees estimation in IQ-TREE. The level of ILS is represented by the theta value. We first estimated the branch lengths in mutation units (i.e., substitutions per site) in RAxML with the fixed topology of the ASTRAL tree and the concatenated data matrix, and subsequently the theta value at each internode in the species tree was calculated by dividing the branch length in the RAxML tree (mutation unit) by that in the ASTRAL tree (coalescent unit). To quantify the level of gene tree estimation error at every internode, we first used

Seq-Gen v.1.3.4 (Rambaut and Grassly 1997) to simulate gene alignments—the number and length of simulated alignments were equal to the number and median length of those in our empirical dataset—with the ASTRAL tree as the reference tree and the same substitution model parameters of the empirical ML gene trees. We then inferred gene trees from the simulated alignments and applied them to calculate bipartition support values on each node of the ASTRAL species tree in RAxML. Bipartition values here represent the anticipated level of estimation error. The intensity of potential gene flow is represented by the reticulation index, i.e., the percentage of unbalanced triples at each node, which is estimated by three steps: (1) simulating gene trees under the coalescent model for each bootstrap ASTRAL tree (from a second ASTRAL analysis with multi-locus bootstrapping based on empirical gene trees and their bootstrap replicates), (2) counting triplet frequency in empirical and simulated gene trees, and (3) finding significantly unbalanced triplets and mapping them to the species tree. Finally, we formatted these four estimated variables into a matrix of a dependent (i.e., gene tree discordance) and three independent (i.e., ILS, gene tree estimation error, and gene flow) variables across all internodes with each row representing these values for one node. The relative importance of three regressors can be decomposed using a linear regression method implemented by the functions “boot.relimp()” and “booteval.relimp()” in the R package relaimpo (Grömping 2006).

*Divergence Time Estimation*

To generate a dated nuclear tree, we first identified a set of 20 genes showing minimal topological conflict with the species tree as well as more ‘clock-like’ molecular evolution (i.e., low levels of root-to-tip variance in branch lengths) using SortaDate (Smith et al. 2018). A Bayesian dating analysis was then performed in BEAST v.2.6 (Bouckaert et al. 2014) under an uncorrelated lognormal relaxed clock, using a concatenated matrix of the 20 clock-like genes, that included all Fagaceae species with species outside Fagaceae excluded. We employed a Yule speciation process and GTR-GAMMA substitution model and fixed the topology with the concatenated ML tree. Ten well-documented fossils (e.g., Takahashi et al. 2008; Barrón et al. 2017; Denk et al. 2017) were used to calibrate the crown age of Fagaceae, the crown age of *Fagus*, the stem age of *Castanopsis*, the stem age of *Quercus* and five major clades of *Quercus* (see Supplementary Table S6 for detailed prior settings). Monte Carlo Markov chains were run for 600 million generations sampling every 10,000 generations. Convergence was assessed in Tracer v.1.7 (Rambaut et al. 2018) with effective sample size of each parameter more than 200. The maximum clade credibility tree was summarized in TreeAnnotator v.2.6 (Bouckaert et al. 2014) with the first 20% of posterior trees as burn-in, and was used for $D_{\mathrm{FOIL}}$ and biogeography analyses.

Because chloroplast capture events are products of gene flow (also involving the nuclear genome), the timing of chloroplast divergences between oak lineages and other genera should followed the timing of nuclear divergences between them and serve as an independent source of evidence for the timing of hybridization events, since gene flow predicts discordant clades consistently younger than the divergence times of the nuclear species tree estimate, unlike ILS expectations. We specifically focused on the divergence times (1) between the clade of *Castanopsis* + *Castanea* and *Q.* subg. *Cerris*, (2) between *Lithocarpus* and *Q.* subg. *Cerris*, (3) between *Notholithocarpus* and *Q.* subg. *Quercus*, and (4) between *Chrysolepis* and *Q.* subg. *Quercus*. In the chloroplast phylogeny, three castaneoid genera (*Castanea*, *Castanopsis*, and *Lithocarpus*) nest within an Old World clade including the species of Old World oak clade (i.e., subg. *Cerris*), and two castaneoid genera (*Chrysolepis* and *Notholithocarpus*) nest within a New World clade including the species of New World oak clade (i.e., subg. *Quercus*). All five castaneoid genera except for *Chrysolepis* are monophyletic in the chloroplast tree. In light of these findings and inferences of ancient hybridization supported by our and previous studies, we argue that the present plastomes of *Castanea*, *Castanopsis*, and *Lithocarpus* were most likely captured from the Old World oak clade, while the present plastomes of *Chrysolepis* and *Notholithocarpus* were most likely captured from the New World oak clade.

Dating of the chloroplast tree (inferred using the unpartitioned GTR-GAMMA model) was performed using penalized likelihood in treePL (Smith and O'Meara 2012), with cross-validation tests to determine the optimal smoothing parameters for the treePL analyses. Comparable treePL analyses were also performed on the nuclear ML tree to allow comparison between the nuclear and chloroplast divergence times. Given topological differences between the nuclear and chloroplast phylogenies for

Quercoideae, two calibration strategies were used: (1) two fossil calibrations at concordant nodes: crown group (CG) of Fagaceae (max = 89.8 Ma; min = 81 Ma), and CG of genus *Fagus* (min = 47 Ma); and (2) four fossil calibrations at concordant nodes: CG of Fagaceae, CG of genus *Fagus*, stem group (SG) of the East Asian clade in sect. *Cerris* (min = 30 Ma; monophyletic in chloroplast tree), and SG of sect. *Lobatae* (min = 48.7 Ma; monophyletic in chloroplast tree); see Supplementary Table S6 for more details.

*Assessment of Ancient Gene Flow*

*$D_{FOIL}$ analyses*.—$D_{\mathrm{FOIL}}$ tests are a system of applying *D*-statistics (Durand et al. 2011) to a symmetric five-taxon phylogeny (((P1,P2),(P3,P4)),O) (Pease and Hahn 2015). We only tested the four-taxon combinations of the genera *Castanea*, *Castanopsis*, *Lithocarpus*, *Chrysolepis*, *Notholithocarpus*, and *Quercus* (due to the strong cytonuclear discordance observed particularly in Quercoideae), with combinations of taxa from the same genus (or the same section of *Quercus*) excluded (given our focus here on deep ancient introgression). *Trigonobalanus doichangensis* was selected as the outgroup, and the concatenated alignment was used for the sites count. To achieve computational efficiency (Lambert et al. 2019), we reduced the HYB-98RT dataset to 150 taxa by selecting 20 representatives of each genus and each section of *Quercus* (for genera/sections with more than 20 species); we selected species that maximized data quality (i.e., the most loci recovered from assembly) as well as phylogenetic breadth for each genus/section. We also conducted analyses using the full

RNA-2821RT dataset given its more limited taxon sampling. For simplicity, we mainly focused on introgression results involving *Quercus*, with visualization of the results performed using an R script (https://github.com/SheaML/ExDFOIL/).

*Phylogenetic network analyses*.—Three strategies were used to generate a taxonomically highly reduced-representation dataset for species network analyses. One strategy involved selecting two species of each genus of Fagaceae and one outgroup (hereafter "planA"). The second strategy involved reducing our taxon sampling to one species of each non-*Quercus* genus of Fagaceae, one species of each section in *Quercus,* and one outgroup (hereafter "planB"). The third strategy involved selecting two species of each section in *Quercus* and one outgroup (hereafter "planC"). The first two strategies were aimed at identifying possible reticulation between oaks and other genera, while the third one was aimed at identifying possible deep reticulation within oaks. Rooted gene trees were pruned from the HYB-114RT and RNA-4853RT datasets (as these two datasets included the most loci of the Hyb-Seq and transcriptome datasets) to generate each reduced-representation dataset (i.e., planA, planB, and planC) using a custom R script. Only pruned gene trees with > 33% species for each reduced taxon set were used for PhyloNet analyses. For each dataset, we ran five network searches allowing 1 to 6 reticulation events and 10 runs using a maximum pseudo-likelihood approach (command "InferNetwork_MPL"). To infer the optimal number of reticulations (0 to 6, with the strictly bifurcating species tree used to represent 0 reticulations; the likelihood score of this tree was calculated using the command “CalGTProb” in PhyloNet), we then carried out model selection using the bias-corrected Akaike information criterion (Sugiura 1978) and the Bayesian information criterion (Schwarz 1978). The number of parameters was set to the sum of the number of branch lengths and inheritance probabilities (IPs) being estimated. The sample size was set to the number of gene trees for finite sample correction. The reticulation scenario with the lowest information criteria score was selected as the best model.

*Ancestral Niche Estimation*

We collected occurrence records for each sampled species from GBIF, iDigBio, NSII, and the literature. For each species, we removed records that were cultivated and outside the native distribution (detected by the visualization of all records in ArcGIS; native ranges were based on two online databases: World Checklist of Selected Plant Families [http://wcsp.science.kew.org/qsearch.do] and Oaks of the world [http://oaks.of.the.world.free.fr/index.htm]). We also only retained one record per 1 km grid cell (i.e., 30-seconds resolution). This resulted in 187,117 occurrences for subsequent analyses. Fourteen Fagaceae species with fewer than three records after cleaning were excluded from the ancestral niche reconstructions.

We collected 35 environmental layers at 30 seconds resolution that covered aspects of climate (19 temperature and precipitation variables from WorldClim: https://worldclim.org/data/worldclim21.html), soil (seven layers from SoilGrids1km [https://soilgrids.org/]: bulk density, pH, coarse fragment percentage, sand percentage, clay percentage, silt percentage, organic carbon content), topography (three layers: elevation from GTOPO30 [https://lta.cr.usgs.gov/GTOPO30], slope and aspect were calculated from elevation layer), and landcover (six layers from EarthEnv [https://www.earthenv.org/landcover]: percentage cover of needle-leaf trees, evergreen broadleaf trees, deciduous broadleaf trees, mixed trees [including mixed forest and savanna], shrubs, and herbaceous vegetation). We extracted environmental data from 35 layers and cleaned occurrence records for all Fagaceae species using the R package Raster (http://CRAN.R-project.org/package=raster). A pair-wise correlation analyses among environmental variables was run using the R package PKUss (https://github.com/filBe87/PKUss/), and accordingly one from each highly correlated (Pearson's $r \geq 0.7$) variable pair was removed, which resulted in 23 variables. We then chose 12 representative layers relevant to Fagaceae from this set: mean annual temperature, temperature annual range, mean annual precipitation, precipitation seasonality, elevation, slope, bulk density, mean pH, mean organic carbon content, evergreen broadleaf landcover percentage, deciduous broadleaf landcover percentage, and mixed trees landcover percentage.

We created predicted niche occupancy profiles (PNOs, Evans et al. 2009) to represent the distribution of niche space occupancy for all 12 representative layers. A PNO profile is a density histogram that integrates cumulative probabilities of suitability for each value on a single environmental layer (typically derived from an ecological niche model). Here, for each species and layer, we generated a PNO profile based on observed environmental values—that were extracted directly from layer at occurrence records—under a uniform distribution (i.e., the probability of value for each occurrence equals to "1/the number of occurrences").

Ancestral niche estimation was performed in BayesTraits v.2.0 (http://www.evolution.rdg.ac.uk/BayesTraitsV2.html) using the python wrapper 'ambitus' (Folk et al. 2018). Specifically, 100 samples for environmental values were sampled proportionally from a PNO profile for each species and variable, and subsequently each sample was used as "observed value" of each species in a single ancestral niche estimation analysis (leading to 1200 estimation analyses). Each analysis was run on the best ML tree and 1000 RAxML fast bootstrap trees from Hyb-Seq dataset. In each variable, the prior was uniform over the range of minimum and maximum of the observed tip values. For model and reconstruction steps, the MCMC chains were run for 10 million generations sampling every 1000 generations, with the first 50% of posteriors as burn-in. The final results were mapped on the best ML tree.

*Fossil-based Ecological Niche Modeling*

The focus of this analysis is to model the potential distribution of oaks (i.e., *Q.* subg. *Quercus* and *Q.* subg. *Cerris*) and their relatives (i.e., *Notholithocarpus*, *Chrysolepis*, *Lithocarpus*, *Castanea*, and *Castanopsis*) during the Paleogene, the geologic period encompassing the ancient hybridization events inferred here. We first compiled fossil distribution data from the Paleogene (i.e., Paleocene, Eocene, and Oligocene) for oaks and relatives through a comprehensive (to the extent possible) literature survey (e.g., Zhou et al. 1999; Grímsson et al. 2015, 2016; Barrón et al. 2017; Liu et al. 2020; Supplementary Materials) and online resources (Cenozoic Angiosperm Database, Xing et al. 2016; Paleobiology Database, https://paleobiodb.org) (last accessed 21 November 2022). Fossil data were binned by major clade (i.e., *Q.* subg. *Quercus*, *Q.* subg. *Cerris*, *Notholithocarpus*, *Chrysolepis*, *Lithocarpus*, *Castanea*, and *Castanopsis*), based on determinations from the literature involving comparisons with nearest living relatives (obtained from literatures or database Paleoflora, http://www2.geo.uni-bonn.de/Palaeoflora/Palaeoflora_home.htm). Fossil data with ambiguous affinities were excluded from further analysis. When not provided in the literature, modern coordinates of fossils were obtained from Google Earth based on the described fossil localities. The final fossil dataset included 468 occurrences, representing 116 records of *Q.* subg. *Quercus*, 187 records of *Q.* subg. *Cerris*, one records of *Notholithocarpus*, two records of *Chrysolepis*, 17 records of *Lithocarpus*, 70 records of *Castanea*, and 75 records of *Castanopsis* (Supplementary Materials).

The paleo-climate data were derived from HadCM3 (specifically HadCM3LB-M2.1D) from the University of Bristol (Valdes et al. 2017; 2021). The original paleo-climate data at a resolution of 3.75° longitude × 2.5° latitude were downscaled to a high spatial resolution (5' × 5' in each grid) using a bilinear interpolation method in the function “resample” of R package Raster. The boundary conditions of these data were described in detail in Zhao et al. (2022). We selected eight climatic variables representing important vegetation predictors at a resolution of

5' × 5' (~10 × 10 km) and with highly related seasonal variables excluded for PaleoENM analysis: mean annual temperature (°C), warmest month mean surface air temperature (°C), coldest month mean surface air temperature (°C), warmest month–coldest month temperature difference (°C), mean annual precipitation (mm), wettest month precipitation (mm), driest month precipitation (mm), and wettest month–driest month precipitation difference (mm). These paleo-climate models cover seven stages in the Paleogene: Early Paleocene (66.0 Ma), Late Paleocene (60.6 Ma), Early Eocene (55.8 Ma), Middle Eocene (44.5 Ma), Late Eocene (35.9 Ma), Early Oligocene (31.0 Ma), and Late Oligocene (25.6 Ma).

We used the workflow of Meseguer et al. (2015) to conduct the fossil-based ecological niche modeling (PaleoENM). Briefly, modern coordinates of fossils were converted to paleo-coordinates based on the fossil ages using the function “reconstruct” from the R package chronosphere (Kocsis and Raja 2020). We extracted climate values for each fossil paleo-coordinate (only one record per 10 km grid cell was retained for each lineage, each stage, and each variable) from paleo-climate models of corresponding stage using the R package Raster. These climate values were used for the multiple comparisons of niche among oak lineages and closely related genera during the Paleogene using the R package agricolae (de Mendiburu 2009). In addition, given the requirement of adequate fossil occurrences to model past distribution and the global temperature change in the Paleogene, we used two strategies to estimate a past climate tolerance (i.e., climate range in shared grid cells that satisfies the requirement of each climatic variable—within the range of maximum and minimum values of each variable) for each lineage: (1) by grouping fossil record of the Paleocene and Eocene into one time slice (hereafter, PE) to estimate a past climate tolerance; (2) by grouping fossil records of the Paleocene, Eocene, and Oligocene into one time slice (hereafter, PEO) to estimate a single past climate tolerance. Assuming niche conservatism through the Paleogene, the climate tolerance was projected over a suite of paleo-climate models (i.e., Late Paleocene, Early Eocene, Middle Eocene, and Late Eocene layers under the PE scenario; Late Paleocene, Early Eocene, Middle Eocene, Late Eocene, Early Oligocene, and Late Oligocene layers under the PEO scenario). The mahalanobis distance (MD) score was calculated to represent the environmental suitability in each grid cell. The MD score was scaled from 0 to 1; a smaller MD score indicates an area with higher suitability, and vice versa. The PaleoENM analysis was not performed for *Chrysolepis* and *Notholithocarpus*, which included fewer than three fossil occurrences; the past distributions of these two genera were represented directly by a fossil distribution map.

### SUPPLEMENTARY RESULTS

#### *Assembly and Alignments*

We recovered a mean number of 1,733,167 Hyb-Seq reads (SD = 1,964,004) per sample with an average of 68% (SD = 12%) of reads on-target, and the sequencing coverage across all samples ranged from 12× to 5143× (mean = 977×; Supplementary Table S2). After data filtering, the final Hyb-Seq datasets encompassed all genera of Fagaceae and all sections and subclades of oaks recognized by Hipp et al. (2020) (Supplementary Table S1). The concatenated matrices of the two Hyb-Seq data sets (HYB-89MO and HYB-98RT) used for phylogenetic analyses had an aligned length of 96,122 bp–105,564 bp with 59,886–64,772 parsimony-informative sites and 21.78%–25.86% undetermined or gap characters (Supplementary Table S3).

With the exception of sect. *Ponticae*, the transcriptome datasets included all other sections of oaks and all genera of Fagaceae (Supplementary Table S1). The concatenated matrices of the two transcriptome datasets (RNA-977MO and RNA-2821RT) used for phylogenetic analyses had an aligned length of 1,229,405 bp–6,050,182 bp with 904,917–1,704,829 parsimony-informative sites and 52.03%–55.25% undetermined or gap characters (Supplementary Table S3).

The plastome dataset included all sections of oaks and all other genera of Fagaceae (Supplementary Table S1). The concatenated plastome matrix had an aligned length of 135,243 bp with 116,579 parsimony-informative sites and 28.71% undetermined or gap characters.

*Phylogenetic Analyses and Topological Conflicts*

In phylogenies from different nuclear datasets and phylogenetic methods (Fig. 1; Supplementary Fig. S4–S11), areas of lower support include the placements of *Chrysolepis*, which was weakly supported as sister to *Lithocarpus* (bootstrap value [BS] = 30), and *Quercus* sect. *Cyclobalanopsis*, which was placed as sister to *Notholithocarpus* (BS = 25) in the concatenated ML tree of the HYB-98RT dataset (Supplementary Fig. S7). The monophyly of *Quercus* received varied support: maximum or moderate support (BS = 100; local posterior probabilities [LPP] = 0.71–1.0) in trees of transcriptome datasets (Supplementary Fig. S8–S11) and moderate or weak support (BS = 48; LPP = 0.49–0.57) in trees of Hyb-Seq datasets (Supplementary Fig. S4–S7). Furthermore, we found a high level of gene tree discordance within the clade formed by *Lithocarpus*, *Chrysolepis*, *Notholithocarpus*, and *Quercus* (Supplementary Fig. S17). No dominant alternative topologies were detected, with the exception of *Chrysolepis*, which had a dominant alternative topology (sister to *Lithocarpus*) in the RNA-2821RT dataset (Supplementary Fig. S17b). The phyparts results from the full taxon set are shown in Supplementary Figure S18–S19. The position of *Chrysolepis* (i.e., sister to *Notholithocarpus* + oaks) received weak QS support (0.26/0.24/0.82) with a skewed frequency in discordant topologies in the HYB-98RT dataset, while this topology of *Chrysolepis* received counter-support (-0.18/0.27/0.95) in the RNA-2821RT dataset, indicating a majority of quartets supported one of the discordant topologies of *Chrysolepis* (other than the topology showing it sister to *Notholithocarpus* + oaks). The monophyly of *Quercus* received low QS support in both the HYB-98RT (0.27/0.67/0.86) and RNA-2821RT (0.17/0.57/0.95) datasets; the monophyly of *Q.* subg. *Cerris* (i.e., the placement of sect. *Cyclobalanopsis*) had weak QS support (0.19/0.27/0.83) with a skewed frequency in discordant topologies in the HYB-98RT dataset, while moderate QS support was present (0.43/0.18/0.97) in the RNA-2821RT dataset (Supplementary Fig. S17).

In comparison with nuclear trees, the chloroplast ML trees revealed starkly different relationships among genera in Quercoideae and sections in *Quercus* (Fig. 2a; Supplementary Fig. S20-S21). The topologies of partitioned and unpartitioned chloroplast ML trees were more or less identical to each other with regard to relationships among major lineages (Supplementary Fig. S27–S28). Chloroplast trees did not support the monophyly of *Chrysolepis*, *Quercus*, sect. *Cyclobalanopsis*, sect. *Ilex*, sect. *Protobalanus*, sect. *Ponticae*, sect. *Virentes*, or sect. *Quercus* (Supplementary Fig. S27–S28), most of which were moderately to strongly supported as monophyletic in nuclear species trees (Supplementary Fig. S4–S11). Furthermore, the clade *Castanea* + *Castanopsis* and *Lithocarpus* were recovered as sister (BS = 66), with these together nested in *Q.* subg. *Cerris*. All of these taxa (with the exception of several *Castanea* species) are presently restricted to the Old World. The small genus *Notholithocarpus* and *Chrysolepis*, both from western North America, were recovered as sisters, with these together as sister (BS = 72–73) to a North American oak clade composed of sect. *Protobalanus* and sect. *Quercus* s.l. (sect. *Ponticae* and sect. *Virentes* included, the Eurasia Roburoid lineage and *Q. pontica* excluded). The north temperature disjunct oak clade (sect. *Ponticae*) includes only two species: the Eurasian *Q. pontica* clustered with the Roburoid lineage, and the western North American *Q. sadleriana* clustered with a lineage of sect. *Quercus* (mainly consisting of western North American Dumosae species). This cytonuclear discordance was conspicuously strong even though all nodes below 50% BS (or 0.5 LPP) were collapsed (Fig. 2a; Supplementary Fig. S20-S21). The patterns of cytonuclear discordance above show that chloroplast relationships largely track geography (rather than evolutionary history per se), perhaps due to extensive gene flow.

*Phylogenetic Signal of Alternative Topologies for Five Uncertain Nodes*

The polytomy tests across datasets were generally able to reject the null hypothesis that each of these five uncertain nodes should be replaced by a polytomy ($P < 0.05$); the two exceptions were *Chrysolepis* and sect. *Cyclobalanopsis* in the HYB-98RT dataset (Supplementary Table S5). The AU tests showed that T2 or T3 (i.e., chloroplast topology, the ML topology of the HYB-98RT dataset, or topologies from previous studies) was significantly worse ($P < 0.05$) than T1 (i.e., nuclear topology recovered by all ASTRAL and most concatenated ML trees) for five uncertain nodes in the RNA-2821RT dataset; the same results were shown in three of five uncertain nodes on the basis of the HYB-98RT dataset while T2 of *Chrysolepis* (i.e., sister to *Lithocarpus*) could not be rejected ($P > 0.05$) and T1 of sect. *Cyclobalanopsis* (i.e., sister to sect. *Cerris* + sect. *Ilex*) was rejected instead ($P < 0.05$; Supplementary Table S5).

Examination of the distribution of phylogenetic signal (i.e., ΔGLS and ΔSLS) in alternative tree topologies of the five uncertain nodes showed that the proportions of genes and sites favoring T1 were usually greater than those favoring T2 or T3, but a large amount of conflicting signal was observed across all uncertain nodes and datasets (Supplementary Fig. S12–S13). For the position of *Castanea* + *Castanopsis*, T1 (i.e., sister to the rest of Quercoideae except *Trigonobalanus)* was supported by 78% of genes and 84% of sites in the HYB-98RT data matrix (Supplementary Fig. S12a) and 85% of genes and 72% of sites in the RNA-2821RT data matrix (Supplementary Fig. S13a), while the rest of the genes and sites supported T2 (i.e., nested in *Q.* subg. *Cerris*). For the position of *Lithocarpus*, T1 (i.e., *Lithocarpus* sister to the clade *Chrysolepis* + *Notholithocarpus* + *Quercus*) was supported by 76% of genes and 91% of sites in the HYB-98RT data matrix (Supplementary Fig. S12b) and 70% of genes and 74% of sites in the RNA-2821RT data matrix (Supplementary Fig. S13b), while the rest of the genes and sites supported T2 (i.e., *Lithocarpus* nested in *Q.* subg. *Cerris*). For the position of *Chrysolepis*, T2 (i.e., sister to *Lithocarpus*) had the highest proportion of favoring genes (50%) and sites (65%) in the HYB-98RT data matrix, followed by T1 (i.e., sister to the clade *Notholithocarpus* + *Quercus*, supported by 22% of genes and 26% of sites) and T3 (i.e., nested in *Q.* subg. *Quercus*, supported by 28% of genes and 9% of sites; Supplementary Fig. S12c); in the RNA-2821RT data matrix, the two nuclear (i.e., T1 and T2) and chloroplast (i.e., T3) topologies were supported by similar proportions of genes (30%, 39%, and 31%, respectively) and sites (39%, 36%, and 25%, respectively; Supplementary Fig. S13c), indicating that ILS might be the cause of the two discordant nuclear topologies and ancient introgression the cause of the chloroplast topology. For the position of *Notholithocarpus*, the proportions of genes (51% and 48% in the HYB-98RT and RNA2821RT data matrices, respectively) and sites (65% and 61% in the HYB-98RT and RNA2821RT data matrices, respectively) in favor of T1 (i.e., sister to *Quercus*) were generally greater than those favoring T2 (i.e., nested in *Q.* subg. *Quercus*; Supplementary Fig. S12d–S13d). For the position of sect. *Cyclobalanopsis*, T1 (i.e., sister to the clade of sect. *Cerris* + sect. *Ilex*), T2 (sister to the rest of oaks), and T3 (i.e., sister to *Notholithocarpus*) were supported by 46%, 25%, and 29% of genes and 12%, 27%, and 60% sites, respectively, in the HYB-98RT data matrix (Supplementary Fig. S12e), which might explain the recovery of T1 in the ASTRAL tree and T3 in the concatenated ML tree; in the RNA-2821RT data matrix, T1 had the highest proportions of supporting genes (45%) and sites (79%), followed by T3 (33% of genes and 20% of sites) and T2 (21% genes and 10% sites; Supplementary Fig. S13e).

Collectively, the proportions of nuclear genes and sites supporting the chloroplast topologies were non-negligible, ranging from 15% to 52% of genes and 9% to 39% of sites across four uncertain nodes and two data matrices (Supplementary Fig. S12–S13). This suggests that ancient introgression is likely responsible for much of the conflict observed in these conflicting relationships. It is noteworthy that a handful of genes (one in HYB-98RT dataset; 104 [104/2821 ≈ 3.7%] in RNA-2821RT dataset) have an extensive amount of signal (ΔGLS > 50). The overall pattern of a handful of genes having strong phylogenetic signal (i.e., ΔGLS > 50) has been seen before (e.g.,

Shen et al. 2017; Zhou et al. 2022), suggesting that phylogenetic signal outlier genes are possibly common in phylogenomic data. After careful checking, we speculate that the reason for the strong signal from those genes may be complex and multifarious, such as misidentified orthology, sequencing and gene assembling error, longer gene alignment, or simply differing evolutionary regimes among gene lineages. The topologies for the five uncertain deep nodes did not change when a single outlier gene with strong signal (i.e., maximum ΔGLS) or a set of outlier genes of strong signal (defined as in Shen et al. 2017) were excluded from the full dataset (data not shown). In summary, phylogenetic signal tests show that genes with strong did not affect the relevant conclusions.

*Simulating Chloroplast Trees under the Coalescent*

The organellar gene tree distribution simulated from the scaled ASTRAL tree under the coalescent showed that much of the nuclear backbone topology was within ILS predictions and had moderate to high clade probabilities (with some notable exceptions for sect. *Lobatae* and sect. *Quercus*); little of the topology of the chloroplast was within ILS predictions (clade probabilities mostly ~0; Fig. 2a; Supplementary Fig. S20–S23). Focusing on the deep conflicting topologies between the nucleus and chloroplast at the clades corresponding to generic and sectional scales, clade probabilities of simulated trees (with the guide tree scaled by a factor of four) evaluated on the chloroplast tree were less than 4% and 3% in the HYB-98RT and RNA-2821RT datasets, respectively (see red node labels in Fig. 2a and Supplementary

Fig. S20–S21), suggesting that the deep topologies of the chloroplast tree (that conflict with the nuclear species tree) are not within ILS expectations (thus gene flow is a better explanation for deep cytonuclear discordance in Quercoideae). A similar pattern was also found in an organellar gene tree distribution simulated under the ASTRAL tree scaled by two (Supplementary Fig. S22–S23); the only difference was that the overall clade probabilities were slightly higher or lower than those under the scale factor of four.

*Co-quantification of ILS, Gene Tree Estimation Error, and Gene Flow*

Relative importance decomposition analysis demonstrated that the linear models (on the basis of ILS, gene tree estimation error, and gene flow) explained 10.45% and 28.36% of the total gene tree variation in the HYB-98RT and RNA-2821RT datasets, respectively, across all internodes of Fagaceae, while the models explained 10.89% and 52.28% of gene tree variation in the HYB-98RT and RNA-2821RT datasets, respectively, across all internodes of *Quercus* (see $R^2$ in Supplementary Fig. S26). ILS and gene flow were the two main factors explaining gene tree conflict across the two datasets and taxonomic levels. Specifically, the relative contributions of ILS, gene flow, and estimation error explaining gene tree variation across Fagaceae were 3.25%, 6.58%, and 0.62% in the HYB-98RT dataset (i.e., 31.11%, 62.95%, and 5.94% of $R^2$ [=10.45%] respectively; Supplementary Fig. S26a) and 19.64%, 5.15%, and 3.57% in the RNA-2821RT dataset (Supplementary Fig. S26c). Across *Quercus*, ILS was the dominant factor, explaining 7.79% of the total gene tree variation in the HYB-98RT dataset (Supplementary Fig. S26b) and 36.31% in RNA-2821RT dataset (Supplementary Fig. S26d); the second most important factor was gene flow, which explained 2.19% of gene tree variation in the HYB-98RT dataset and 13.02% in the RNA-2821RT dataset; gene tree estimation error explained the least variation (0.91% in the HYB-98RT dataset and 2.95% in the RNA-2821RT dataset).

*Divergence time estimation*

The chloroplast and nuclear dating analyses using treePL showed that the divergence times among the non-*Quercus* genera (in Quercoideae) and *Quercus* were younger in the chloroplast tree vs. the nuclear tree in both calibration scenarios (i.e., with two and four calibrations; Fig. 2b). The MCC tree from the BEAST analysis including nine calibrations showed older divergence times than both the nuclear and chloroplast treePL analyses. The inferred divergence times of four key nodes from the dated chloroplast tree (four calibrations in treePL), the dated nuclear tree (four calibrations in treePL), and the nuclear MCC tree (nine calibrations in BEAST), respectively, are also follows: the split of *Castanopsis* + *Castanea* and *Q.* subg. *Cerris* (47.0573, 54.1271, 64.2473 Ma); the split of *Lithocarpus* and *Q.* subg. *Cerris* (47.0573, 51.9321, 59.7094 Ma), the split of *Chrysolepis* and *Q.* subg. *Quercus* (46.9273, 51.1016, 57.1719 Ma), as well as the split of *Notholithocarpus* and *Q.* subg. *Quercus* (46.9273, 49.9004, 55.3428 Ma).

*Five-Taxon Analyses in Quercoideae*

We found significant signals of introgression (2.49% in the $D_{\text{FOIL}}$ tests, 132,435/5,316,699 for the HYB-98RT dataset; 3.58% in $D_{\text{FOIL}}$ tests, 8,058/224,998 for the RNA-2821RT dataset). Most positive tests were for signatures of ancestral introgression (129,770/132,435 in dataset HYB-98RT and 7,436/8,058 in dataset RNA-2821RT), while inter-group signatures (i.e., between two terminal taxa from different subgroups) were observed less frequently.

Overall, signatures of ancient introgression were inferred between lineages of *Quercus* and other genera in Quercoideae across the two datasets (except for *Notholithocarpus* and *Trigonobalanus*, possibly because only one individual sampled for *Notholithocarpus,* which could not satisfy the requirement of a symmetric five-taxon phylogeny for $D_{\text{FOIL}}$ tests, and because *Trigonobalanus* was used as the outgroup); however, signatures within *Quercus* were the most frequent overall (Supplementary Fig. S24–S25). At the generic level in the HYB-98RT dataset, introgression signatures involving *Lithocarpus* and *Quercus* lineages (e.g., *Quercus* / *Quercus* <–> *Lithocarpus*; and *Lithocarpus* / *Lithocarpus* <–> *Quercus*) were detected the most in positive results for *Lithocarpus* while a similar pattern was observed in *Chrysolepis* (the strongest introgression signature was *Chrysolepis* / *Chrysolepis* <–> *Quercus*; Supplementary Fig. S24). In *Castanopsis*, the introgression signatures including *Castanopsis* and *Quercus* lineages (e.g., *Castanopsis* / *Castanopsis* <–> *Quercus*) were found in similar proportion (even lower proportion on some nodes), with signatures involving *Castanopsis* and non-*Quercus* lineages (e.g., *Castanopsis* / *Castanopsis* <–> *Lithocarpus* in Supplementary Table S7; "others" in Supplementary Fig. S24), while a similar pattern was observed in *Castanea* (with the two strongest introgression signatures involving *Castanea* / *Castanea* <–> *Quercus* and *Castanea* / *Castanea* <–> *Lithocarpus*; Supplementary Table S7; Supplementary Fig. S24). The pattern of introgression at the generic level in the RNA-2821RT dataset was generally similar to that of the HYB-98RT dataset, except that introgression between *Castanopsis* (or *Castanea*) and non-*Quercus* lineages was recovered less often, possibly due to the relatively low taxon sampling of non-*Quercus* species in the transcriptome dataset (Supplementary Table S7; Supplementary Fig. S25).

Furthermore, a summary of all positive $D_{\mathrm{FOIL}}$ tests (categorized by the two subgenera of *Quercus* and closely related genera) showed a geographic pattern of introgression signatures, in which the number of significant $D_{\mathrm{FOIL}}$ tests between non-*Quercus* genera and subgenera of *Quercus* from the same continent was greater than that between non-*Quercus* genera and subgenera of oaks distributed on different continents (Supplementary Table S8), consistent with the likely hypothesis that species with overlapping distributions were most likely to hybridize. For instance, the number of positive tests for introgression signatures between the Old World *Lithocarpus* and *Q.* subg. *Cerris* were more than those between *Lithocarpus* and the New World clade *Q.* subg. *Quercus* (e.g., 7,147 vs. 3,851 for signature "*Lithocarpus* / *Lithocarpus* <–> X"; Supplementary Table S8). The number of positive tests for introgression signatures between the New World *Chrysolepis* and *Q.* subg. *Cerris* were less than those between *Chrysolespis* and *Q.* subg. *Quercus* (e.g., 201 vs. 730 for signature "*Chrysolepis* / *Chrysolepis* <–> X"). The above geographic pattern of introgression revealed by the reduced taxon set of HYB-98RT (with similar species number of the two subgenera of genus *Quercus*, 49 vs. 51) was relatively obscure in RNA-2821RT, most likely due to the biased taxon sampling of the two subgenera of *Quercus* (14 vs. 41; Supplementary Table S10).

*Phylogenetic Network Inference*

Model selection suggested that any network was a better model than the strictly bifurcating species tree across the two datasets (Supplementary Table S11–S12). The optimal networks of the three representation strategies of the two dataset types often presented incongruent scenarios, but all supported complex reticulations in the early evolutionary history of subfamily Quercoideae (Supplementary Fig. S16). Regarding the reticulation between oaks and their relatives, optimal networks recovered gene flow from oaks to the *Trigonobalanus excelsa* lineage, with an inheritance probability (IP) of 0.205 (Supplementary Fig. S16a), and from an ancestral lineage of *Quercus* + *Notholithocarpus* to the clade of sect. *Cerris* + sect. *Ilex* (IP = 0.353; Supplementary Fig. S16b) and to *Quercus* with a low IP of 0.001 (Supplementary Fig. S16d). Three reticulation events with parents showing relatively equivalent inheritance probabilities might represent instances of hybrid speciation, resulting in the following reticulate lineages: *Trigonobalanus* (0.465 vs. 0.535; Supplementary Fig. S16a), the lineage comprising *Lithocarpus*, *Notholithocarpus*, and oaks (0.442 vs. 0.558; Supplementary Fig. S16a), and the lineage comprising *Notholithocarpus* and oaks (0.455 vs. 545 in

Supplementary Fig. S16d; 0.434 vs. 0.566 in Supplementary Fig. S16e). The best species networks did not support gene flow between *Lithocarpus* (*Chrysolepis*, *Castanopsis*, or *Castanea*) and oaks, although $D_{\mathrm{FOIL}}$ tests suggested significant ancient introgression between them. Within oaks, prevalent gene flow was inferred among major lineages. For example, gene flow was detected from the ancestral lineage of the clade sect. *Virentes* + sect. *Quercus* to the *Q. palmeri* lineage of sect. *Protobalanus* (IP = 0.214; Supplementary Fig. S16c), and from the ancestral lineage of *Q.* subg. *Quercus* to the clade sect. *Cerris* + sect. *Ilex* (IP = 0.319; Supplementary Fig. S16f). The Eurasian species *Q. pontica* in sect. *Ponticae* might represent the product of hybrid speciation, involving the Eurasian Roburoid lineage (represented by *Q. robur* here) in sect. *Quercus* and an ancestral lineage of sect. *Ponticae* (IPs = 0.417 vs. 0.583; Supplementary Fig. S16c).

*Biogeographic Inference*

The DEC model (AIC = 1205; AIC weight = 1.00) was selected as the best fit biogeographic model (Supplementary Table S13).

*Ancestral Niche Reconstruction*

Analysis of ancestral niche reconstruction based on extant species showed that the environmental spaces of the most recent common ancestor (MRCA) of *Castanea* + *Castanopsis*, the MRCA of *Lithocarpus*, and the MRCA of *Q.* subg. *Cerris* overlapped for all representative variables, which included the aspects of climate, topography, soil, and landcover (Fig. 4a; Supplementary Fig. S32). Overlapped niche spaces were also inferred between the MRCA of *Chrysolepis* and the MRCA of *Q.* subg. *Quercus*, as well as between *Notholithocarpus* and the MRCA of *Q.* subg. *Quercus* (Fig. 4a; Supplementary Table S9). The reconstructed temperature and precipitation values of these MRCAs indicated the ancestors of these clades lived in a warm and moist environment (Fig. 4a). Furthermore, ancestral niche reconstruction of all variables showed niche divergence between subclades of some oak sections yet niche conservatism within subclade, such as sect. *Lobatea*, sect. *Quercus*, sect. *Cyclobalanopsis* (except for the variable of deciduous broadleaf landcover showing niche conservatism between subclades), and sect. *Ilex* (except for two variables of mean annual temperature and deciduous broadleaf landcover showing niche conservatism between subclades) (Supplementary Fig. S32). These four sections together comprise roughly 95% of oak species (Denk et al. 2017), and this finding suggests the essential role of niche divergence in the rapid diversification of *Quercus*.

Kearse M., Moir R., Wilson A., Stones-Havas S., Cheung M., Sturrock S., Buxton S., Cooper A., Markowitz S., Duran C., Thierer T., Ashton B., Meintjes P., Drummond

A. 2012. Geneious Basic: an integrated and extendable desktop software platform for the organization and analysis of sequence data. Bioinformatics 28:1647–1649.

Knowles L.L., Huang H., Sukumaran J., Smith S.A. 2018. A matter of phylogenetic scale: distinguishing incomplete lineage sorting from lateral gene transfer as the cause of gene tree discord in recent versus deep diversification histories. Am. J. Bot. 105:376–384.

Kocsis A.T., Raja N.B. 2020. chronosphere: Earth system history variables. Available from https://cran.r-project.org/web/packages/chronosphere/index.html.

Lambert S.M., Streicher J.W., Fisher-Reid M.C., Mendez de la Cruz F.R., Martinez-Mendez N., Garcia-Vazquez U.O., Nieto Montes de Oca A., Wiens J.J. 2019. Inferring introgression using RADseq and $D_{\mathrm{FOIL}}$: power and pitfalls revealed in a case study of spiny lizards (*Sceloporus*). Mol. Ecol. Resour. 19:818–837.

Langmead B., Salzberg S.L. 2012. Fast gapped-read alignment with Bowtie 2. Nat. Methods 9:357–359.

Li H. 2011. A statistical framework for SNP calling, mutation discovery, association mapping and population genetical parameter estimation from sequencing data. Bioinformatics 27:2987–2993.

Li H. 2013. Aligning sequence reads, clone sequences and assembly contigs with BWA-MEM. arXiv:1303.3997v1302 [q-bio.GN].

Li H., Handsaker B., Wysoker A., Fennell T., Ruan J., Homer N., Marth G., Abecasis G., Durbin R., Subgroup G.P.D.P. 2009. The sequence alignment/map format and SAMtools. Bioinformatics 25:2078–2079.

Liu X.Y., Song H.Z., Jin J.H. 2020. Diversity of Fagaceae on Hainan island of south China during the middle Eocene: Implications for phytogeography and paleoecology. Front. Ecol. Evol. 8: 255.

Maddison W.P. 1997. Gene trees in species trees. Syst. Biol. 46:523–536.

Manos P.S., Cannon C.H., Oh S.H. 2008. Phylogenetic relationships and taxonomic status of the paleoendemic Fagaceae of western North America: recognition of a new genus, *Notholithocarpus*. Madroño 55:181–190.

Manos P.S., Doyle J.J., Nixon K.C. 1999. Phylogeny, biogeography, and processes of molecular differentiation in *Quercus* Subgenus *Quercus* (Fagaceae). Mol. Phylogenet. Evol. 12:333–349.

Meseguer A.S., Lobo J.M., Ree R., Beerling D.J., Sanmartín I. 2015. Integrating fossils, phylogenies, and niche models into biogeography to reveal ancient evolutionary history: The case of *Hypericum* (Hypericaceae). Syst. Biol. 64:215–232.

Minh B.Q., Schmidt H.A., Chernomor O., Schrempf D., Woodhams M.D., von Haeseler A., Lanfear R. 2020. IQ-TREE 2: new models and efficient methods for phylogenetic inference in the genomic era. Mol. Biol. Evol. 37:1530–1534.

Morales-Briones D.F., Gehrke B., Huang C.H., Liston A., Ma H., Marx H.E., Tank D.C., Yang Y. 2021. Analysis of paralogs in target enrichment data pinpoints multiple ancient polyploidy events in *Alchemilla* s.l. (Rosaceae). Syst. Biol. 71:190–207.

Morales-Briones D.F., Liston A., Tank D.C. 2018. Phylogenomic analyses reveal a deep history of hybridization and polyploidy in the Neotropical genus *Lachemilla* (Rosaceae). New Phytol. 218:1668–1684.

Pease J.B., Brown J.W., Walker J.F., Hinchliff C.E., Smith S.A. 2018. Quartet Sampling distinguishes lack of support from conflicting support in the green plant tree of life. Am. J. Bot. 105:385–403.

Pease J.B., Hahn M.W. 2015. Detection and polarization of introgression in a five-taxon phylogeny. Syst. Biol. 64:651–662.

Rambaut A., Drummond A.J., Xie D., Baele G., Suchard M.A. 2018. Posterior summarization in Bayesian phylogenetics using Tracer 1.7. Syst. Biol. 67:901–904.

Rambaut A., Grassly N.C. 1997. Seq-Gen: an application for the Monte Carlo simulation of DNA sequence evolution along phylogenetic trees. Bioinformatics 13:235–238.

Ramos A.M., Usie A., Barbosa P., Barros P.M., Capote T., Chaves I., Simoes F., Abreu I., Carrasquinho I., Faro C., Guimaraes J.B., Mendonca D., Nobrega F., Rodrigues L., Saibo N.J.M., Varela M.C., Egas C., Matos J., Miguel C.M., Oliveira M.M., Ricardo C.P., Goncalves S. 2018. The draft genome sequence of cork oak. Sci. Data 5:180069.

Rose J.P., Toledo C.A.P., Lemmon E.M., Lemmon A.R., Sytsma K.J. 2021. Out of sight, out of mind: widespread nuclear and plastid-nuclear discordance in the flowering plant genus *Polemonium* (Polemoniaceae) suggests widespread historical gene flow despite limited nuclear signal. Syst. Biol. 70:162–180.

Sayyari E., Mirarab S. 2018. Testing for polytomies in phylogenetic species trees using quartet frequencies. Genes (Basel) 9:132.

Schmid-Siegert E., Sarkar N., Iseli C., Calderon S., Gouhier-Darimont C., Chrast J., Cattaneo P., Schütz F., Farinelli L., Pagni M., Schneider M., Voumard J., Jaboyedoff M., Fankhauser C., Hardtke C.S., Keller L., Pannell J.R., Reymond A., Robinson-Rechavi M., Xenarios I., Reymond P. 2017. Low number of fixed somatic mutations in a long-lived oak tree. Nat. Plants 3:926–929.

Schwarz G. 1978. Estimating the dimension of a model. Ann. Stat. 6:461–464.

Shen X.X., Hittinger C.T., Rokas A. 2017. Contentious relationships in phylogenomic studies can be driven by a handful of genes. Nat. Ecol. Evol. 1:126.

Shimodaira H. 2002. An approximately unbiased test of phylogenetic tree selection. Syst. Biol. 51:492–508.

Simeone M.C., Grimm G.W., Papini A., Vessella F., Cardoni S., Tordoni E., Piredda R., Franc A., Denk T. 2016. Plastome data reveal multiple geographic origins of *Quercus* Group *Ilex*. PeerJ 4:e1897.

Slater G.S., Birney E. 2005. Automated generation of heuristics for biological sequence comparison. BMC Bioinform. 6:31.

Smith-Unna R., Boursnell C., Patro R., Hibberd J.M., Kelly S. 2016. TransRate: reference-free quality assessment of *de-novo* transcriptome assemblies. Genome Res. 26:1134–1144.

Smith S.A., Brown J.W., Walker J.F. 2018. So many genes, so little time: a practical approach to divergence-time estimation in the genomic era. PLoS One 13:e0197433.

Smith S.A., Moore M.J., Brown J.W., Yang Y. 2015. Analysis of phylogenomic datasets reveals conflict, concordance, and gene duplications with examples from animals and plants. BMC Bioinform. 15:150.

Smith S.A., O'Meara B.C. 2012. treePL: divergence time estimation using penalized likelihood for large phylogenies. Bioinformatics 28:2689–2690.

Song L., Florea L. 2015. Rcorrector: efficient and accurate error correction for Illumina RNAseq reads. GigaScience 4:48.

Sork V.L., Fitz-Gibbon S.T., Puiu D., Crepeau M., Gugger P.F., Sherman R., Stevens K., Langley C.H., Pellegrini M., Salzberg S.L. 2016. First draft assembly and annotation of the genome of a California endemic oak *Quercus lobata* Née (Fagaceae). G3: Genes, Genomes, Genet. 6:3485–3495.

Stamatakis A. 2014. RAxML Version 8: A tool for phylogenetic analysis and post-analysis of large phylogenies. Bioinformatics 30:1312–1313.

Stull G.W., Soltis P.S., Soltis D.E., Gitzendanner M.A., Smith S.A. 2020. Nuclear phylogenomic analyses of asterids conflict with plastome trees and support novel relationships among major lineages. Am. J. Bot. 107:790–805.

Sugiura N. 1978. Further analysts of the data by akaike's information criterion and the finite corrections. Commun. Stat. Theory Methods 7:13–26.

Sukumaran J., Holder M.T. 2010. DendroPy: a Python library for phylogenetic computing. Bioinformatics 26:1569–1571.

Takahashi M., Friis E.M., Herendeen P.S., Crane P.R. 2008. Fossil flowers of Fagales from the Kamikitaba locality (early Coniacian; late Cretaceous) of northeastern

Japan. Int. J. Plant Sci. 169:899–907.

Valdes P.J., Armstrong E., Badger M.P.S., Bradshaw C.D., Bragg F., Crucifix M., Davies-Barnard T., Day J.J., Farnsworth A., Gordon C., Hopcroft P.O., Kennedy A.T., Lord N.S., Lunt D.J., Marzocchi A., Parry L.M., Pope V., Roberts W.H.G., Stone E.J., Tourte G.J.L., Williams J.H.T. 2017. The BRIDGE HadCM3 family of climate models: HadCM3@Bristol v1.0. Geosci. Model Dev. 10:3715–3743.

Valdes P.J., Scotese C.R., Lunt D.J. 2021. Deep ocean temperatures through time. Clim. Past 17:1483–1506.

van Dongen S.M. 2000. Graph clustering by flow simulation. PhD diss., Ultrecht University.

Xing Y.W., Gandolfo M.A., Onstein R.E., Cantrill D.J., Jacobs B.F., Jordan G.J., Lee D.E., Popova S., Srivastava R., Su T., Vikulin S.V., Yabe A., Linder H.P. 2016. Testing the biases in the rich Cenozoic Angiosperm macrofossil record. Int. J. Plant Sci. 177:371–388.

Yang Y., Smith S.A. 2014. Orthology inference in nonmodel organisms using transcriptomes and low-coverage genomes: improving accuracy and matrix occupancy for phylogenomics. Mol. Biol. Evol. 31:3081–3092.

Yang Y.Y., Qu X.J., Zhang R., Stull G.W., Yi T.S. 2021. Plastid phylogenomic analyses of Fagales reveal signatures of conflict and ancient chloroplast capture. Mol. Phylogenet. Evol. 163:107232.

Zhao J.G., Li S.F., Farnsworth A., Valdes P.J., Reichgelt T., Chen L.L., Zhou Z.K., Su T. 2022. The Paleogene to Neogene climate evolution and driving factors on the

Qinghai-Tibetan Plateau. Sci. China Earth Sci. 65:1339–1352.

Zhou B.F., Yuan S., Crowl A.A., Liang Y.Y., Shi Y., Chen X.Y., An Q.Q., Kang M., Manos P.S., Wang, B.S. 2022. Phylogenomic analyses highlight innovation and introgression in the continental radiations of Fagaceae across the Northern Hemisphere. Nat. Commun. 13:1320.

Zhou Z.K. 1999. Fossils of the Fagaceae and their implications in systematics and biogeography. Acta Phytotax. Sin. 37: 369–385.
